# Genomics-Driven Discovery of Myxopyromides, a Class of Polyketidic Amides Associated with Predation of *Myxococcus* sp. SDU36

**DOI:** 10.1101/2023.04.01.535179

**Authors:** Luo Niu, Wei-Feng Hu, Wen-Juan Zhang, Zhao-Min Lin, Jing-Jing Wang, Le-Le Zhu, Jia-Qi Hu, Chao-Yi Wang, Rui-Juan Li, Yue-Zhong Li, Changsheng Wu

**Affiliations:** State Key Laboratory of Microbial Technology, Institute of Microbial Technology, Shandong University, 266237 Qingdao, P.R. China; Institute of Medical Science, The Second Hospital, Cheeloo College of Medicine, Shandong University, Jinan 250033, P.R. China

## Abstract

Myxobacteria are renowned for their genuine prowess to deliver multifarious bioactive natural products. Genome mining of *Myxococcus* sp. SDU36 identified a hybrid PKS-NRPS BGC (*mpd*) that presumably specified previously uncharacterized molecules. The expression of *mpd* was activated by exchanging the innate promoter of core PKS-NRPS genes with our recently characterized strong constitutive promoter BBa_J23104. Comparative metabolic profiling allowed facile isolation of the elicited compounds **1**–**5** designated myxopyromides A–E, a group of structurally related polyketidic amides. Especially, myxopyromides D was appended with an uncommon pyrrolinone warhead at the carboxylic terminus, whereas myxopyromide C was instead decorated with a rare structural unit of aminobutanone. Although myxopyromides basically follow a textbook modular PKS-NRPS biosynthetic trajectory, an anteriorly unappreciated flexible chain release strategy is adopted to enrich the chemical repertoire of *mpd* BGC. Interestingly, myxopyromides were associated with the predation of *Myxococcus* sp. SDU36.

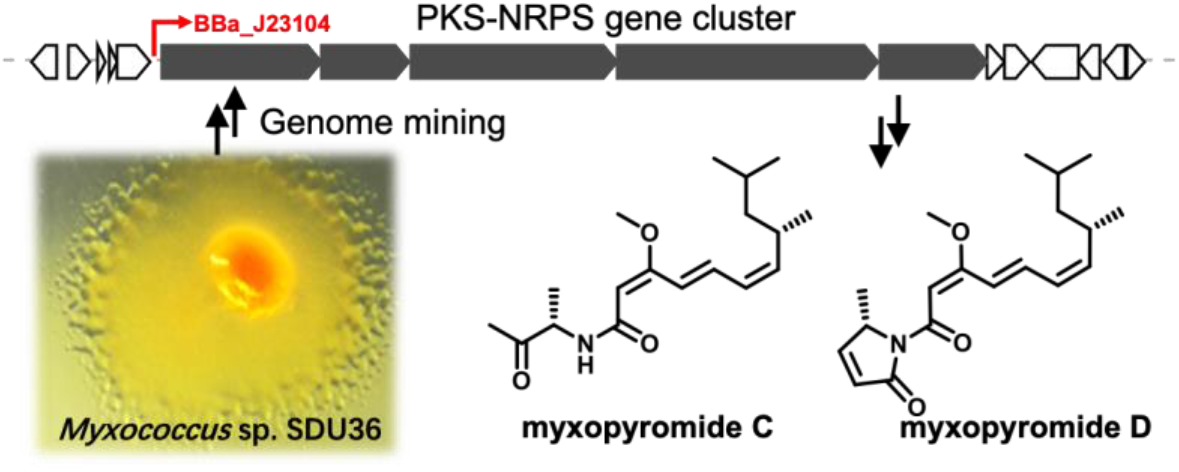

---

Myxobacteria represent an extraordinary member of Gram-negative *δ*-proteobacteria due to their ability to swarm, prey, and form fruiting bodies.^1^ This phylum of microorganisms has been regarded as a vast reservoir of bioactive natural products (NPs).^2^ However, compared with other “gifted” taxa including filamentous actinomycetes, *Bacillus, Pseudomonas*,^3–5^ the exploitation of myxobacteria is largely plagued by the difficulty in the resource availability. Given this, appropriate discovery pipelines are of paramount importance to maximally exploit the existing myxobacterial collections, particularly in terms of awakening the tremendous amount of untapped biosynthetic gene clusters (BGCs). Among others, promoter engineering is commonly deemed as an enticing strategy to activate the silent BGCs because the intricacy in the operon regulation could be bypassed through replacing the native promotor with an exotic promoter.^6^ In light of this, we recently characterized a panel of constitutive promotors workable in myxobacteria, greatly strengthening the synthetic biology toolkit available for the engineering of myxobacterial BGCs.^7^

The secondary metabolites discovered from myxobacteria oftentimes exhibit sheer chemical complexity and enormous pharmaceutical potential,^8^ among which, the majority is synthesized by hybrid BGCs with a mash-up of type I modular polyketide synthases (PKSs) and non-ribosomal peptide synthases (NRPSs).^9^ The prominent examples derived from this type of enzymatic machinery include but not limited to the anticancer drug epothilone,^10^ RNA polymerase inhibitor corallopyronin,^11^ mitochondrial respiratory inhibitor myxothiazol.^12^ Generally, PKS-NRPS megasynthetases perform sequential condensations of short carboxylic acids and/or (non-)proteinogenic amino acids to construct an amazing diversity of chemical skeletons. Each enzymatic module canonically incorporates one extender unit (either carboxyacyl monomers or amino acids) into the nascent polymeric chain. For each module, a gatekeeper domain selects the correct extender unit within each elongation step (AT domains in case of PKSs, and A domains for NRPSs).^13^ As a consequence, the organization and the order of multiple catalytic domains are co-linearly correlated with the NPs, and also greatly facilitated rational engineering to produce unnatural NPs.^9,14^

In this study, genome mining of *Myxococcus* sp. SDU36 prioritized a cryptic PKS-NRPS BGC (*mpd*). The genetic amenability of SDU36 enabled to elicit the expression of *mpd* through *in situ* promotor replacement. Subsequent comparative metabolic profiling facilitated the rapid characterization of a class of linear polyketidic amides **1–5** termed as myxopyromides A**–**E. Especially, compounds **3–5** were endowed with uncommon structural traits including aminobutan-2-one and/or pyrrolinone, and thus further enriched the chemical repertoire of myxobacterial NPs. We also proposed a biosynthetic model based on intensive bioinformatic analysis of the functional domains of central biosynthetic genes *mpdA–E*, and investigated biological functions for myxopyromides.

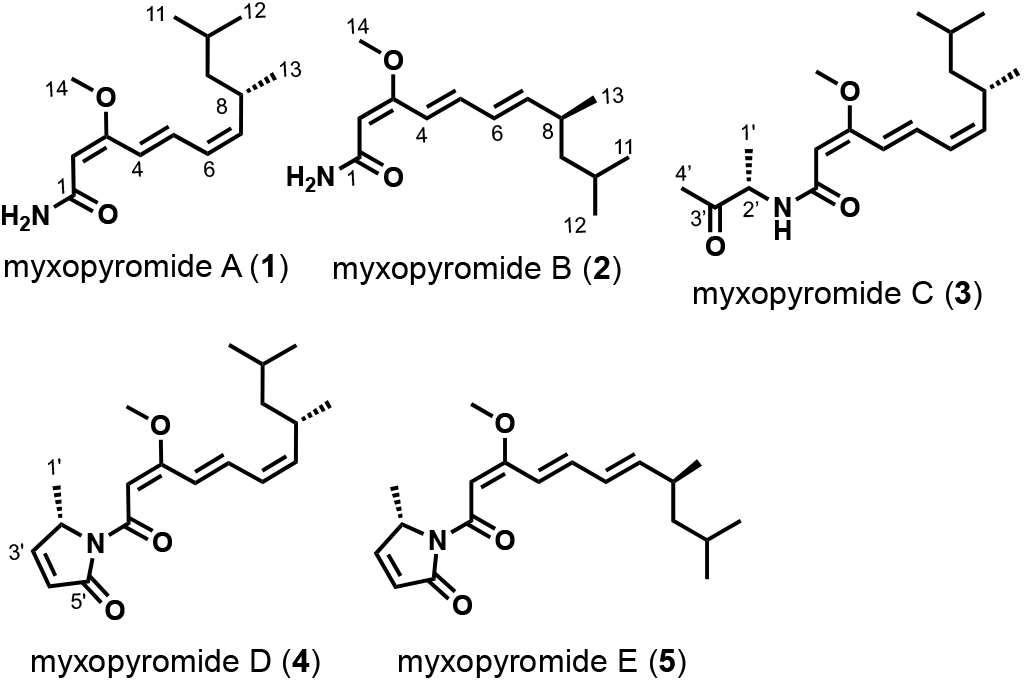

## RESULTS AND DISCUSSION

### Bioinformatic analysis of *mpd* and its promotor engineering

Genome mining of *Myxococcus* sp. SDU36 identified 28 BGCs during our previous investigation into the biosynthesis of myxadazole.^15^ Herein, a PKS-NRPS BGC designated *mpd* (∼43 kb, Table S1) in SDU36 particularly attracted our attention (Figure 1), because antiSMASH analysis of the domain architecture of five central biosynthetic genes *mpdA–E* indicated it very likely orchestrated a class of novel polyketidic compounds. To be specific, except *mpdD* that was an NRPS gene, the other four genes, namely *mpdA–C*, and *mpdE* encoded type I PKS megasynthetases. A grand total of six modules were specified by *mpdA–E*, and the substrate specificities of the AT domains and the single A domain were bioinformatically predictable. Therefore, the final product(s) of *mpd* were predicted to be assembled by six units of carboxyacyl monomers and one portion of alanine. Noteworthy, the arrangement of initiation module 1 encoded by *mpdA* seemingly deviated from the standard, since the loading AT_L_ domain should have been located in front of ACP_L_ but was instead adjoined with AT_1_. The concurrent existence of the loading and condensation AT domain in a same module was actually widespread in the hybrid PKS-NRPS BGCs of myxobacteria, as found in the assembly line of myxalamid,^16^ ajudazol,^17^ melithiazol,^18^ among others. Module 6 was supposed to be the last subunit in the pathway in view of the presence of the beacon TE domain. However, it was rather arduous to infer the chemical output of modules 2–5, because the irregular order and iterative use of functional domains is one of the idiosyncrasies of myxobacterial PKS.^19^

**Figure 1.**
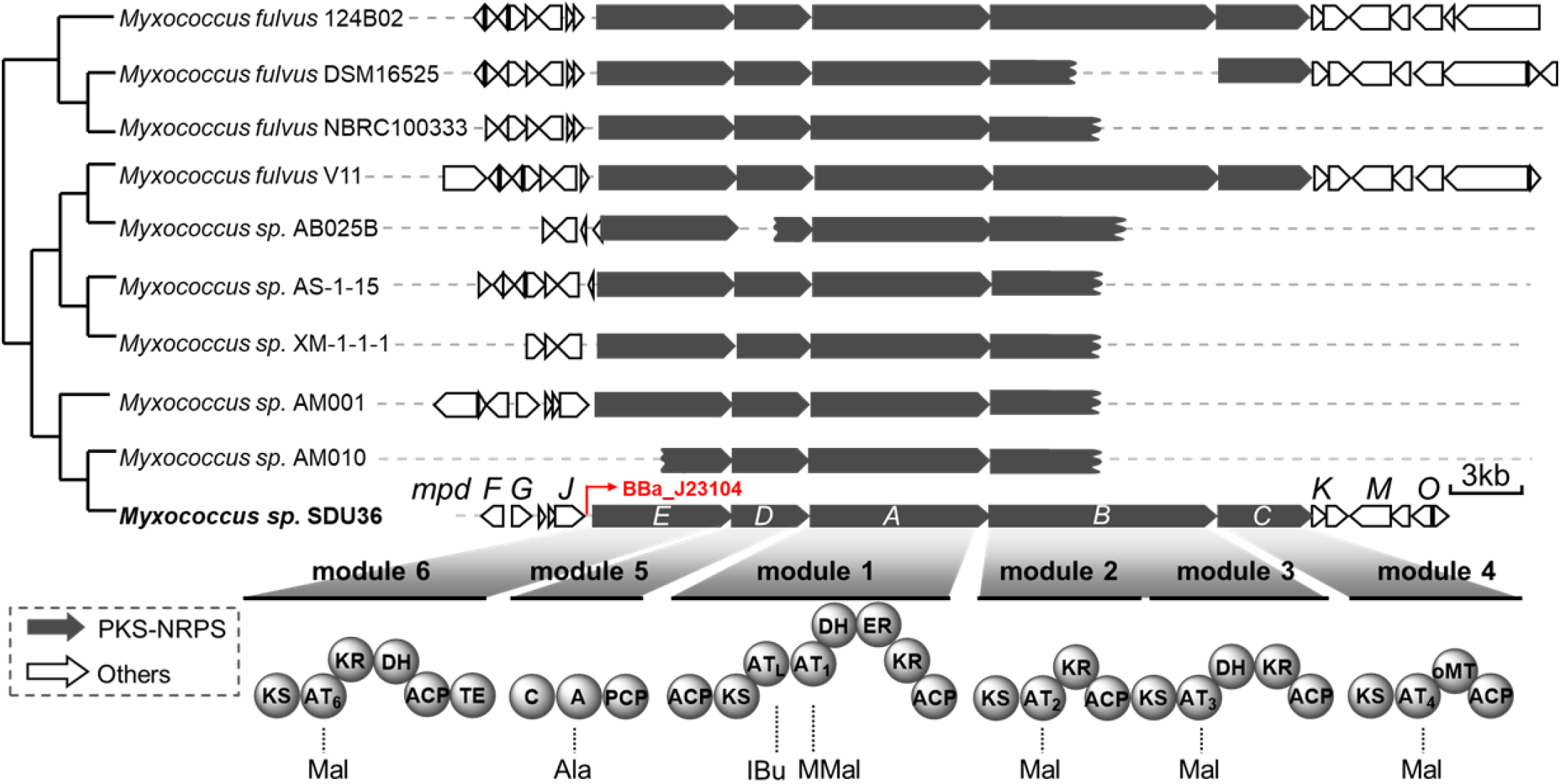
Distribution and organization of *mpd* clusters in myxobacteria. The conserved structural genes *mpdA-mpdE* were depicted in black and other auxiliary genes were in white. The schematic of the encoded PKS-NRPS subunits showed the component modules and domains. Domain abbreviations: KS, ketoacyl synthase; AT, acyltransferase; KR, ketoreductase; DH, dehydratase; ER, enoylreductase; oMT, *O*-methyltransferase; ACP, acyl carrier protein; C, condensation; A, adenylation; PCP, peptidyl carrier protein; TE, Thioesterase; IBu, isobutyryl-CoA; Mal, malonyl-CoA; MMal, methylmalonyl-CoA. Evolutionary distance of the cognate *mpd* BGCs were clustered by CORASON tool.^20^ The insertion position of the strong constitutive promotor BBa_J23104 in *mpd* of *Myxococcus* sp. SDU36 was rendered in red.

We conducted a bioinformatic inquiry wherein we searched for loci akin to *mpd* in all myxobacterial genomes. By using *mpdA* as the query to search genome databases, an additional nine cognate BGCs of *mpd* were discovered, which were highly conserved in the gene arrangement and domain architecture of the five central PKS-NRPS genes *mpdA–E*, in despite of variation in the auxiliary genes. All the 10 *mpd* BGCs were exclusively originated from the genus *Myxococcus*, implicating the unique physiological and/or ecological significance of the encoded metabolites. The phylogenetic grouping of these 10 BGCs was consistent with the evolutionary distance of the host strains.

We employed promotor engineering to elevate the expression of *mpd* by virtue of the genetic amenability of SDU36.^15^ The strong constitutive promoter BBa_J23104 ^7^ was used to replace the innate promoter of *mpdE* by single crossover homologous recombination, affording the promotor-engineered mutant SDU36-P_*mpd*_ (Figure S1). Subsequently, RT-PCR experiments showed the transcriptions of *mpdA-mpdE* were significantly bolstered by BBa_J23104 in SDU36-P_*mpd*_, otherwise unexpressed in the wild type SDU36 (Figure S2). Comparative HPLC-PDA profiling evidently presented a number of peaks SDU36-P_*mpd*_, which were absent in the SDU36. These metabolites gave a UV absorption spectrum similar to each other, indicating they were structural congers. This comparative metabolic profiling study prompted us to carry out the up-scale fermentation (30 L) of SDU36-P_*mpd*_ to clarify the chemical identity of the newly formed peaks. The repeated chromatographic isolation led to compounds **1** (5.1 mg), **2** (0.6 mg), **3** (1.3 mg), **4** (2.5 mg), and **5** (0.3 mg).

### Structure elucidation of compounds 1–5

Myxopyromide A (**1**) isolated as a yellowish oil, had the molecular formula C_14_H_23_NO_2_ established from its HRESIMS peak [M + H]^+^ at *m/z* 238.1794 (calcd. for C_14_H_24_NO_2_). The compound gave a UV absorption at λ_max_ 300 nm, supporting the occurrence of a long conjugation system. ^1^H NMR spectrum (Table 1) of **1** exhibited four conjugated vinylic protons at *δ*_H_ 7.54 (d, *J* = 15.6 Hz), 7.11 (dd, *J* = 15.6, 11.4 Hz), 6.09 (t, *J* = 11.4 Hz), and 5.36 (d, *J* = 10.2 Hz), judged by the coupling constant; one characteristic singlet at *δ*_H_ 5.19; one methoxy at *δ*_H_ 3.70; and three methyl doublets at *δ*_H_ 0.99 (d, *J* = 6.6 Hz), 0.90 (d, *J* = 6.6 Hz), and 0.88 (d, *J* = 6.6 Hz). Consistent with these assignments, the ^13^C NMR spectrum (Table 2) of **2** possessed seven sp^2^-hybridized carbons including five vinylic methine carbons, three methyl doublets, two sp^3^ methine carbons, one methylene, and one methoxy. Owing to the proficiency of protons, the gross structure of **1** was elucidated explicitly by the combination of COSY, HSQC, and HMBC (Figure 3A). To be specific, a long spin system from C-4 to C-13 was identified by COSY experiment, containing two conjugated double bonds *Δ*^4,5^ and *Δ*^6,7^. The β-methoxyacrylate moiety was supported by HMBC correlations from the *O*-methyl singlet (H-14) as well as the vinyl proton singlet (H-2) to *δ*_C_ 125.4 (C-4) and 164.7 (C-3), among others. An additional HMBC correlation between H-2 and carbonyl at *δ*_C_ 164.9 (C-6), and thus completed the identification of the gross structure of **1**. The geometry of the double bonds *Δ*^4,5^ and *Δ*^6,7^ were determined to be *E* and *Z*, respectively, judged by the ^1^H-^1^H coupling constants. The key NOESY correlation between H_3_-14 and H-2 established the *E* configuration of the double bond *Δ*^2,3^ (Figure S3). The absolute configuration at C-8 was determined as *S* based on ECD calculation (Figure 3B), which was consistent with the biosynthetic origin (see below). Therefore, myxopyromide A_1_ was determined as a linear functionalized fatty amide adorned with an *α,β,γ,δ,ε,ζ-* unsaturation, explanatory for its characteristic UV absorption at λ_max_ 300 nm.

**Table 1.**
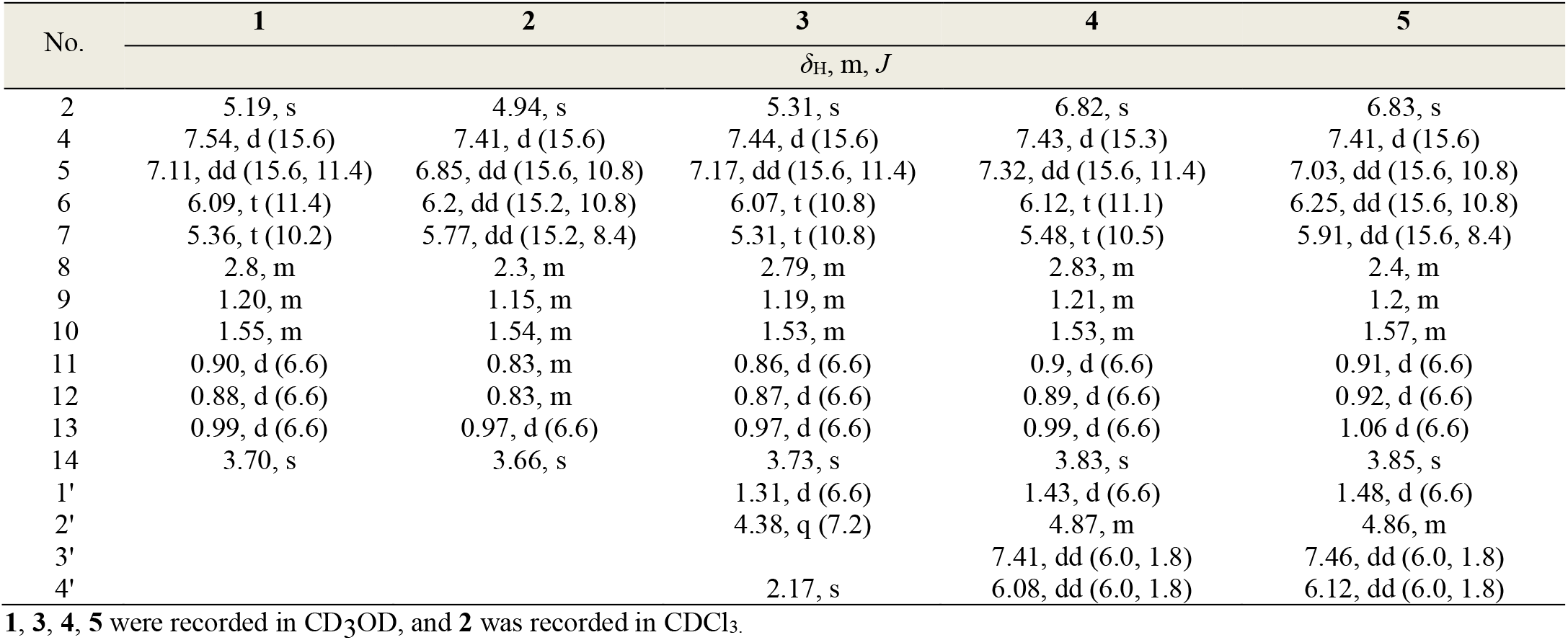
^1^H NMR Data (600 MHz, at 300 K) for myxopyromides.

**Table 2.** ^13^C NMR Data for myxopyromides.

| No. | 1 | 2 | 3 | 4 | 5 |
| --- | --- | --- | --- | --- | --- |
| | $\delta_{\text{C}}$, type | $\delta_{\text{C}}$, type | $\delta_{\text{C}}$, type | $\delta_{\text{C}}$, type | $\delta_{\text{C}}$, type |
| 1 | 163.4, C | 169.1, C | 165.4, C | 169.8, C | 170.8, C |
| 2 | 98.9, CH | 92.6, CH | 95, CH | 94.7, CH | 94.7, CH |
| 3 | 164.7, C | 165.9, C | 165.6, C | 166, C | 166.6, C |
| 4 | 126.0, CH | 122.1, CH | 125.4, CH | 125.6, CH | 124.2, CH |
| 5 | 129.2, CH | 136.1, CH | 130.5, CH | 132.7, CH | 138.9, CH |
| 6 | 128.3, CH | 128.7, CH | 128.1, CH | 127.7, CH | 130, CH |
| 7 | 143.2, CH | 146.6, CH | 144.1, CH | 145.5, CH | 148.6, CH |
| 8 | 31.6, CH | 35.2, CH | 27.9, CH | 31.5, CH | 36.6, CH |
| 9 | 48.2, CH<sub>2</sub> | 46.6, CH<sub>2</sub> | 48.1, CH<sub>2</sub> | 47.6, CH<sub>2</sub> | 47.8 CH<sub>2</sub> |
| 10 | 27.1, CH | 25.9, CH | 26.9, CH | 26.7, CH | 29.5, CH |
| 11 | 22.5, CH<sub>3</sub> | 23.4, CH<sub>3</sub> | 22.5, CH<sub>3</sub> | 22.2, CH<sub>3</sub> | 20.9, CH<sub>3</sub> |
| 12 | 23.8, CH<sub>3</sub> | 22.6, CH<sub>3</sub> | 23.7, CH<sub>3</sub> | 23.4, CH<sub>3</sub> | 23.5, CH<sub>3</sub> |
| 13 | 22.1, CH<sub>3</sub> | 21.1, CH<sub>3</sub> | 21.9, CH<sub>3</sub> | 21.6, CH<sub>3</sub> | 21.4, CH<sub>3</sub> |
| 14 | 55.6, CH<sub>3</sub> | 55.4, CH<sub>3</sub> | 55.8, CH<sub>3</sub> | 56.1, CH<sub>3</sub> | 56.5, CH<sub>3</sub> |
| 1' |  |  | 16.4, CH<sub>3</sub> | 17.9, CH<sub>3</sub> | 18.5, CH<sub>3</sub> |
| 2' |  |  | 56.1, CH | 59.5, CH | 60.0, CH |
| 3' |  |  | 210.5, C | 155.4, CH | 155.8, CH |
| 4' |  |  | 26.2, CH<sub>3</sub> | 125.9, CH | 126.4, CH |
| 5' |  |  |  | 171.9, C | 172.3, C |

Myxopyromide B (**2**) had the molecular formula C_14_H_23_NO_2_ according to the HRESIMS [M + H]^+^ peak at *m/z* 238.1809 (calculated for C_14_H_24_NO_2_. The ^1^H and ^13^C NMR spectra closely resembled those of **1**, and a detailed 2D NMR interpretation confirmed that they shared the same C-C connectivity. However, the geometry of *Δ*^6,7^ in **2** was *E* instead of *Z* based on the large coupling constant 15.2 Hz between H-6 and H-7. Myxopyromide C (**3**) was also obtained as a yellowish oil. The molecular formula was determined to be C_18_H_29_NO_3_ by HR-ESITOFMS. While the ^1^H NMR spectrum of **3** contained the full set of signals found in **1**, an additional three resonances presented, including two methyls at *δ*_H_ 2.17 (s, H-4’) and 1.31 (d, *J* = 6.6 Hz, H-1’), and 4.38 (d, *J* = 7.2 Hz, H-2’). The ^13^C NMR spectrum of **3** showed four more carbon signals than **1**, including a ketone at *δ*_C_ 210.5, and three aliphatic carbons at *δ*_C_ 26.2 (C-4’), 56.1 (C-2’), and 16.4 (C-1’), corresponding to the three new protons as resolved by HSQC experiment. This strongly suggested **3** was a derivative of **1** with an additional appendage at the carboxyl terminal. The identity of the new structural moiety was determined as a 3-aminobutan-2-one, based on the key HMBC correlations from H-1’ and H-4’ to the C-2’ and C-3’, respectively (Figure 3A). Although the HMBC correlation between H-2’/C-1 (*δ*_C_ 165.4) was not observed, the linkage of C-1 to C-2’ should be through a nitrogen atom considering the molecular formula and chemical shifts of H-2’. In consideration that 3-aminobutan-2-one was originated from L-alanine (see below), the absolute configuration at C-2’ was determined to be *S*. The structure of **3** is intriguing because, to our knowledge, it represents the first natural topology combining aminobutan-2-one with fatty acyl moieties.

Myxopyromide D (**4**) was isolated as a yellowish oil, and determined to have a molecular formula of C19H27NO3 according to the HRESIMS [M + H]^+^ peak at *m/z* 318.2034 (calculated for C19H28NO3). The NMR data for **4** contained the full set of fatty acyl group for **1**, indicating that the **4** was another amide derivative of **1**. The connectivity of the five extra carbons C-1’∼C-5’ that were present in **4** but absent in **1** was accomplished by the joint use of COSY and HMBC. A contiguous spin system that possessed a deshielded doublet methyl (*δ*_H_ 1.43, H-1’, d, *J* = 6.6 Hz), one aliphatic methine H-2’, and two vinylic methine protons H-3’ and H-4’ was detected by COSY experiment. The chemical shifts of H-2’ (*δ*_H_ 4.87, m) and C-2’ (*δ*_C_ 59.5) were diagnostic of the attachment to a heteroatom. The chemical shifts and coupling constants of H-4’ (*δ*_H_ 6.08, dd, *J* = 6.0, 1.8 Hz) and H-3’ (*δ*_H_ 7.41, dd, *J* = 6.0, 1.8 Hz) suggested they formed a polarized *Z* double bond and thus an *α,β*-unsaturated amide functionality. This moiety was confirmed by HMBC correlations from protons H-3’ and H-4’ to a carbonyl carbon at *δ*_C_ 171.9 (C-1’) (Figure 3A). Further HMBC correlation between H-2’/C-5’allowed the closure of a pyrrolinone ring. Although HMBC correlations between H-2’/C-1 was not observed, pyrrolinone ring should be amidated with **1** through the carbonyl carbon (*δ*_C_ 169.8), considering the molecular formula of **4**. As in **3**, the absolute configuration of C-2’ in **4** was assigned as *S*, based on the ECD calculation (Figure 3B) and biosynthetic origin of this carbon from L-alanine. This stereochemistry assignment was further endorsed by the same structural unit present in ypaoamide C ^21^ and palmyrrolinone,^22^ eliamid,^23^ and jamaicamides.^24^

Myxopyromide E (**5**) shared the same molecular formula of C19H27NO3 with **4**. The ^1^H and ^13^C NMR spectra of **5** closely resembled those of **4**. They marginally differed from each other at the geometry of double bond *Δ*^6,7^ based on the splitting pattern and ^1^H-^1^H coupling constants of H-6 and H-7 (Table 2). It was noteworthy that the repertoire of NPs featuring pyrrolinone is very limited, and especially, myxopyromide D_1_ (**4**) and myxopyromide D_2_ (**5**) further expands the complexity of myxobacterial NPs.

### Biosynthetic model of myxopyromides

The elaborated structure elucidation and sequence analysis of *mpd* compelled us to propose the biosynthetic scheme for myxopyromides (Figure 4). The assembly was initiated by the first AT (AT_L_) of MpdA that recognizes the starter unit isobutyryl-CoA (IBu) and then transfers to the initial ACP_L_. The second AT (AT_1_) of MpdA loads methylmalonate (MMal) that is subsequently used for the first chain extension step. The remaining elongation steps can be assigned readily to the composite modules of subunits MpdB–MpdE, wherein the integral AT or A domains recruit appropriate extender units, respectively (Figure 1). Finally, a dedicated TE domain at C-terminus of MdpE is tentatively attributed to the lactamation and chain release to afford the five-membered pyrrolinone ring. This proposition was tested by the phylogenetic analysis of TE domains, wherein *mpd* indeed clustered together with *jam* that encode jamaicamides containing the same pyrrolinone ring (Figure S4).^24^ The *mpd* pathway is organized in a remarkably colinear arrangement with respect to the structure of myxopyromide D_1_ (**4**), except for the absence of an integral DH in the first module of MpdB that would produce the *cis* double bond between C-6 and C-7. This reaction is presumably complemented by the iterative action of DH domain in the downstream module 3, because this biosynthetic trajectory is quite commonplace in myxobacterial PKS (e.g. spirangiene,^25^ epothilone,^10^ chivosazol ^26^, and archangiumide ^27^). A high level of colinearity between the genotype and the chemotype also reflects in the stereocontrol. Sequence alignment of ER domain in MdpA revealed a distinct tyrosine residue (Y^43^) in the catalytic triad (Figure S5),^28^ which results in an *S* configuration of the methyl branch at C-8 position of **1**–**5** as confirmed by ECD calculations. For the stereochemistry of double bonds, the B-type (KR_2_ and KR_6_) or A-type (KR_1_ and KR_3_) KR domains (Figure S6) catalyze the generation of *E* or *Z* configuration,^29^ respectively, in agreement with the geometry at the double bonds *Δ*^4,5^ (*E*), *Δ*^6,7^ (*Z*), and *Δ*^3’,4’^ (*Z*) as determined by spectroscopic methods. We hypothesized that the *E* configuration of *Δ*^6,7^ in the minor products **2** and **5** was derived from the isomerization of the major products **1** and **4** by the action of the isomerase MpdN, respectively. The double bond *Δ*^2,3^ (*E*) is thought to be generated by the oMT domain in MdpC which probably methylate the enol form of the nascent diketide intermediate. This mechanism has been postulated to also direct the biosynthesis of *E-*configured *β*-methoxyacrylate functionalities of myxothiazol,^30^ melithiazol,^18^ and ajudazols,^17^ among others. The absence of an epimerase domain in MpdD provides a rationale for L-alanine incorporation into the growing polymeric chain, and thus eventually leads to the *S* configuration of the *N*-bearing chiral center C-2’ in **3**–**5**.

While **4** is thought to be the final product of the whole *mpd* biosynthetic pathway, **1**–**3** are obviously the shunt products resulting from the premature release of partially-assembled polyketide chains. After scrutinized interrogation of *mpd*, we speculated that an *α, β-*fold hydrolase gene *mpdL* probably gets involved in the truncation of assembly line by mimicking the function of a type II thioesterase (TE). The resultant β-ketone acid is prone to spontaneous decarboxylation to generate the 2-aminobutanone functionality in **3**, which is probably processed by a putative oxidase MpdQ to generate **1**. Intriguingly, it seems that MpdL outcompetes MpdE-TE domain to off-load the intermediate tethered on MpdE-ACP domain prior to the action of MpdE-KR domain, since the major product of *mpd* is **1** rather than **4**. The flexibility in the chain release from the modular PKS assembling machinery engenders the structural perturbation to the programmed chemistry of *mpd*, which might represent a common but anteriorly overlooked strategy adopted by microbes for diversity-oriented biosynthesis, in addition to the well-known post-PKS modification.

### Biological function of myxopyromides

Myxopyromides (**1**–**5**) were inactive against *Staphylococcus aureus, Acinetobacter baumannii, Escherichia coli* K-12, and *Candida albicans* at 40 μg/disc during agar diffusion assay. They showed weak cytostatic activity towards the three cell lines Hela, PC, and MCF-7, with IC_50_ value at ∼30 μM (Table S2). These preliminary assays compelled us to investigate the potential physiological functions of myxopyromides in milieu. Therefore, the morphologies characteristic to myxobacteria, including growth curve, social motility (S-motility), adventurous motility (A-motility), exopolysaccharides secretion, fruiting body formation,^1^ were compared between SDU36 and SDU36-P_*mpd*_, wherein no significant differences were observed between these two strains (data not shown). Nonetheless, SDU36-P_*mpd*_ exhibited an evidently stronger ability to prey on *E. coli* K-12 than SDU36 grown on an agar plate. When SDU36 culture was supplemented with 0.5 mg/ml of crude extract of SDU36-P_*mpd*_, we observed a likewise elevated predation of SDU36 towards *E. coli* K-12 (Figure 5). We reasoned that the alleged phenotypic occurrence was ascribed to the production of myxopyromides in SDU36-P_*mpd*_, since the mere metabolic difference of SDU36 and SDU36-P_*mpd*_ was the suite of myxopyromides produced by the latter

(Figure 2). Regretfully, owing to the limited amount of the purified **1**–**5**, it was impractical to evaluate and/or compare the predation-enhancing potency of individual compound. Noteworthy, the observed small molecule-mediated predation was intriguing but not unprecedented in the paradigm of myxobacteria, because myxovirescin and corallopyronin have been reported to functionate as chemical weapons to coordinate predation of *Myxococcus xanthus* and *Corallococcus coralloides*, respectively.^31^ Instead of antibiotics, myxopyromides are more likely to act as signaling molecules, since this class of polyketides were devoid of antibacterial efficacies. It is tempting to understand mechanistically how myxopyromides contribute to the predation phenotype of SDU36 at the molecular level.

**Figure 2.**
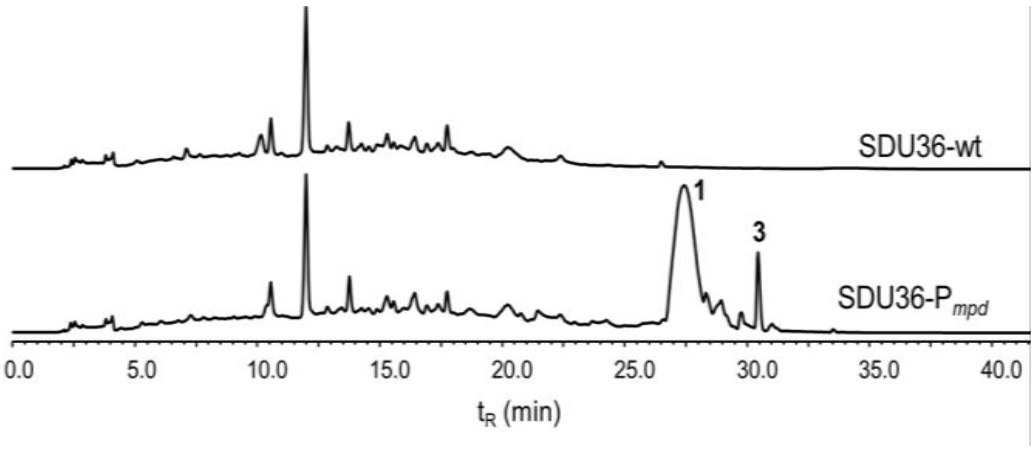
Metabolic comparison of SDU36 wild type and mutants by HPLC-PDA. Each trace was recorded at 280 nm, and the region containing BBa_J23104-induced metabolites were dash boxed highlighted.

**Figure 3.**
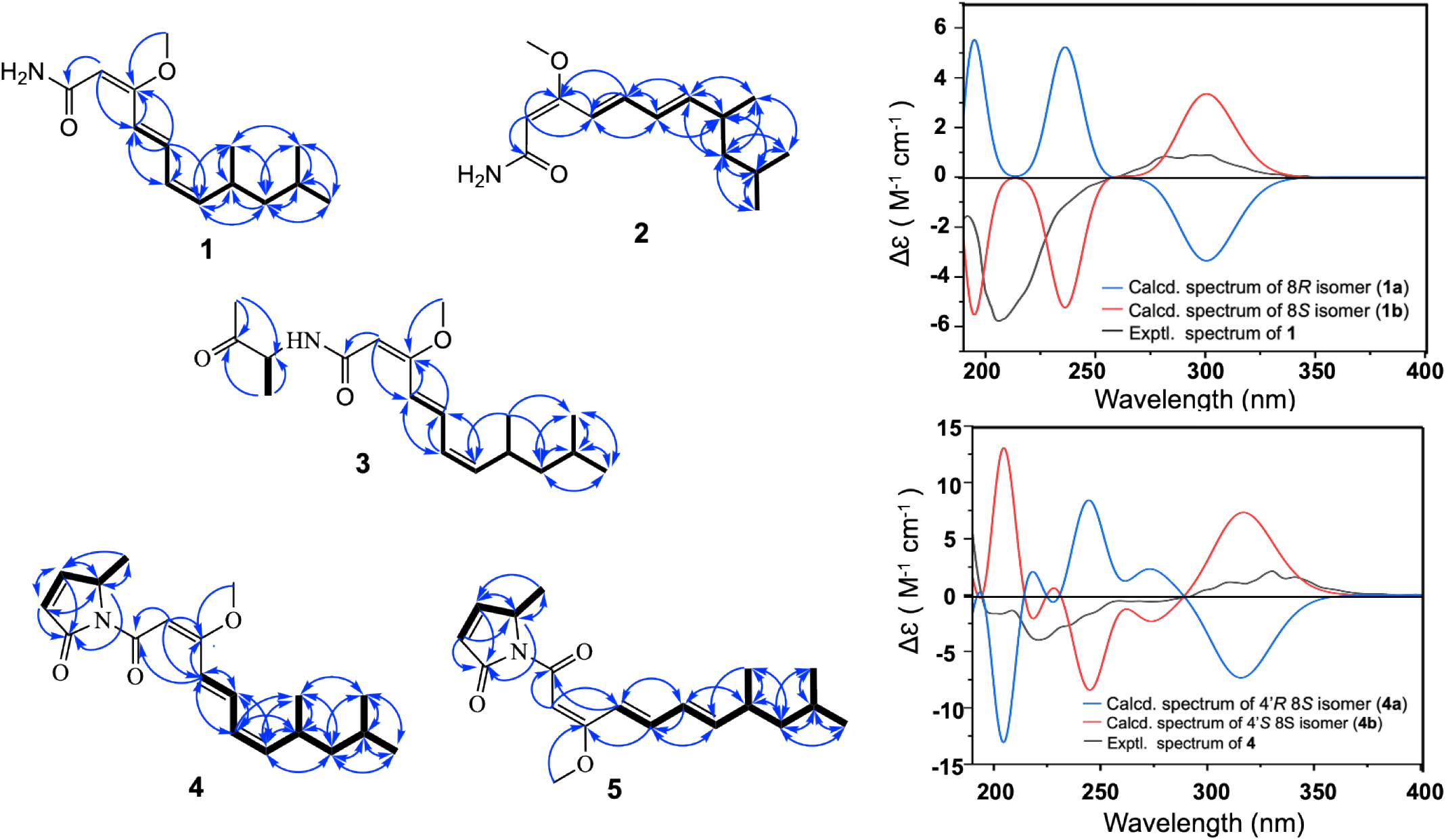
Structure elucidation of myxopyromides 1–5. (**A**) HMBC () and COSY () correlations of myxopyromides **1**–**5**. (**B**) experimental and calculated ECD spectra of **1** and **4**

**Figure 4.**
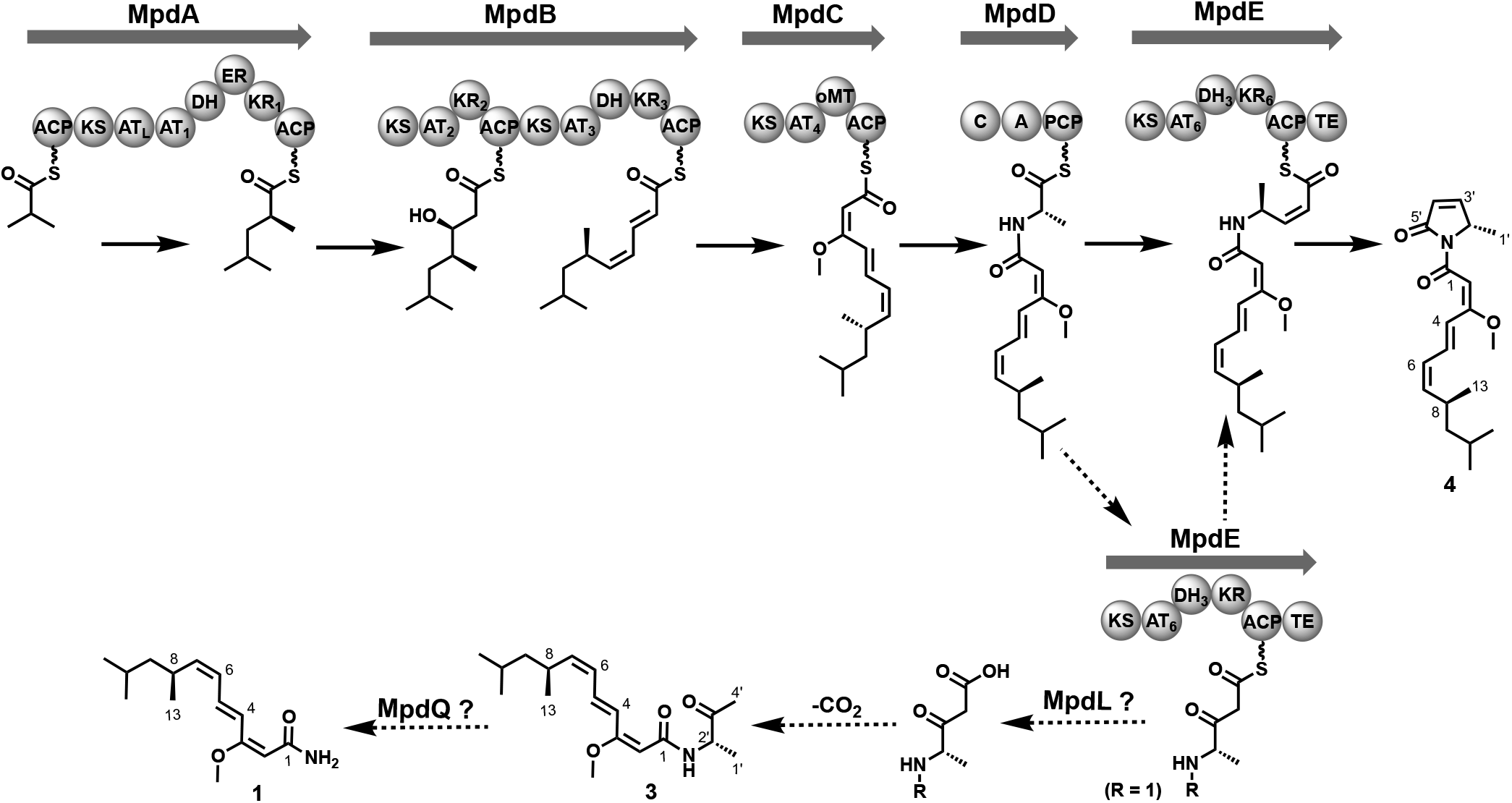
Model for the biosynthesis of myxopyromides. The intact pathway to myxopyromide D_1_ (**4**) is depicted in solid arrow, whereas the bifurcation pathways to **1**–**3** were depicted in dashed arrow. The stereochemistry of β-hydrxoyl, *a*-methyl, double bonds, and amino acid (_L_-alanine) of the polyketidic intermediate is indicated based on the sequence analysis of functional domains (KR, ER). The note for domain abbreviations referred to Figure 1.

**Figure 5.**
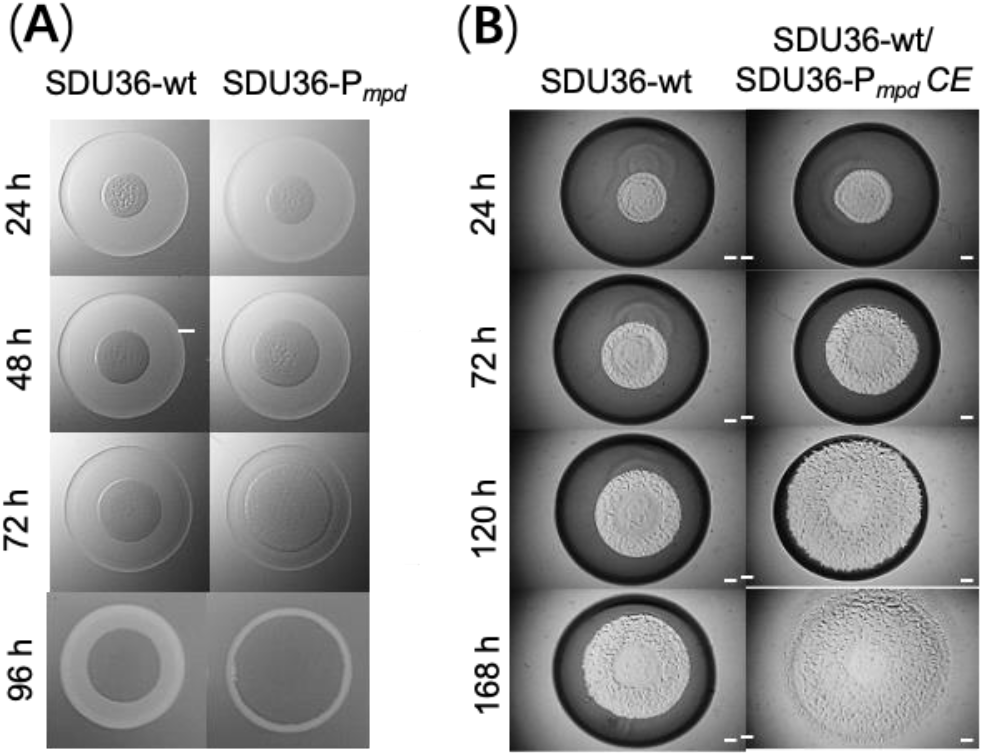
Myxopyromides enhanced the predatory ability of *Myxococcus* sp. SDU36. (**A**) Time-course predation circle assay of SDU36 wild type and the myxopyromides-producing strain SDU36-P_*mpd*_, wherein predation circle of SDU36-P_*mpd*_ was invariably larger than SDU36-P_*mpd*_ after 24 hours; (**B**) Time-course predation circle assay of SDU36 wild type that was supplemented with crude extract of SDU36-P_*mpd*_, which enhanced the predatory ability of SDU36. The predation circle of SDU36 and/or SDU36-P_*mpd*_ that were grown on 1.5 % agar TPM plate was observed using a stereomicroscope. The colony size reflected the predatory ability of the tested strains. Bar in the picture was 1 mm.

## CONCLUCSION

Myxobacteria are amongst the top producers of NPs, providing fertile grounds for novel PKS-NRPS biosynthetic machineries ripe for further exploration. We identified a mixed PKS-NRPS BGC (*mpd*) by genome mining of *Myxococcus* sp. SDU36, and the feasibility of the expedient genetic system enabled to switch on its expression by promotor exchanging. The subsequent comparative metabolic profiling streamlined the characterization of myxopyromides, a class of linear polyketidic amides featuring uncommon pyrrolinone warhead or aminobutanone. Since myxobacteria are a family of biomedical resources that are highly underexplored, we have provided an additional proof of principle that there is still a bright future ahead for our recently characterized myxobacterial constitutive promotors^7^ applicable to awaken cryptic BGCs. Moreover, as synthetic biology is rapidly changing the landscape of discovery of bacterial NPs,^32^ the previously unappreciated chain release pattern implicated in the biosynthesis of myxopyromides, would be leveraged to enhance the structural diversity orchestrated by a broad range of modular PKS and/or NRPS machinery. Last but not least, this study enriched our knowledge in relation to small molecules influencing the developmental processes of myxobacteria, which would in turn favor in depth exploitation of this pharmaceutically important resource.

## EXPERIMENTAL SECTION

### General Experimental Procedure

NMR spectra were recorded on a Bruker AVNEO 600 MHz calibrated to a residual CD_3_OD (3.31 ppm) and CDCl_3_ (7.26 ppm). The HR-Q-TOF ESIMS analyses were performed on a rapid separation liquid chromatography system (Dionex, UltiMate3000, UHPLC) coupled with an ESI-Q-TOF mass spectrometer (Bruker Daltonics, Impact HD). HPLC analysis was performed with a Thermo Scientific Ultimate 3000 HPLC system equipped with a column (Thermo Fisher Scientific, C_18_, 4.6 × 250 mm, 5μm). HR-Q-TOF ESIMS instruments equipped with a C_18_ column (Thermo Fisher Scientific, C_18_, 250 × 4.6 mm, 5μm) were used. Samples were detected by the UV absorbance at 210, 254, 280, and 365 nm. Silica gel 60 F254 (Merck, Darmstadt, Germany) was used for TLC analysis, migrated with CHCl_3_/MeOH (10:1), and visualized with anisaldehyde/sulfuric acid reagent. According to specific experiments, all organic solvents and chemicals were of analytical or HPLC grade. Polymerase chain reactions (PCR) catalyzed by Phanta Super-Fidelity DNA Polymerase (Vazyme) were performed on a GE4852T Gene-Explorer Touch series thermal cycler (Bio-Gener Technology, Hangzhou, Zhejiang, China). PCR products were gel-purified using Gel Extraction Kit (Omega, GA, USA). Gibson cloning was based on ClonExpress Ultra One Step Cloning Kit (Vazyme, Najing, China). TIANprep Rapid Mini Plasmid Kit (TIANGEN BIOTECH, Beijing, China) was used for plasmid extraction. Cells were disrupted using a high-pressure homogenizer NanoGenizer (Genizer LLC, Irvine, CA, USA).

### Bioinformatic Analysis of *mpd*

All the 268 sequenced genomes of *Myxococcales* downloaded from the NCBI RefSeq database. The amino acid sequence of MpdA was queried against the database using BLASTp to retrieve the homologs. The hits with BLAST value of “query coverage” and “percent identity” ≥85% was deliberately ascertained to be the sought-after candidates. The genome sequences that contained MpdA homologs were analyzed by offline antiSMASH analysis.^33^ The counterparts of *mpd* were manually detected according to the domain architecture of the five central PKS-NRPS genes (*mpdA–E*). The CORASON tool ^20^ was employed to generate a multilocus phylogeny of all the *mpd* BGCs. The .fasta file of the gene *mpdA* was used as the query gene, and the *mpd* from *Myxococcus* sp. SDU36 was used as the reference BGC. The settings of CORASON: e_value = e^−169^, and the other parameters were set as default. The MEGA 7.0 software was used to the visualize the phylogenetic tree generated by CORASON.

### Promotor Engineering of *mpd*

The basal plasmid pBa-1^7^ (Table S3) that contained the strong constitutive promotor BBa_BBa_J23104 was used for the promotor exchange of *mpd*. The primer pair cpBa-F/cpBa-R (Table S4) was used to PCR linearize the plasmid pBa-1. The initial ∼1 kb of the gene *mpdE* starting from ATG codon was PCR amplified with primer pairs Amyp-F/Amyp-R. The two resulting amplicons were ligated by Gibson cloning via 20 bp overlap bases to afford the construct pBa-*mpd*, wherein the *mpdE* fragment was under the control of BBa_J23104. The activation vector was electroporated into SDU36 wild type that was grown in liquid CTT medium (10 g/L casitone,1.97 g/L MgSO_4_ × 7H_2_O, 10 mM Tris-HCl, 1 mM K_2_HPO_4_/KH_2_PO_4_, pH 7.6) at 30 °C, 200 rpm for 24 h, basically followed our previous protocol.^7^ Single-crossover recombination mutants were selected by 40 μg/mL kanamycin (Figure S1). The correct genotype SDU36-P_*mpd*_ was checked by colony PCR and further verified by Sanger Sequencing.

### Check Transcription of *mpdA*–*E* by RT-PCR

SDU36 wild type and the mutant SDU36-P_*mpd*_ were in parallel inoculated in liquid CTT medium, with the latter was kept in kanamycin selection pressure. After 48 h of growth, appropriate amount of culture was transferred into 50 mL of fresh CTT medium to achieve OD_600_ value at ∼0.04. The diluted culture was grown for an additional 48 h, and then 1.5 mL of culture was harvested by centrifugation. RNA was extracted immediately using a Bacteria Total RNA Isolation Kit (Sangon Biotech) according to the manufacturer’s instructions. The purified RNA extracts were reverse-transcribed to cDNA. RT-PCR was performed using the resulting cDNA as template, around 200 bp fragments were amplified (Figure S2). The primers were listed in Table S4. Each experiment was done in triplicate.

### Culturing Conditions and Metabolites Extraction of SDU36 Strains

The *mpd*-engineered *Myxococcus* sp. SDU36 mutant as wells as wild type were first cultivated on CTT agar plate for 3 days at 30 °C, respectively. Mycelium of the colony was inoculated into a 50 mL of liquid CTT medium, which was continuously shaken at 200 rpm, 30 °C for another 3 days. A 10 mL of seed culture was diluted in 100 mL of liquid VY/2 medium (5 g/L baker’s yeast, 1 g/L CaCl_2_, 0.5 mg/L vitamin B_12_, 1.97 g/L 4-(2-hydroxyethyl)-1-piperazineethanesulfonic acid (HEPES), pH 7.2), and then continuously shaken at 200 rpm, 30 °C for 7 days. Antibiotics (50 μg/mL kanamycin and/or 25 μg/mL chloramphenicol) were added into the media cultivating the mutant. The cultures were harvested by centrifugation at 5500 ×*g* for 5 min. The compounds in the supernatant were adsorbed with 1 g of HP-20 resin. The resin was collected with cheesecloth and dried at room temperature for 24 h. The column-packed resin was first rinsed with 15 mL of H_2_O, and then eluted with 15 mL of methanol. Next, organic solvent was evaporated under vacuum, and the afforded crude extracts were analyzed by HPLC-PDA and/or LC-MS.

### HPLC-PDA analysis

HPLC analysis was performed with a Thermo Scientific Ultimate 3000 HPLC system equipped with a column (Thermo Fisher Scientific, C_18_, 4.6 × 250 mm, 5 μm). The binary mobile phase consisted of water (A) and methanol (B, HPLC grade) in a linear gradient program from 5% B to 100% B in 46 min at a flow rate of 0.8 mL/min. The methanol concentrations were changed as follows: 5– 100% B (0–30 min), 100% B (30–40 min), 5% B (40–46 min). Chromatograms were recorded at 210, 254, 280, and 365 nm, respectively. The injection volume was 10 μL.

### LC-MS Analysis

LC-MS analysis was performed on a UHPLC system (Thermo Scientific™ Vanquish) coupled to a Q Exactive™ HF-X Hybrid Quadrupole-Orbitrap™ Mass Spectrometer (Q Exactive™ HF-X, ThermoFisher Scientific) equipped with an ESI source. The chromatographic separation was done on a column with solid-core particles (ACQUITY UPLC HSS T3 (100 mm × 2.1 mm i.d., 1.8 μm; Waters, Milford, USA). The binary mobile phase consisted of water (A) and isopropanol/acetonitrile (1:1, B). The gradient was 5% to 100% solvent B in A in 0−5.5 min, isocratic plateau at 100% solvent B in 5.5-7.5 min, and re-equilibration at 5% solvent B in 7.5-10 min, at a constant flow rate of 400 μL/min. ESI source parameters were used as follows: heater temperature 425 °C, sheath gas flow rate 50 arb, Aux gas flow rate 13 arb, Spray voltage ±3500 V, and normalized collision energy 20, 40, 60 eV.

### Chromatographic Isolation of Myxopyromides

Up-scale fermentation of SDU36-P_*mpd*_ in 30 L of VY/2 suppled with 50 μg/mL kanamycin was harvested by centrifugation at 8000 rpm. The resulting supernatant was extracted by 0.6 kg of Dianion HP-20 resin. The methanol-recovered crude extract was applied to reverse-phase preparative medium-pressure liquid chromatography (Buchi, Flawil, Japan) to give 10 fractions (Fr1−Fr10). Fr5-7 was purified by HPLC (Thermo Fisher Scientific, C_18_, 250 × 4.6 mm, 5μm; solvent A: H_2_O (40%); solvent B: MeOH (60%); flow rate 1.8 mL/min; 30 °C; UV detection at 280 nm), to afford compounds (5.1 mg), **2** (0.6 mg), **3** (1.3 mg), **4** (2.5 mg), and **5** (0.3 mg).

*Myxopyromide A* (**1**): yellowish oil; 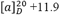(c 0.05, MeOH); UV (MeOH) λ_max_ 217, 297 nm; ECD (MeOH): 216 (Δε +6.04), 294 (Δε +0.90); IR *v*_max_ 3330, 3164, 2955, 1662, 1597, 1418, 1339, 1247, 1207, 1134, 1060, 994, 957, 858, 805, 770, 704 cm^−1^; ^1^H NMR (600 MHz, methanol-*d*_4_) and ^13^C NMR (150 MHz, methanol-d4) data were showed in Table 1 and Table 2; HRESIMS (positive mode) *m/z* 238.1794 [M + H] ^+^ for **1** (calcd. for C_14_H_24_NO_2_, 238.1802).

*Myxopyromide B* (**2**): yellowish oil; 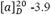(c 0.05, MeOH); UV (MeOH) λ_max_ 219, 300 nm; ECD (MeOH): 214 (Δv +5.62), 290 (Δv -3.45); IR v_max_ 3334, 2956, 2869, 1659, 1456, 1381, 1259, 1034, 802 cm^−1^; ^1^H NMR (600 MHz, CDCl_3_) and ^13^C NMR (150 MHz, CDCl_3_) data were showed in Table 1 and Table 2; HRESIMS (positive mode) *m/z* 238.1809 [M + H] ^+^ for **2** (calcd. for C_14_H_24_NO_2_, 238.1802).

*Myxopyromide C* (**3**): yellowish oil;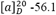 (c 0.05, MeOH); UV (MeOH) λ_max_ 221, 301 nm; ECD (MeOH): 213 (Δ*v* -12.12), 296 (Δ*v* +3.35); IR *v*_max_ 3293, 2963, 1651, 1539, 1453, 1200, 1136, 837, 799, 721cm^−1^; ^1^H NMR (600 MHz, methanol-*d*_4_) and ^13^C NMR (150 MHz, methanol-*d*_4_) data were showed in Table 1 and Table 2; HRESIMS (positive mode) *m/z* 308.2227 [M + H] ^+^ for **3** (calcd. for C_18_H_30_NO_3_, 308.2220).

*Myxopyromide D* (**4**): yellowish oil; 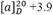 (c 0.05, MeOH); UV (MeOH) λ_max_ 250, 337 nm; ECD (MeOH): 221 (Δ*v* -3.98), 331 (Δ*v* +2.15); IR *v*_max_ 3350, 2959, 1719, 1665, 1558, 1367, 1333, 1297, 1199, 1137,1038, 997, 820, 756, 702 cm^−1^; ^1^H NMR (600 MHz, methanol-*d*_4_) and ^13^C NMR (150 MHz, methanol-*d*_4_) data were showed in Table 1 and Table 2; HRESIMS (positive mode) *m/z* 318.2034 [M + H] ^+^ for **4** (calcd. for C_19_H_28_NO_3_, 318.2064).

*Myxopyromide E* (**5**): yellowish oil; 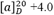(c 0.05, MeOH); UV (MeOH) λ_max_ 251, 333 nm; ECD (MeOH): 223 (Δ*v* +0.64), 260 (Δ*v* -0.70); IR vmax 3350, 2926, 2855, 1716, 1667, 1559, 1456, 1369, 1199, 821, 756, 701 cm^−1^; ^1^H NMR (600 MHz, methanol-*d*_4_) and ^13^C NMR (150 MHz, methanol-*d*_4_) data were showed in Table 1 and Table 2; HRESIMS (positive mode) *m/z* 318.2052 [M + H] ^+^ for **5** (calcd. for C_19_H_28_NO_3_, 318.2064).

### ECD Calculations

Conformational analysis of the candidate structures was performed using the MMFF94s force field. The conformers for each structure within 4 kcal/mol energy window were subjected to DFT geometry optimization at the B3LYP/6-31G* level of theory by using Gaussian 16 software package.^34^ The optimized conformers were Boltzmann averaged based on Gibbs free energy and obtained equilibrium populations at 298K. The theoretical calculations of ECD were performed using the TDDFT at the ωB97xd/TZVP level in MeOH with the IEFPCM model. The ECD spectrum of all conformers were averaged based on Boltzmann and simulated by Multiwfn.

### Antimicrobial and Anti-cancer Bioassays

Antimicrobial activity test by agar diffusion assay and anti-cancer assay by MTT method followed our previous protocol.^15,27,35^ The cytotoxicity was tested on three cell lines A549, HeLa and MCF-7 using the MTT assay. Briefly, the tested cells were seeded in a 96-well microtiter plate (∼2,500 cells/well) and incubated at 37 °C for 24 h with 5% CO2. Compo unds 1, and 3–5 were added to each well at final concentrations ranging from 20 to 0.01 μM. After a 48 h incubation, the cells were incubated with 10 μL of MTT (5 mg/mL) for 4 h. Formazan product was then solubilized with 100 μL of DMSO, and the absorbance was measured at 570 nM by a plate reader (Bio-Rad, USA). IC50 values were calculated from the plotted results, with untreated cells being set at 100%. The solvent DMSO was used as the negative control, and doxorubicin was used as positive control. All experiments were performed in triplicate.

### Central Predation Assay of SDU36 and SDU36-P_*mpd*_

An appropriate amount of *Myxococcus* predator (SDU36 or SDU36-P_*mpd*_) and prey *E. coli* K-12 were washed three times with TPM buffer (1.97 g/L MgSO_4_.7H_2_O, 10 mM Tris-HCl, 1 mM K_2_HPO_4_/KH_2_PO_4_, pH 7.6). The OD_600_ of *Myxococcus* was adjusted to 35 (about 5 × 10^9^ cells/mL) and the OD_600_ of *E. coli* K12 was adjusted to 100 (about 1 × 10^11^ cells/mL). 35 μL of prey was added onto the 1.5 % TPM agar plate and dried in the clean bench. Then, 2 μL of predator was added to the center of the prey plaque. The plates were placed in an incubator at 30 °C. The predation was observed and photographed at 12 hours intervals using a stereomicroscope. Each experiment was performed in triplicate.

## ASSOCIATED CONTENT

### Supporting Information

The Supporting Information is available free of charge on the ACS Publications website.

Plasmids, bacterial strains, and primers; Schematic for promotor engineering; RT-PCR verification; bioinformatic analysis of functional domains; NMR, HRMS, UV, and IR data of **1**−**5** (PDF).

## AUTHOR INFORMATION

### Authors

**Luo Niu** − *State Key Laboratory of Microbial Technology, Institute of Microbial Technology, Shandong University, 266237 Qingdao, P*.*R. China*

**Wei-Feng Hu** − *State Key Laboratory of Microbial Technology, Institute of Microbial Technology, Shandong University, 266237 Qingdao, P*.*R. China*

**Wen-Juan Zhang** *− State Key Laboratory of Microbial Technology, Institute of Microbial Technology, Shandong University, 266237 Qingdao, P*.*R. China*

**Zhao-Min Lin** *− Institute of Medical Science, The Second Hospital, Cheeloo College of Medicine, Shandong University, Jinan 250033, P*.*R. China*.

**Jing-Jing Wang** − *State Key Laboratory of Microbial Technology, Institute of Microbial Technology, Shandong University, 266237 Qingdao, P*.*R. China*

**Le-Le Zhu** *− Fetal Medicine Center, Qingdao Women and Children’s Hospital, Qingdao University, 266071 Qingdao, P*.*R. China*

**Jia-Qi Hu** *− Fetal Medicine Center, Qingdao Women and Children’s Hospital, Qingdao University, 266071 Qingdao, P*.*R. China*

**Chao-Yi Wang** *− Fetal Medicine Center, Qingdao Women and Children’s Hospital, Qingdao University, 266071 Qingdao, P*.*R. China*

**Rui-Juan Li** *− Fetal Medicine Center, Qingdao Women and Children’s Hospital, Qingdao University, 266071 Qingdao, P*.*R. China*

## Author Contributions

^§^ L.N., W.H contributed equally to this work.

### Notes

The authors declare no competing financial interest.

## ACKNOWLEDGMENT

We would like to thank Haiyan Sui, Jing Zhu, and Zhifeng Li of the Core Facilities for Life and Environmental Sciences, State Key laboratory of Microbial Technology of Shandong University for NMR and LC−MS measurements. We gratefully acknowledge Dr. Wen-Xuan Wang (Xiangya College of Pharmacy, Central South University) for ECD calculations. This work was financially supported by the National Key Research and Development Programs of China (no. 2021YFC2101000, 2019YFA0905700, 2018YFA0900400, and 2018YFA0901704), the National Natural Science Foundation of China (NSFC) (no. 32222003, 31900042, 81973215, 31670076, and 31471183), and Excellent Youth Program of Shandong Natural Science Foundation (no. ZR2020YQ62).

