## Supplemental Data 1 for "Genomics-Driven Discovery of Myxopyromides, a Class of Polyketidic Amides Associated with Predation of *Myxococcus* sp. SDU36"

**Table S1.** Annotations for *mpd* biosynthetic gene cluster (BGC).

| ORF | Gene size (bp) | Gene ID | Proposed function | NCBI similarity | Ident% |
|---|---|---|---|---|---|
| 1 | 999 | *mpdF* | DNA-binding transcriptional regulator | WP_163780248.1\| LysR substrate-binding domain-containing protein [Myxococcus vastator] | 98.2% |
| 2 | 906 | *mpdG* | beta-lactamase | WP_176416362.1\| MBL fold metallo-hydrolase [unclassified Myxococcus] | 99.0% |
| 3 | 330 | *mpdH* | DNA-binding transcriptional regulator | NVJ08004.1\| PadR family transcriptional regulator [Myxococcus sp. AM001] | 99.1% |
| 4 | 360 | *mpdI* | | WP_163780240.1\| DUF1048 domain-containing protein [Myxococcus vastator] | 100.0% |
| 5 | 1254 | *mpdJ* | D-alanyl-D-alanine carboxypeptidase | NVJ08006.1\| beta-lactamase family protein [Myxococcus sp. AM001] | 100.0% |
| 6 | 6150 | *mpdE* | KS-AT-KR-DH-ACP-TE | NVJ08007.1\| SDR family NAD(P)-dependent oxidoreductase [Myxococcus sp. AM001]; WP_141324826.1\| type I polyketide synthase [Myxococcus sp. AB025B] | 96.96%; 85.89% |
| 7 | 3330 | *mpdD* | C-A-PCP | WP_163785084.1\| amino acid adenylation domain-containing protein [Myxococcus vastator] | 98.2% |
| 8 | 8025 | *mpdA* | ACP-KS-AT-AT-DH-ER-KR-ACP | WP_074958388.1\| type I polyketide synthase [Myxococcus fulvus] | 100.0% |
| 9 | 10083 | *mpdB* | KS-AT-KR-ACP-KS-AT-DH-KR-ACP | WP_074958389.1\| type I polyketide synthase [Myxococcus fulvus] | 87.7% |
| 10 | 4197 | *mpdC* | KS-AT-oMT-ACP | AKF80292.1\| polyketide synthase [Myxococcus fulvus 124B02] | 91.2% |
| 12 | 690 | *mpdK* | | WP_176421047.1\| DUF2071 domain-containing protein [unclassified Myxococcus] | 100.0% |
| 13 | 969 | *mpdL* | Serine aminopeptidase; Lysophospholipase | NVJ09646.1\| alpha/beta fold hydrolase [Myxococcus sp. AM001] | 98.8% |
| 14 | 1839 | *mpdM* | acyl-CoA synthetase | WP_176421045.1\| AMP-binding protein [Myxococcus sp. AM010] | 95.6% |
| 15 | 807 | *mpdN* | Enoyl-CoA hydratase/carnithine racemase | NVJ09648.1\| enoyl-CoA hydratase/isomerase family protein [Myxococcus sp. AM001] | 100.0% |
| 16 | 924 | *mpdO* | DNA-binding transcriptional regulator | NVJ09649.1\| LysR family transcriptional regulator [Myxococcus sp. AM001] | 100.0% |
| 17 | 594 | *mpdP* | Peroxiredoxin (PRX)-like | WP_217918795.1\| redoxin domain-containing protein [Myxococcus sp. AM010] | 93.9% |
| 18 | 471 | *mpdQ* | 2-oxoglutarate dehydrogenase C-terminal | WP_120585737.1\| PspA [Corallococcus] | 69.7% |

**Table S2.** Half-inhibitory concentrations ($IC_{50}$ in μM) for myxopyromides. The compound docetaxel (doc) is the positive control.

| Cell line | **1** | **3** | **4** | Doc |
|---|---|---|---|---|
| Hela (human cervical carcinoma cells) | 32.12 | 23.75 | 24.98 | 15.42 |
| PC-3 (human prostatic cancer cells) | 39.8 | 38.35 | 35.8 | 18.54 |
| MCF-7 (human breast cancer cells) | 47.59 | 47.22 | 41.02 | 19.79 |

**Table S3.** Strains and plasmids used in the study.

| **Strains** | **Features** | **Source** |
|---|---|---|
| *Myxococcus* sp. SDU36 | Wild-type strain | This study |
| SDU36-$P_{mpd}$ | Activation of *mpd* cluster in SDU36 | This study |
| *E. coli* Top10 | $F^-$ *mcrA* Δ(*mrr-hsdRMS-mcrBC*) Φ80 *lac*ZΔM15 Δ*lac*X74 *recA1* *ara*Δ139 Δ(*ara-leu*)7697 *galU* *galK* *rps,* L($Str^R$) *endA1* *nup*G | Laboratory collection |
| **Plasmids** | **Parts** | **Function** |
| pBa-1 | p15A ori, *kan*$^r$, BBa_J23104-BBa_B0034-eGFP | Basic vector for activation *in vivo*. |
| pBa-*mpd* | p15A ori, *kan*$^r$, BBa_J23104-BBa_B0034-homologous arm (the starting 1026 bp of *mpdA*) of *mpd* cluster | Activation of *mpd* cluster |

**Table S4.** List of primers used in this study.

| Primers | Sequence (5'-3') | Application |
|---|---|---|
| cpBa-F | TGGATGAGCTGTACAAGTGATAATTATCAGAAGAACTCGTCAAGAA GGCG | Linearization of pBa-1 |
| cpBa-R | CACCATCTAGTATTTCTCCTCTTTCGCTAGAG | |
| Ampd-F | AGGAGAAATACTAGATGGTGTCGATGGATACAGGGGGGCAGGAG | homologous arm for activating *mpd* |
| Ampd-R | TCACTTGTACAGCTCATCCAGCGCGGCGACCTCGATGG | |
| part3-F | TCTTCCGCTTCCTCGCTCAC | Clone fragment containing parts ColE1 ori and *galK* |
| part2-R | GCGGGACTCTGGGGTTCG | |
| Cm-F | CTCACCCAGAAACGCTGGTGCCGGATCTGCAGCTTAAGGAAC | Clone fragment of chloramphenicol resistance |
| Cm-R(O)part3 | GTGAGCGAGGAAGCGGAAGAGCCAGTCATTAGGCCTATCTGAC | |
| pBb-TF | CCCAACCTTTCATAGAAGGCGG | Sequencing for the homologous arm |
| pBb-TR | GTTCCTTAAGCTGCAGATCCGG | |
| mpdA-F | TTCGAACCCCAGAGTCCCGCGCTTCATCACCCCCGAAGAGG | Amplification of *mpdA* homologous arm |
| mpdA-R | CACCAGCGTTTCTGGGTGAGTGCTCCGCGAAGACGAGC | |
| mpdB-F | TTCGAACCCCAGAGTCCCGCGCCCGTCTTCCAGGTGATGTTCAA | Amplification of *mpdB* homologous arm |
| mpdB -R | CACCAGCGTTTCTGGGTGAGCGCAGCAGCCGGATTCGCCGCCCC | |
| mpdC-F | TTCGAACCCCAGAGTCCCGCCTTCAGGTGCTCCTGGAGGC | Amplification of *mpdC* homologous arm |
| mpdC-R | CACCAGCGTTTCTGGGTGAGACGCCGGGCGTGGGAAT | |
| mpdD-F | TTCGAACCCCAGAGTCCCGCATTCCCCTCGTGTCGAACGTGA | Amplification of *mpdD* homologous arm |
| mpdD-R | CACCAGCGTTTCTGGGTGAGTCACCATCCACCGCGCGA | |
| T28-F (36) | AAGATCCGTTCAAAAGCGAATAGTCC | Genotype confirmation of SDU36-*mpd* |
| T28-R (36) | CGCGTCCCGGAGTCGAA | |
| GapA-SDU36Q-F | AGAAGGTCATCATCTCCGCC | Transcription verification of *gapA* |
| GapA-SDU36Q-R | GTCATCAGGCCCTTCTCGAT | |
| QmpdA-F | ACCTCTATACACACCTGGCG | Transcription verification of *mpdA* (on the homologous arm of pBa-*mpd*) |
| QmpdA-R | GTCCGACTCGAAGCTCAAGA | |
| QmpdA-F2 | GGGGAAGATGCTCTCGGTG | Transcription verification of *mpdA* |
| QmpdA-R2 | AGGGCGTCATCCACCATG | |
| QmpdB-F | CCTGGCCTACGTCATCTACA | Transcription verification of *mpdB* |
| QmpdB-R | AGATGGAGAGGTCGAAGCAG | |
| QmpdC-F4 | AACTGGTGCTGATGTCCGAG | Transcription verification of *mpdC* |
| QmpdC-R4 | CCGCCATCTCCTCGTAGTG | |
| QmpdD-F | TCGACGCCTACTACATCACC | Transcription verification of *mpdD* |
| QmpdD-R | CAATCATCTCCGTGCTCAGC | |
| Qc7-F | AGAGCGAGCGTTCCTACTAC | Transcription verification of BGC 7 |
| Qc7-R | TTGCAGCATCAGATCCAACG | |
| Qc20-F | CGTTCCTCTACATGCAGCAG | Transcription verification of BGC 20 |
| Qc20-F | AGTCGAGGTCACTGTGGAAG | |

**Figure S1.** Diagrammatic sketch for genetic manipulation using single crossover recombination. The plasmid pBa-1 equipped with a p15A origin of replication (*ori*) and kanamycin-resistant (*km*$^R$) selection marker was used for exchanging the native promotor of *mpdE*. The relative positions of the primers for the PCR verification of promotor engineering and insertion disruption were indicated.

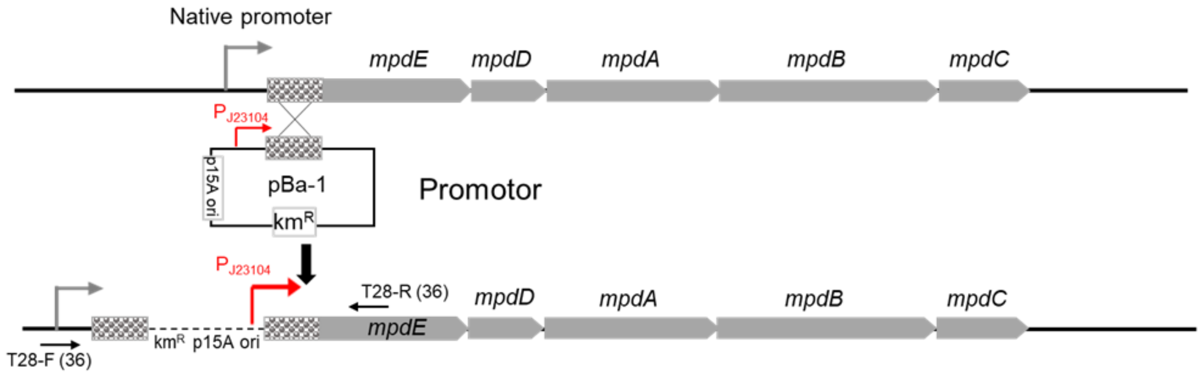


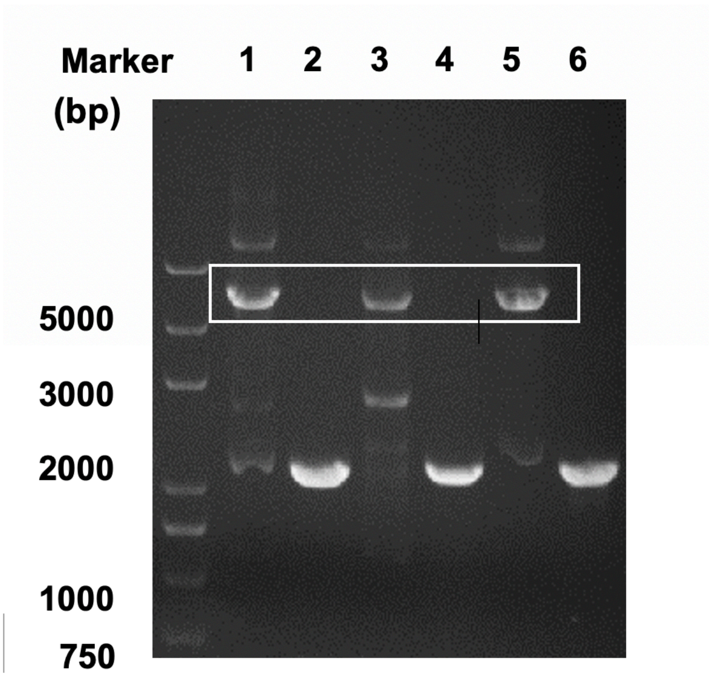

**Figure S2.** Activation and transcriptional determination of *mpd*. (A) Promoter replacement to activate *mpd* cluster. Promoter J23104 was inserted afront of the *mpd*E gene. (B) Transcription of the *mpd* cluster before and after promoter replacement. Reverse transcription (RT-PCR) was used to semi-quantitatively detect the up-promoted *mpd*. Lanes S, Samples, cDNA as the template; Lanes +, genomic DNA as the template for positive controls; Lanes -, RNA as the template to prove the removal of genomic DNA; M, Trans 2K markers.

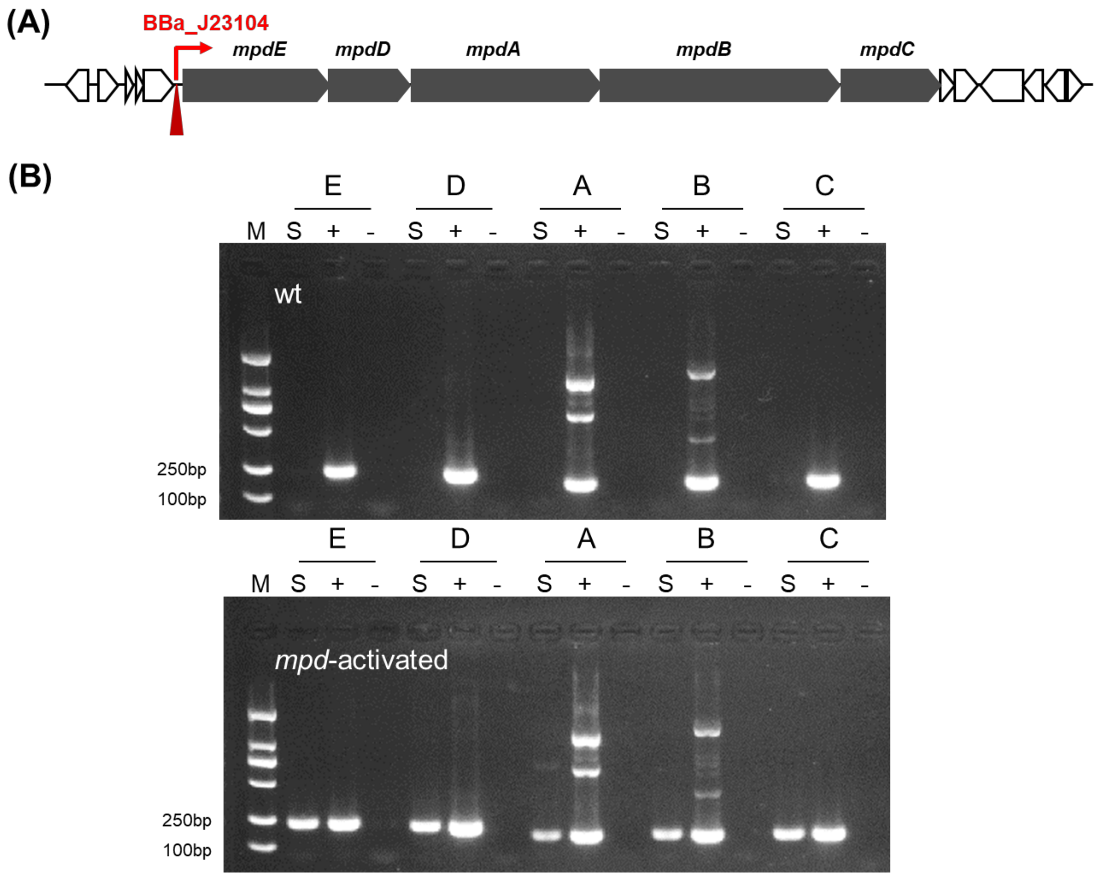

**Figure S3.** NOESY ( **━** ) correlations of myxopyromides **1**–**5**.

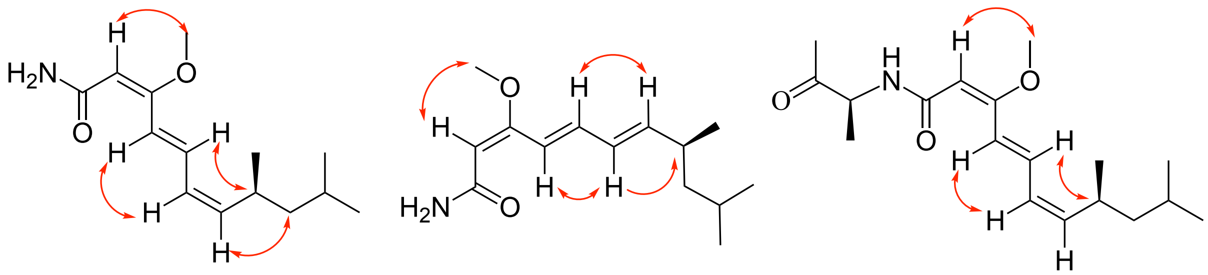

**1** **2** **3**

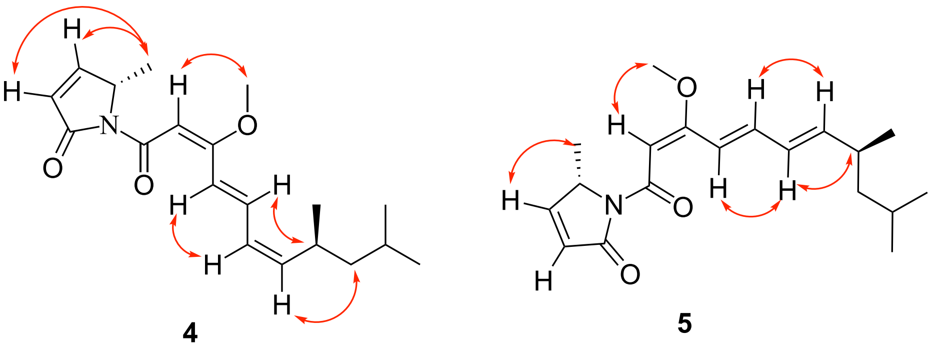

**4** **5**

**Figure S4**. The 330 TE sequences used to construct the phylogenetic tree were obtained from the reference (Horsman et al., 2016), including 38 HMM seed sequences and the rest from the ClusterMine360 database (Conway and Boddy, 2013). The TE domain of *mpd* in strain SDU36 and the other 9 equivalent TE domains from myxobacteria were also involved. MEGA11 software was used to analyze molecular evolution by Maximum Likelihood after sequence alignment by ClustalW method, and all parameters remained default. iTol was used to present the unrooted tree. The TEs of *mpd* clusters from myxobacteria were labeled red. FrsATE was grouped closest to FrsGTE, indicating an evolutionary relationship of the two TEs from the same BGC.

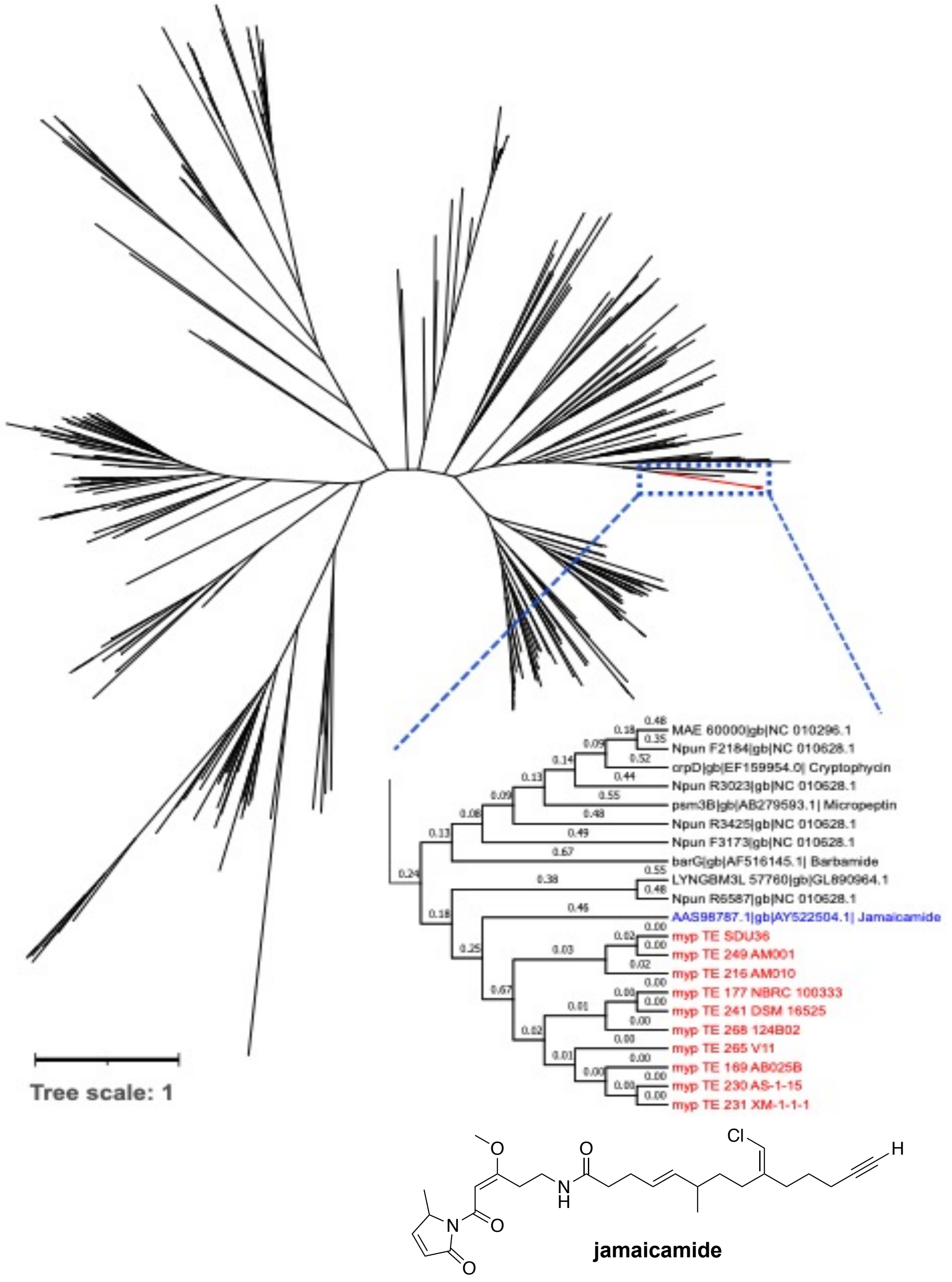

**Figure S5.** Bioinformatics-based configurational assignment of C-8 of myxopyromides by sequence alignment of ER domains. The crucial amino acid residues for the prediction of stereochemistry of methyl-bearing stereocenters were box marked, wherein the crucial tyrosine (Y) is predictive of *S* configuration, and the valine (V) determines *R* configuration. The reference sequences were trimmed from the characterized BGCs specifying known compounds. *S*-type: *eryAII* (Erythromycin), *megAII* (Megalomicin), *lkmAII* (Lankamycin), *rapB* (Rapamycin), *mycAIII* (Mycinamicin), *chmGIII* (Chalcomycin), *tylG* (Tylosin), *pikAII* (Pikromycin), *nanA2* (Nanchangmycin), *monAII* (Monensin), nigAIII (Nigericin), *gdmA1* (Geldanamycin), *tmcA* (Tautomycetin), *merB* (Meridamycin); *R*-type: *rapA* (Rapamycin), *rapC* (Rapamycin), *borA5* (Borrelidin), *olmA3* (Oligomycin), *olmA4* 1(Oligomycin), *nigAIV* (Nigericin), *nigAVIII* (Nigericin), *gdmA1* (Geldanamycin), *tmcB* (Tautomycetin). MUSCLE in MEGA11 was used for alignment with the default parameter. The alignment result was displayed by ESPript 3.0 in B&W color scheme.

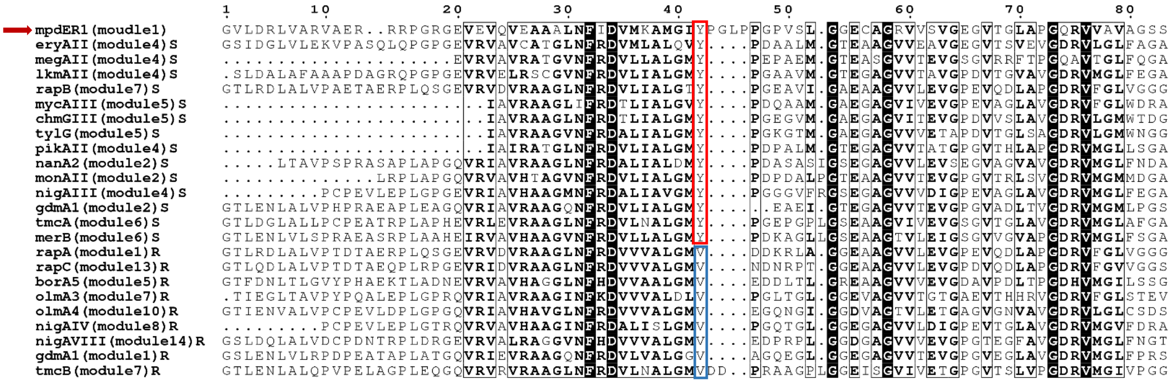

**Figure S6.** Bioinformatics-based configurational assignment of double bonds of myxopyromides by sequence alignment of KR domains. The residue 155 “W” in red frame was responsible for the A-type reduction that leads to *cis* double bonds. The conserved residues “LDD” and “PN” in blue box were the key motifs for B-type reduction that causes the generation of *trans* double bonds. The reference sequences were trimmed from the characterized BGCs specifying known compounds. The referred BGCs are MYX (myxalamid), EPO (epothilone), DEBS (erythromycin), NYS (nystatin). MUSCLE in MEGA11 was used for alignment with the default parameter. The alignment result was displayed by ESPript 3.0 in B&W color scheme.

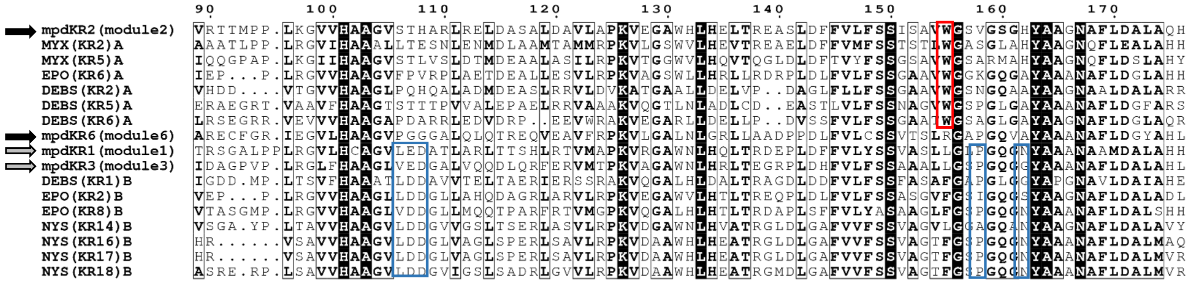

**Figure S7.** $^1$H NMR (600 MHz, $CD_3OD$) spectrum of myxopyromide A (**1**).

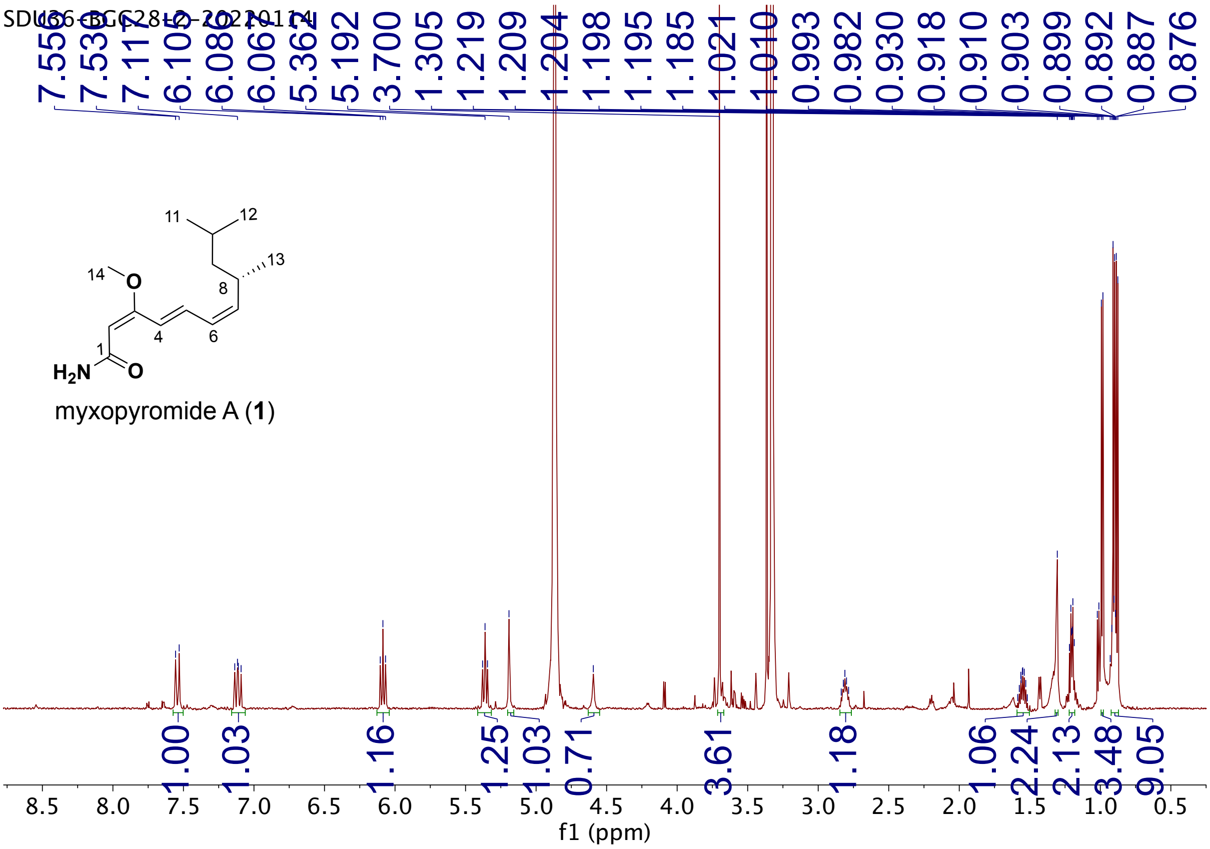

**Figure S8.** $^{13}C$ NMR (600 MHz, $CD_3OD$) spectrum of myxopyromide A (**1**).

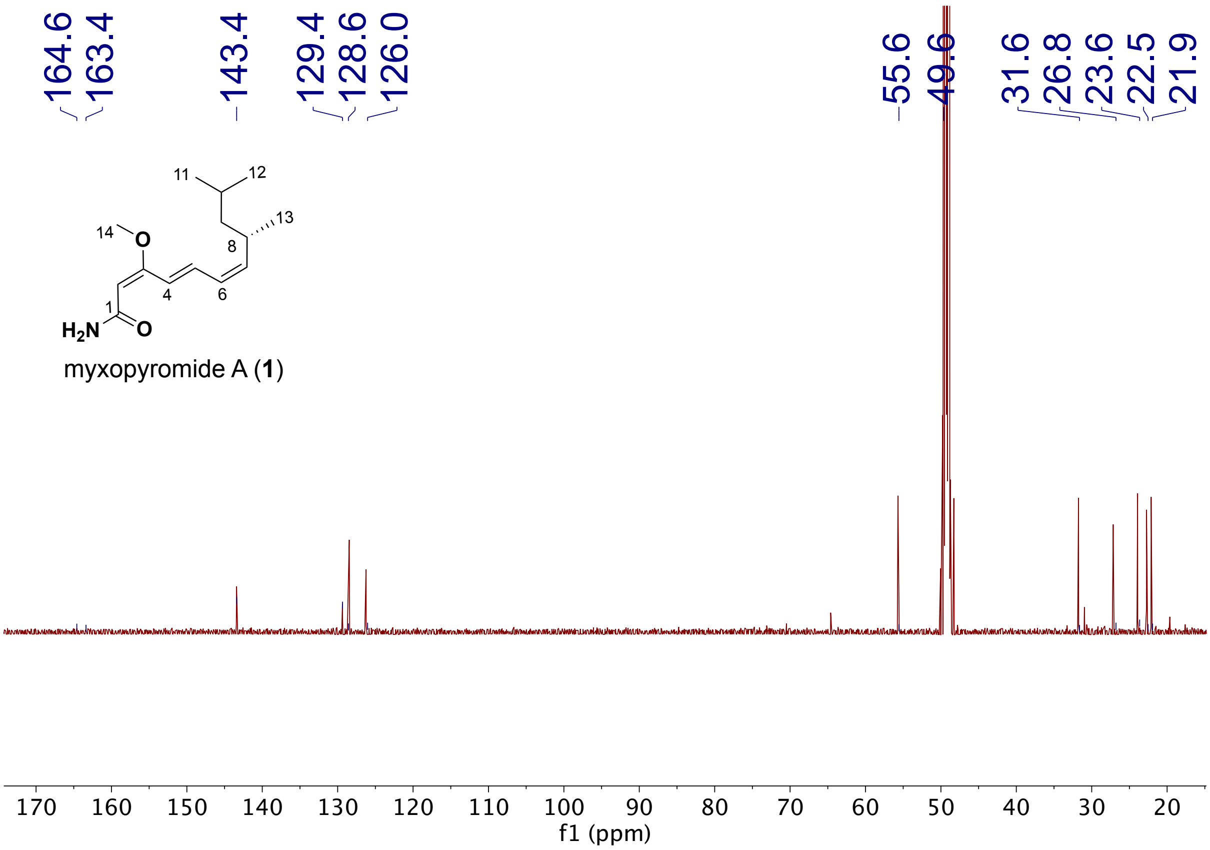

**Figure S9.** $^{1}H$-$^{13}C$ HSQC NMR (600 MHz, $CD_3OD$) spectrum of myxopyromide A (**1**).

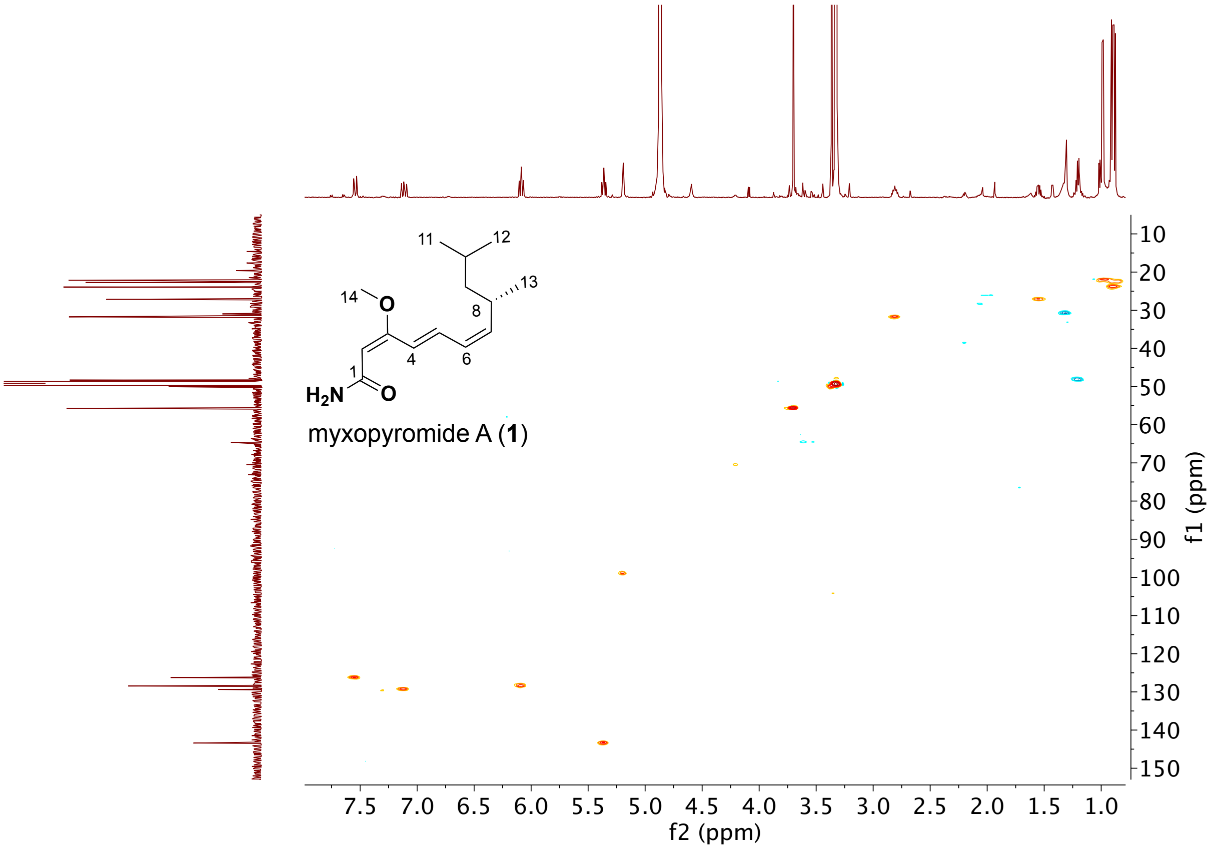

**Figure S10.** $^{1}$H-$^{13}$C HMBC NMR (600 MHz, $CD_3OD$) spectrum of myxopyromide A (**1**).

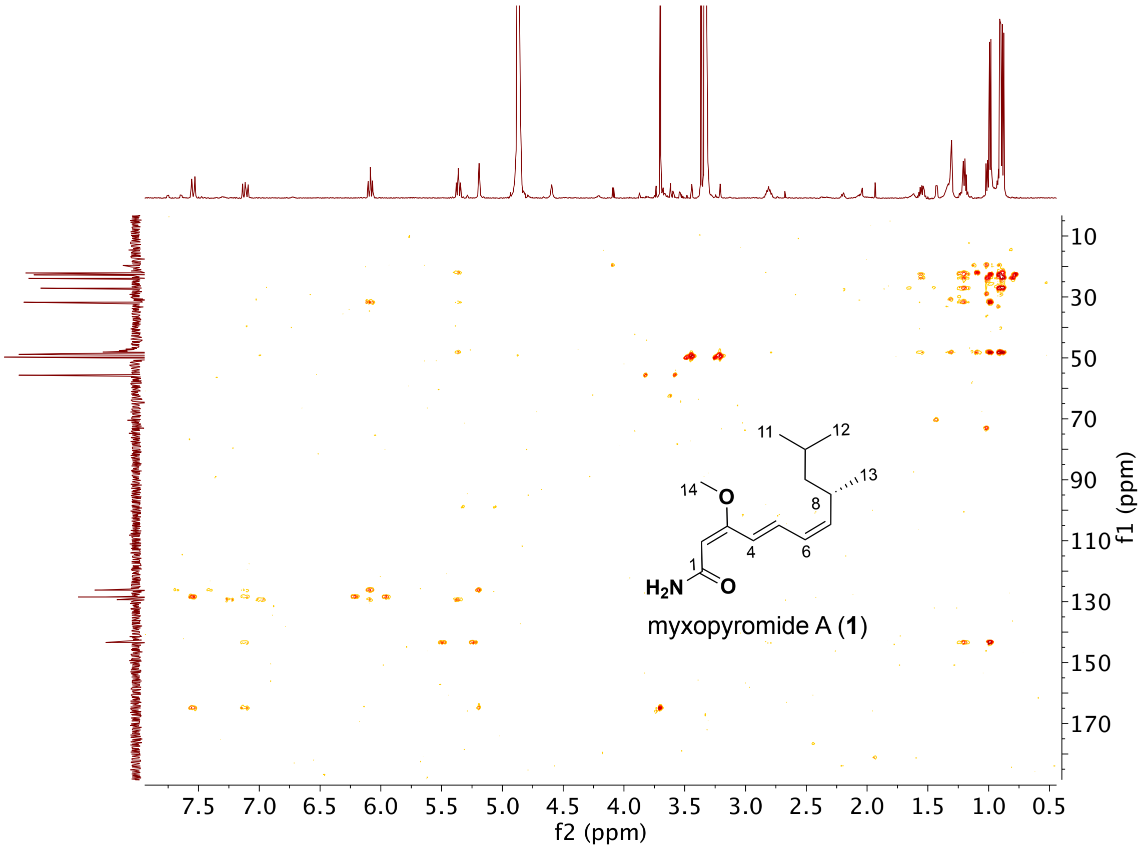

**Figure S11.** [1]H-[1]H COSY NMR (600 MHz, $CD_3OD$) spectrum of myxopyromide A (**1**).

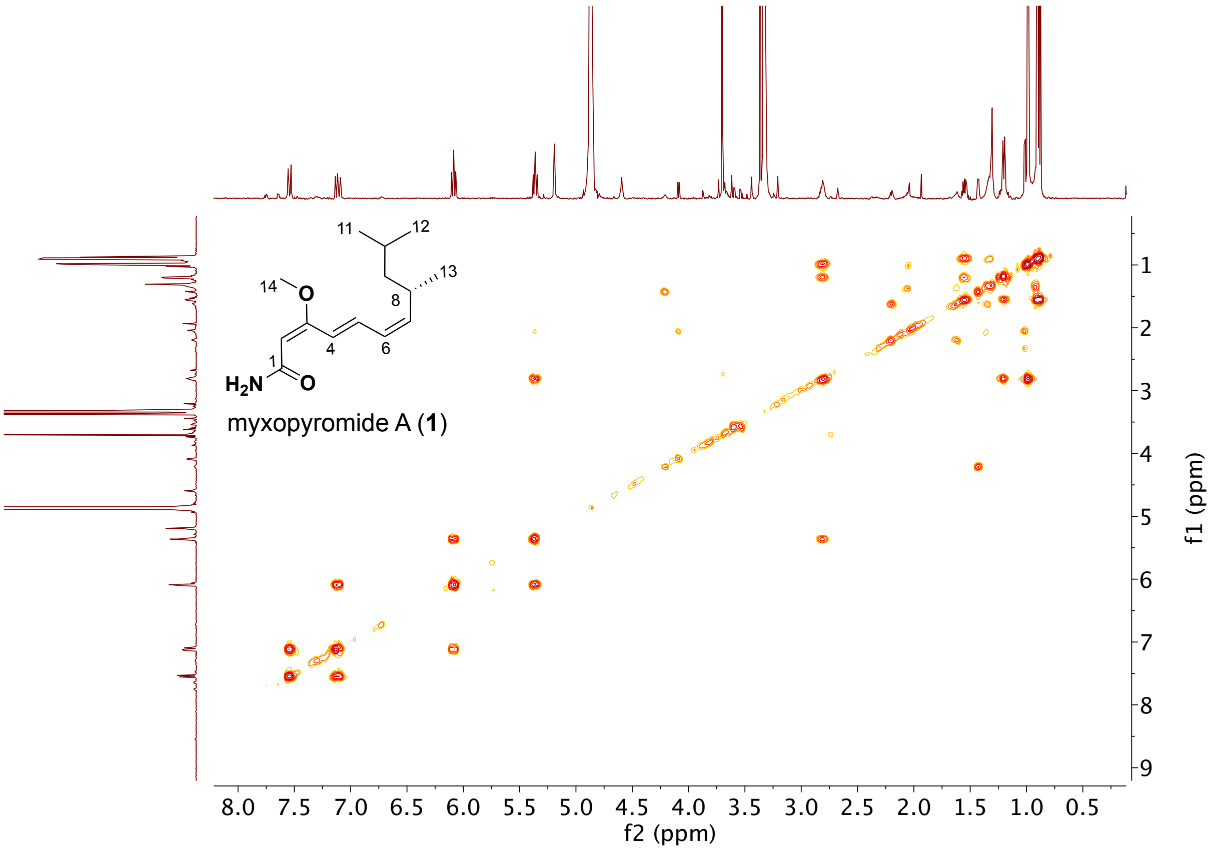

**Figure S12.** NOESY NMR (600 MHz, $CD_3OD$) spectrum of myxopyromide A (**1**).

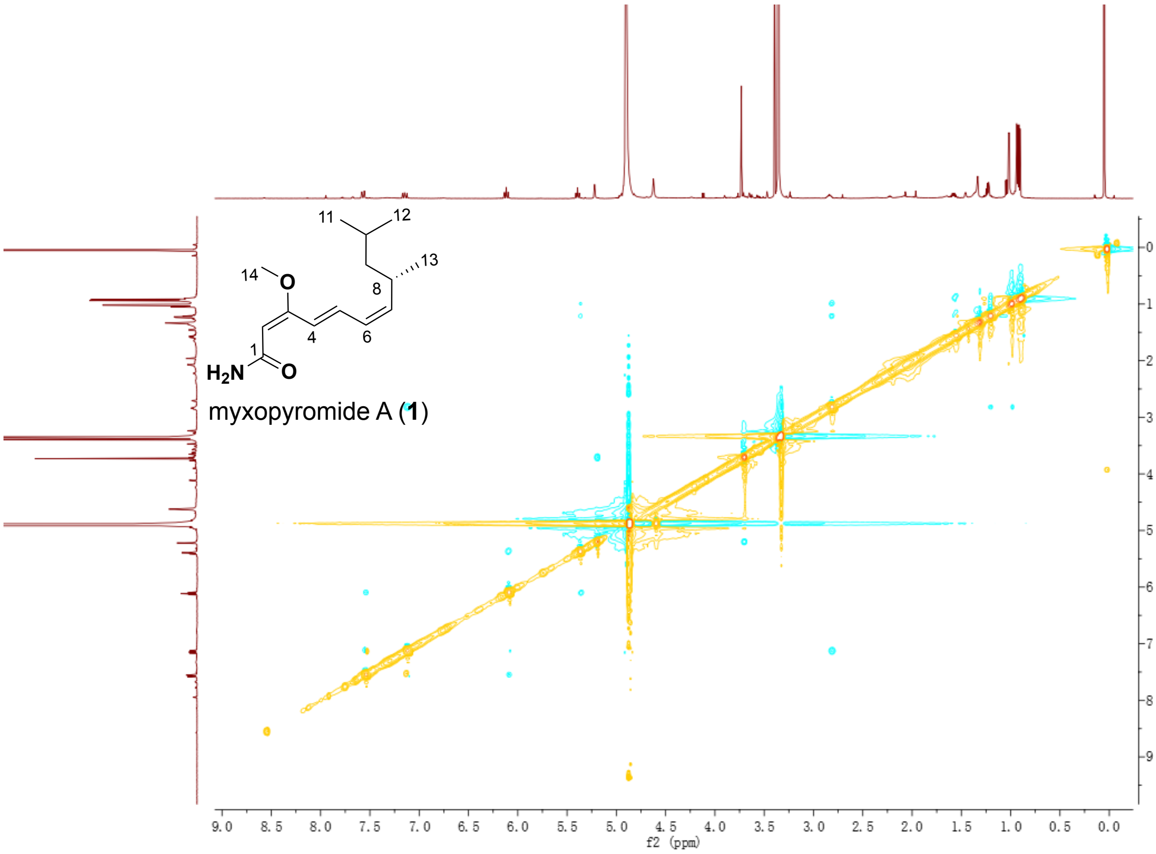

**Figure S13.** High-resolution mass spectrum of myxopyromide A (**1**).

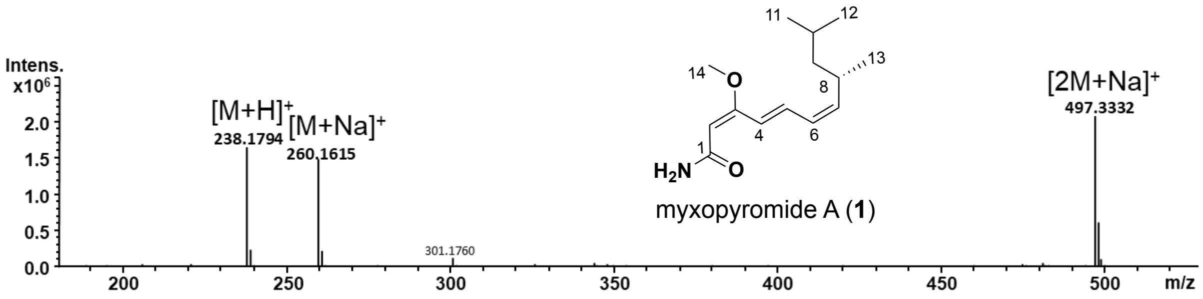


**Figure S14.** UV spectrum of myxopyromide A (**1**).

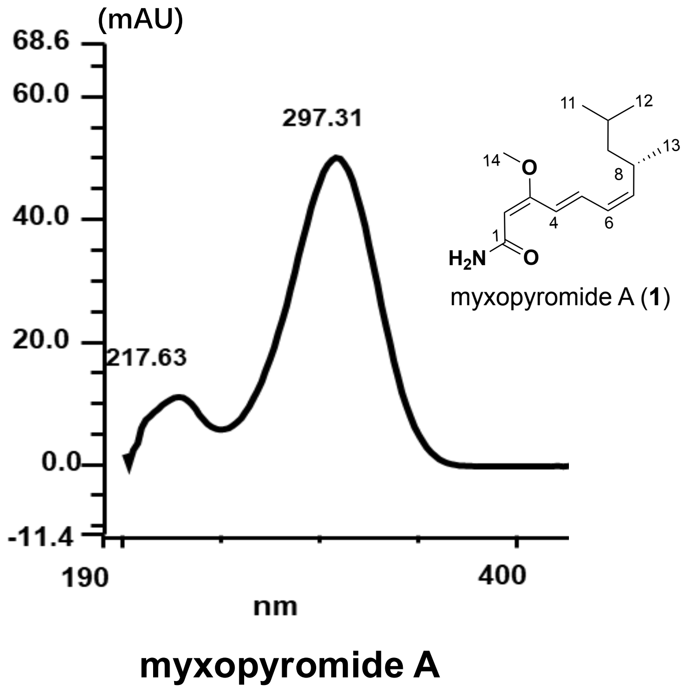

**Figure S15.** IR spectrum of myxopyromide A (**1**).

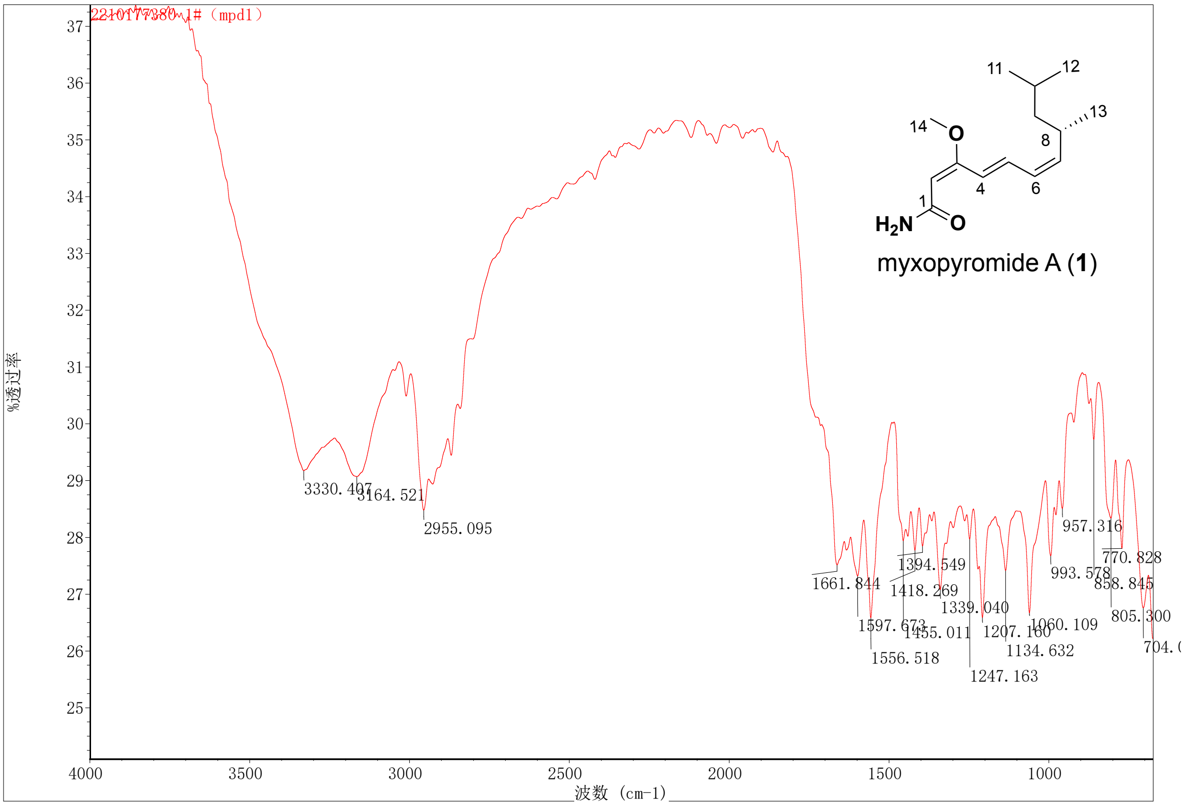

**Figure S16.** $^1$H NMR (600 MHz, $CDCl_3$) spectrum of myxopyromide B (**2**).

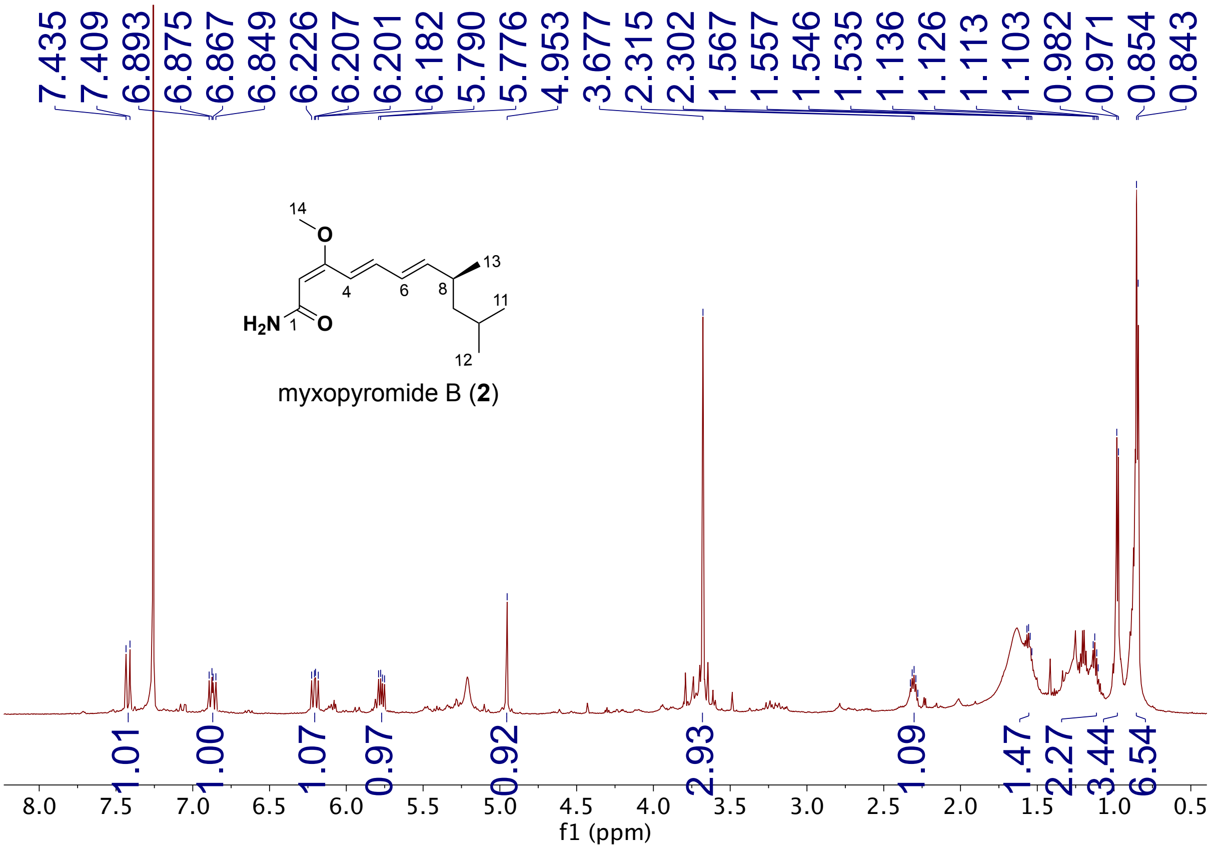

**Figure S17.** $^{13}C$ NMR (600 MHz, $CDCl_3$) spectrum of myxopyromide B (**2**).

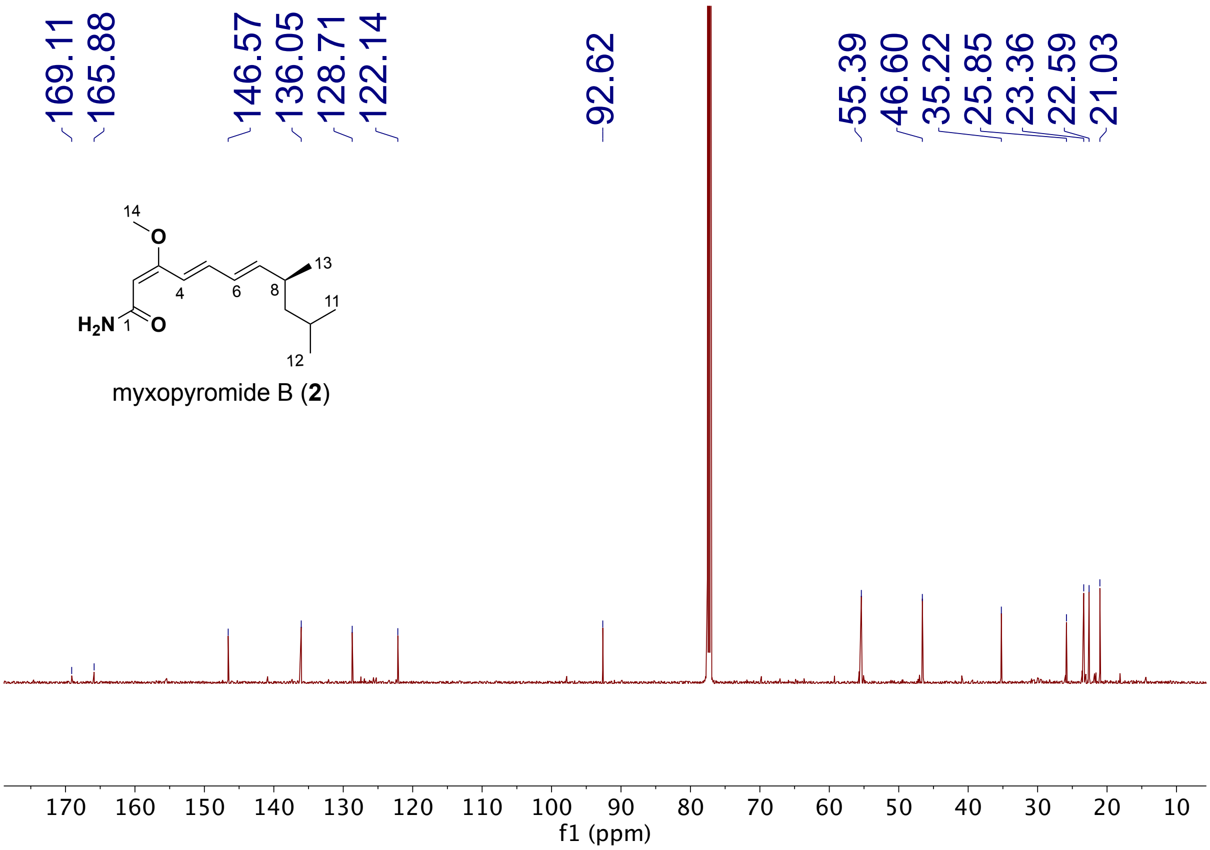

**Figure S18.** [1]H-[13]C HSQC NMR (600 MHz, $CDCl_3$) spectrum of myxopyromide B (**2**).

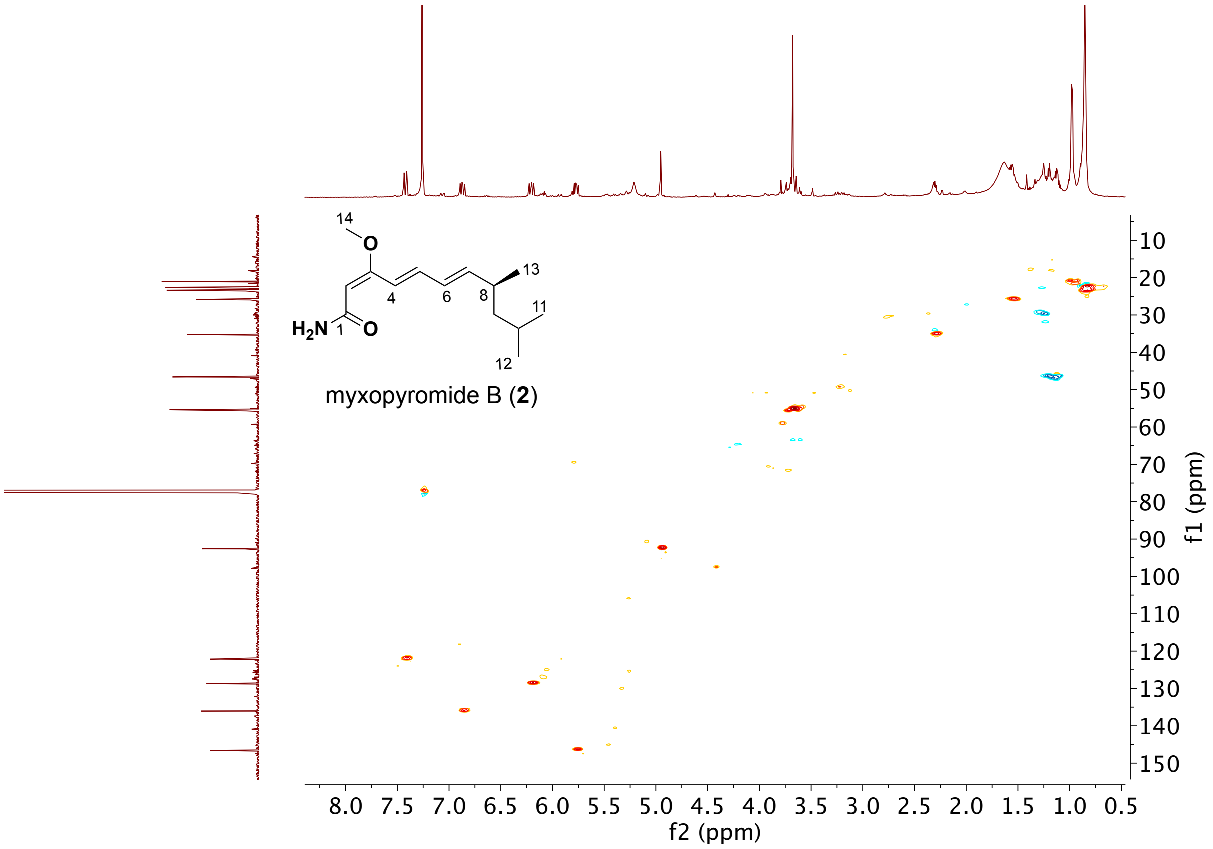

**Figure S19.** $^{1}H$-$^{13}C$ HMBC NMR (600 MHz, $CDCl_3$) spectrum of myxopyromide B (**2**).

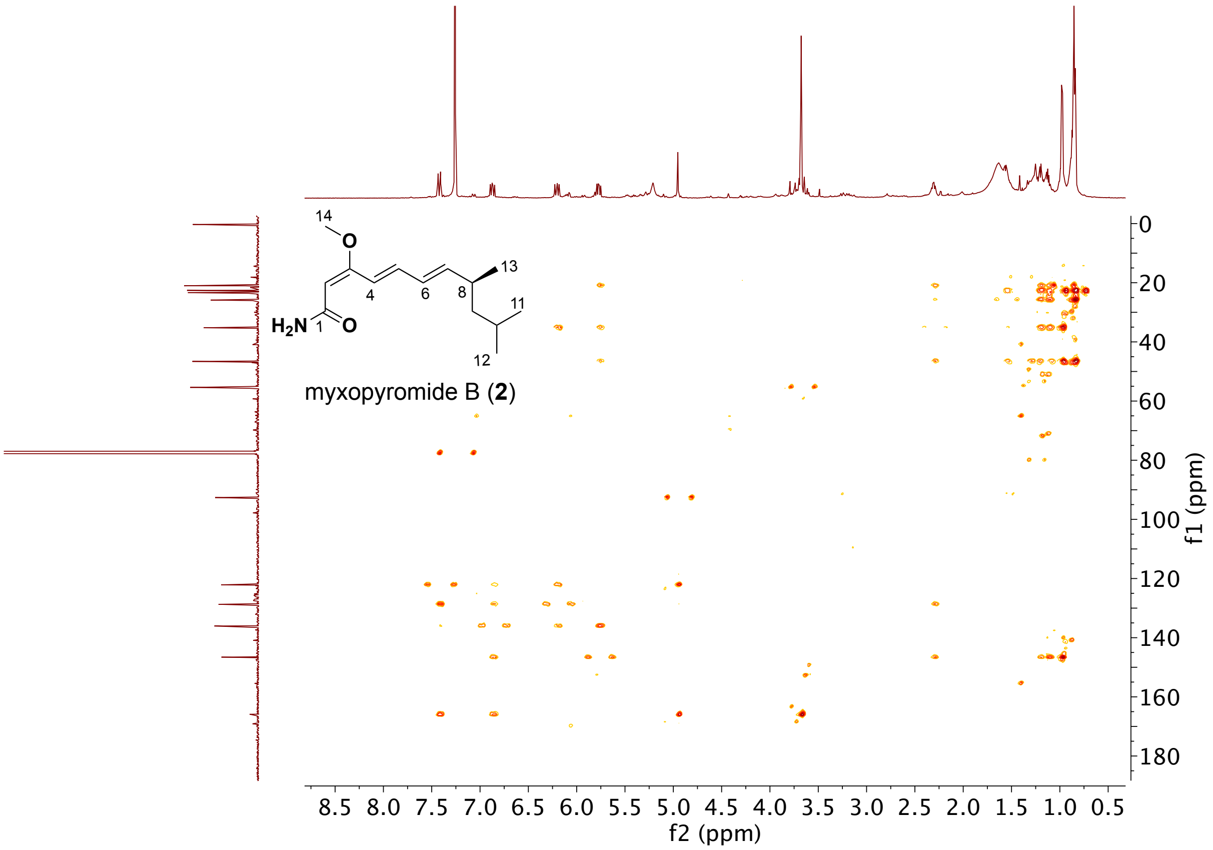

**Figure S20.** $^1H$-$^1H$ COSY NMR (600 MHz, $CDCl_3$) spectrum of myxopyromide B (**2**).

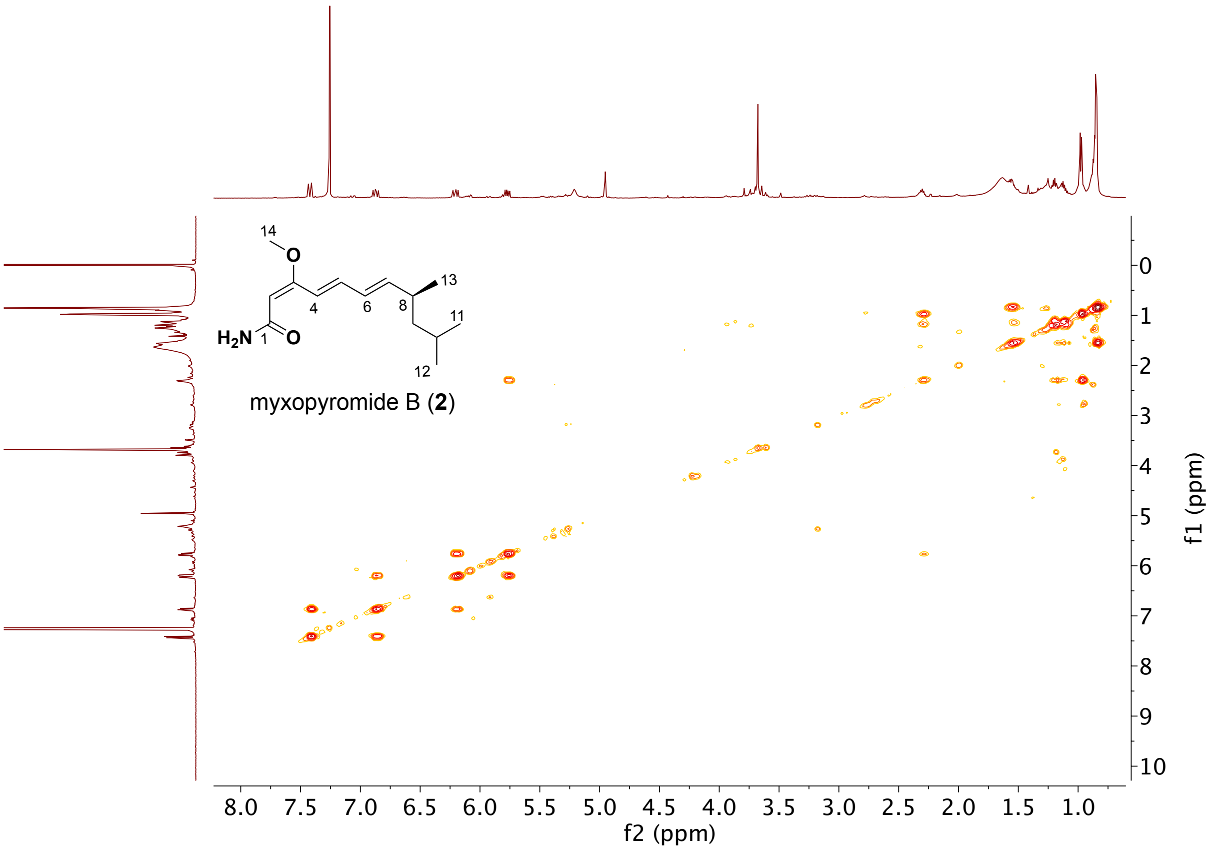

**Figure S21.** NOESY NMR (600 MHz, $CDCl_3$) spectrum of myxopyromide B (**2**).

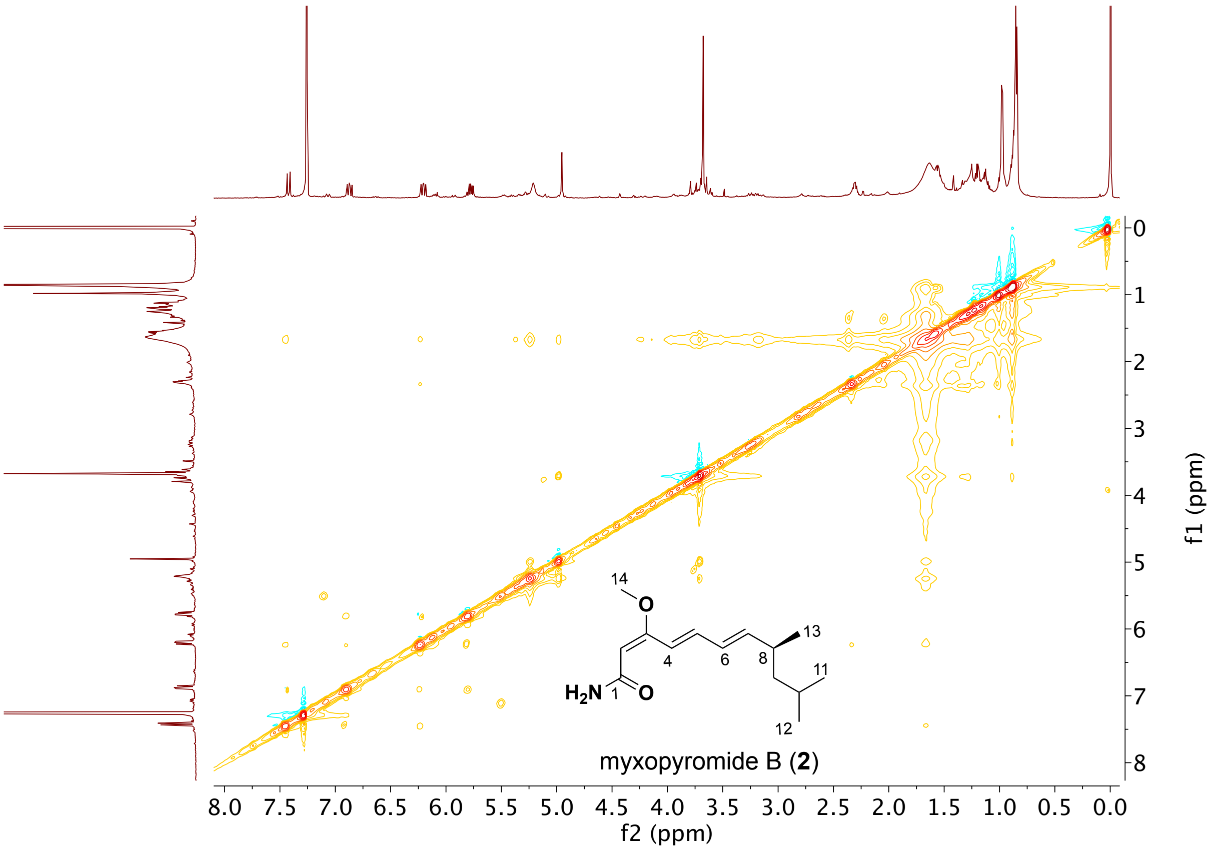

**Figure S22.** High-resolution mass spectrum of myxopyromide B (**2**).

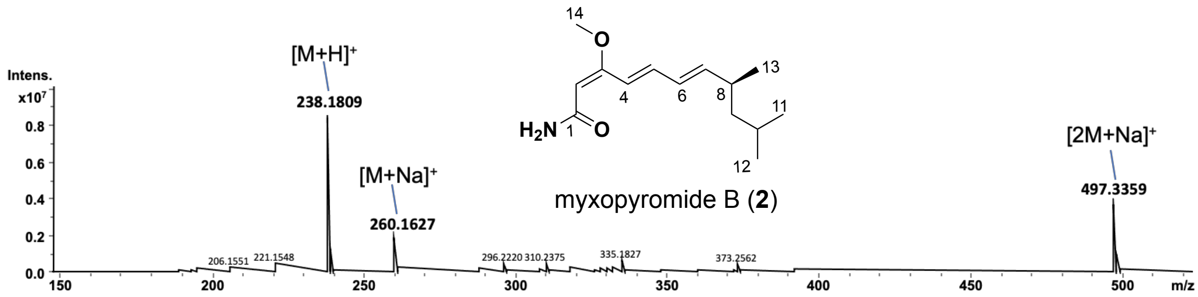


**Figure S23.** UV spectrum of myxopyromide B (**2**).

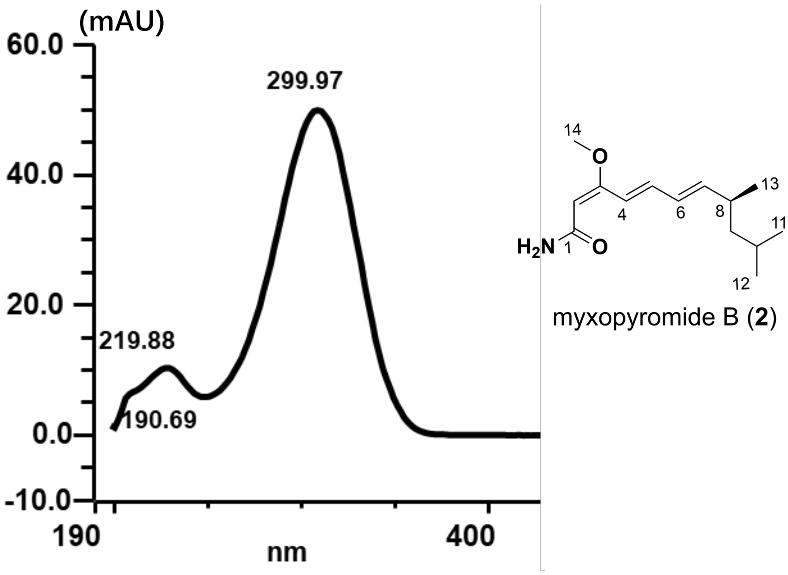

**Figure S24.** UV spectrum of myxopyromide B (**2**).

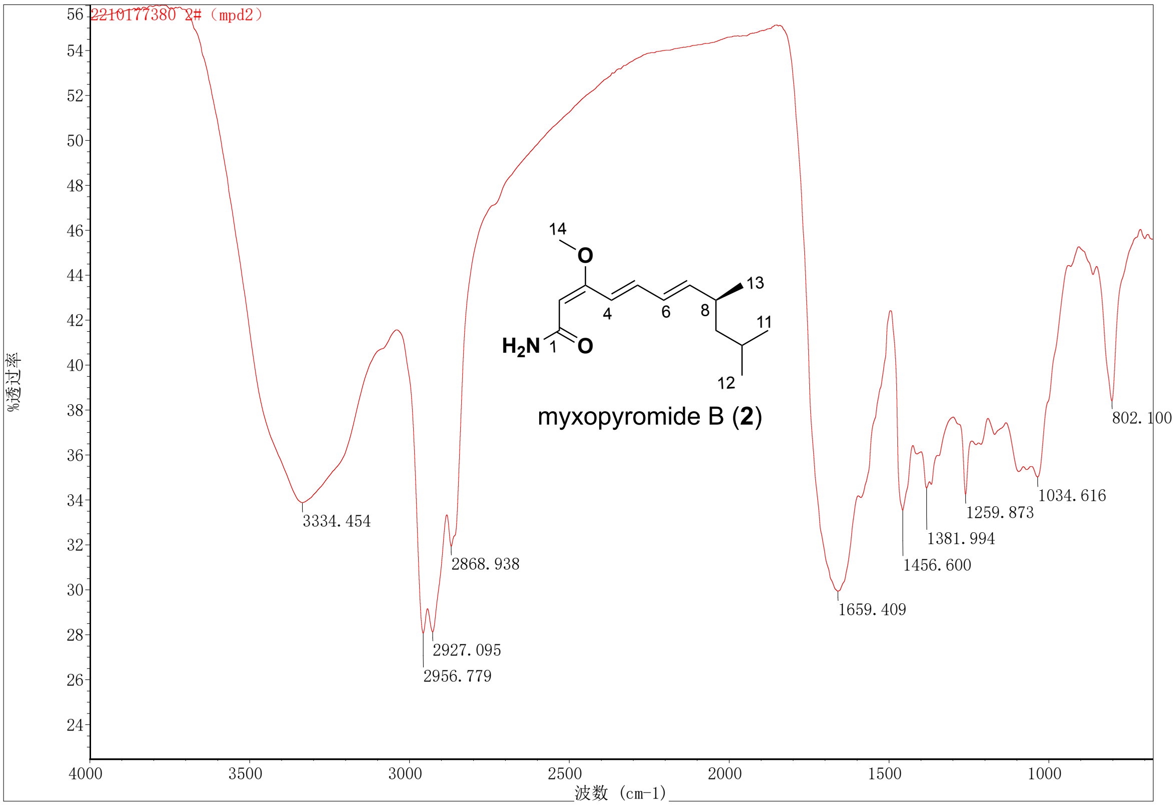

**Figure S25.** $^1$H NMR (600 MHz, $CD_3OD$) spectrum of myxopyromide C (**3**).

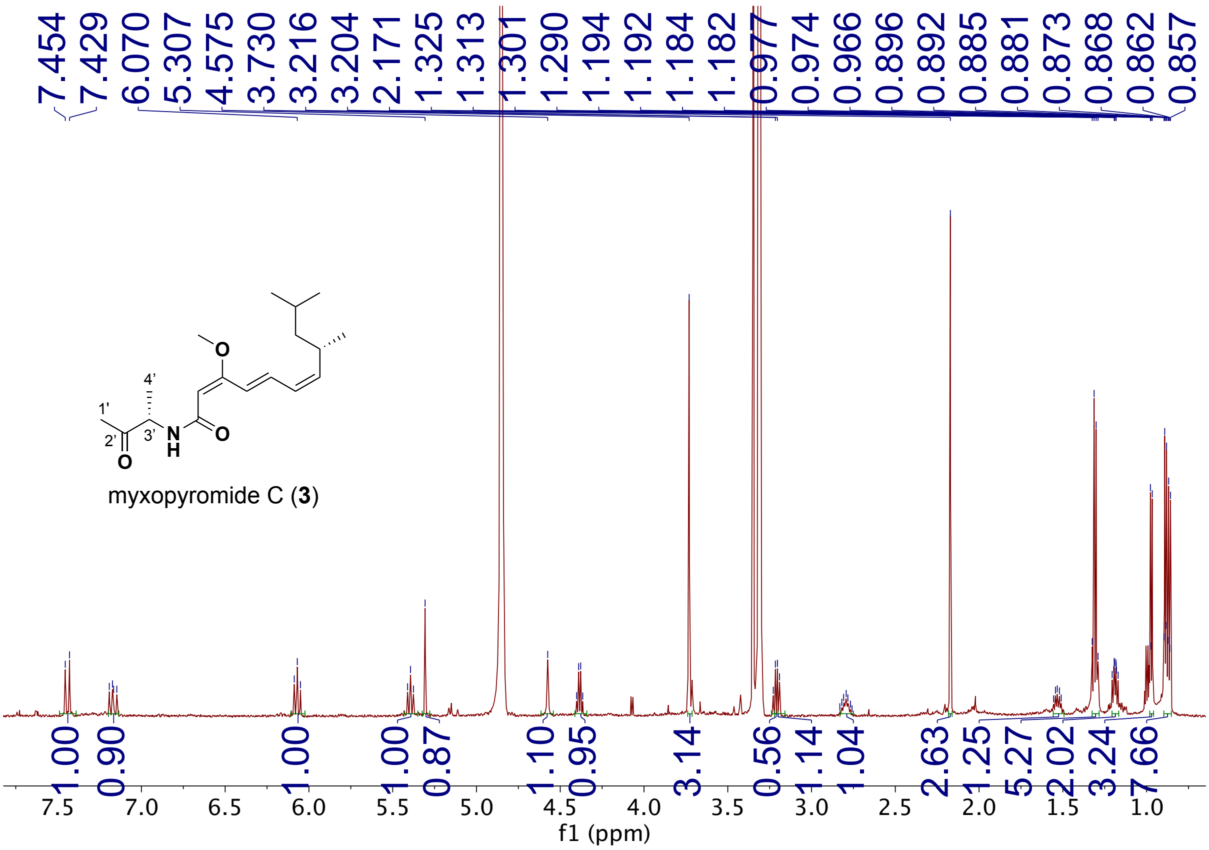

**Figure S26.** $^{13}C$ NMR (600 MHz, $CD_3OD$) spectrum of myxopyromide C (**3**).

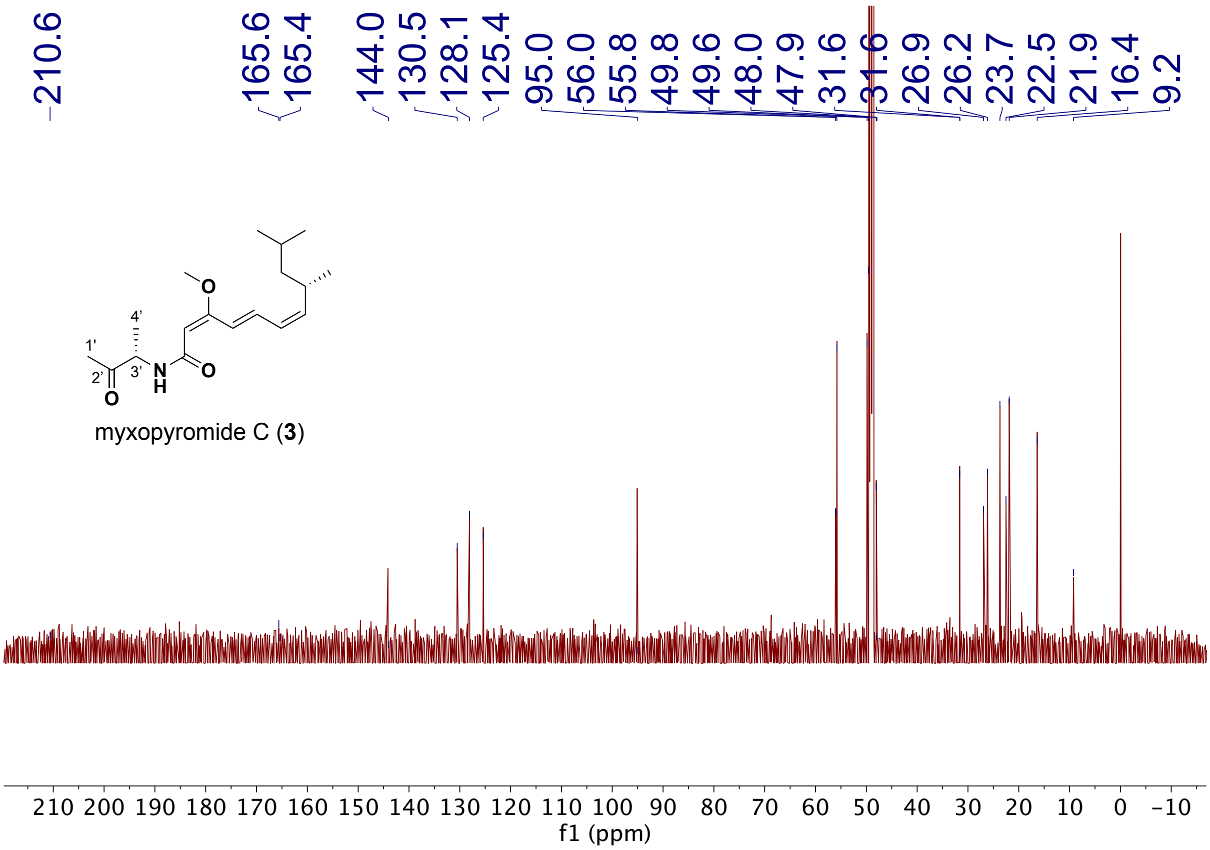

**Figure S27.** $^{1}H$-$^{13}C$ HSQC NMR (600 MHz, $CD_3OD$) spectrum of myxopyromide C (**3**).

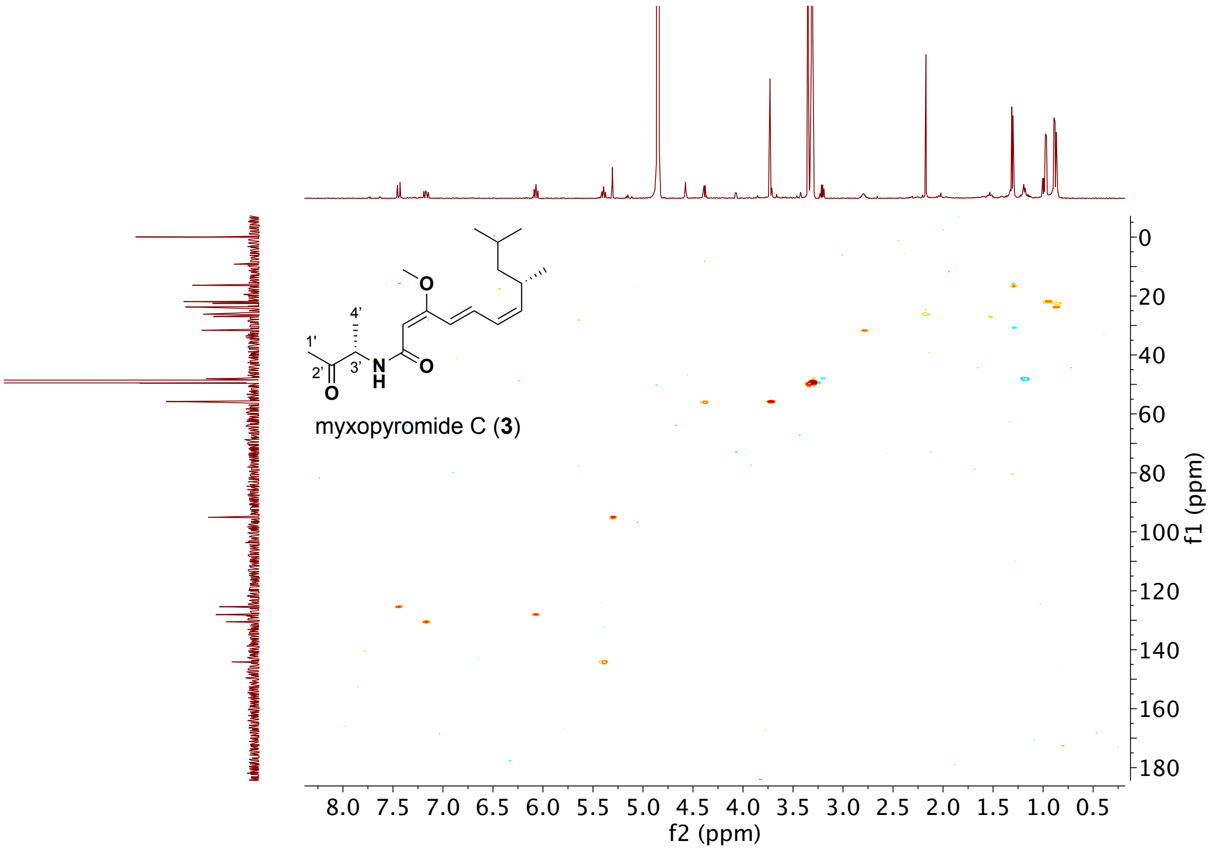

**Figure S28.** $^1$H-$^{13}$C HMBC NMR (600 MHz, $CD_3OD$) spectrum of myxopyromide C (**3**).

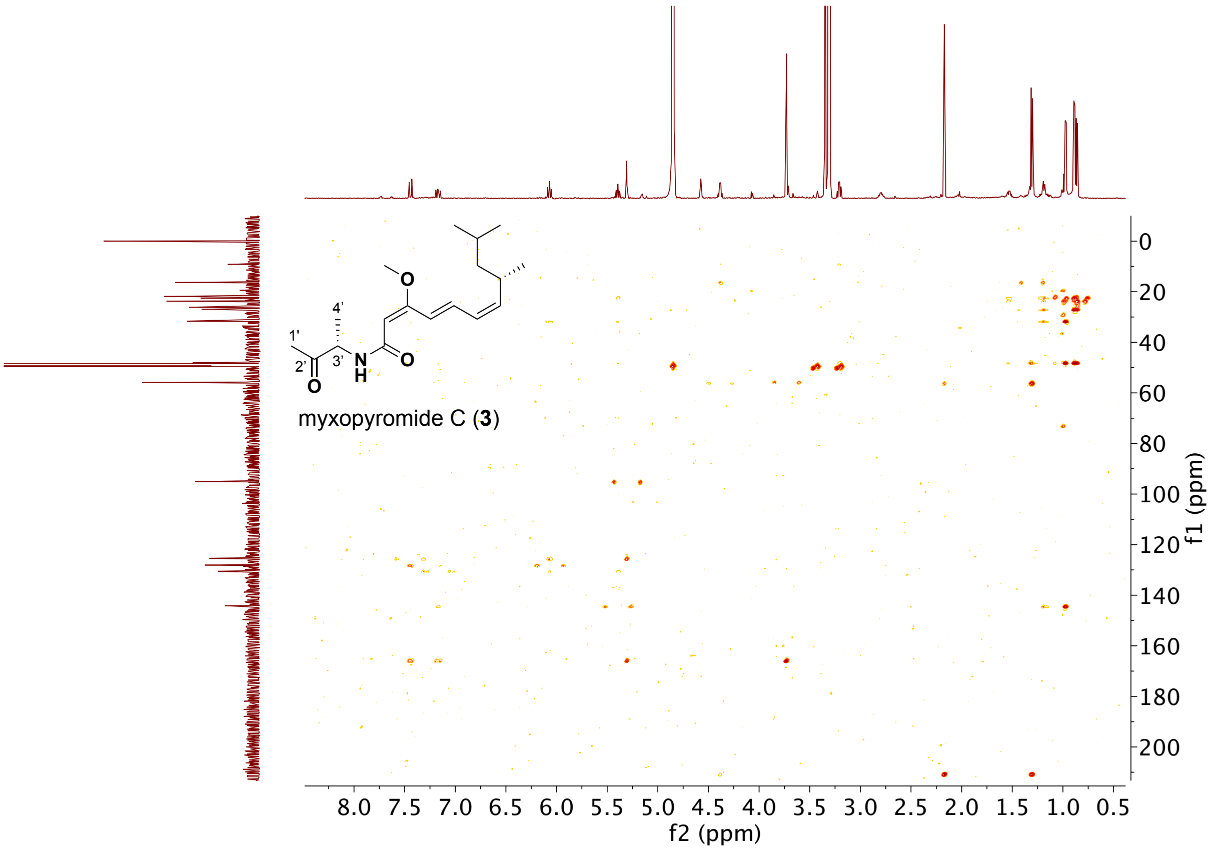

**Figure S29.** $^1$H-$^1$H COSY NMR (600 MHz, $CD_3OD$) spectrum of myxopyromide C (**3**).

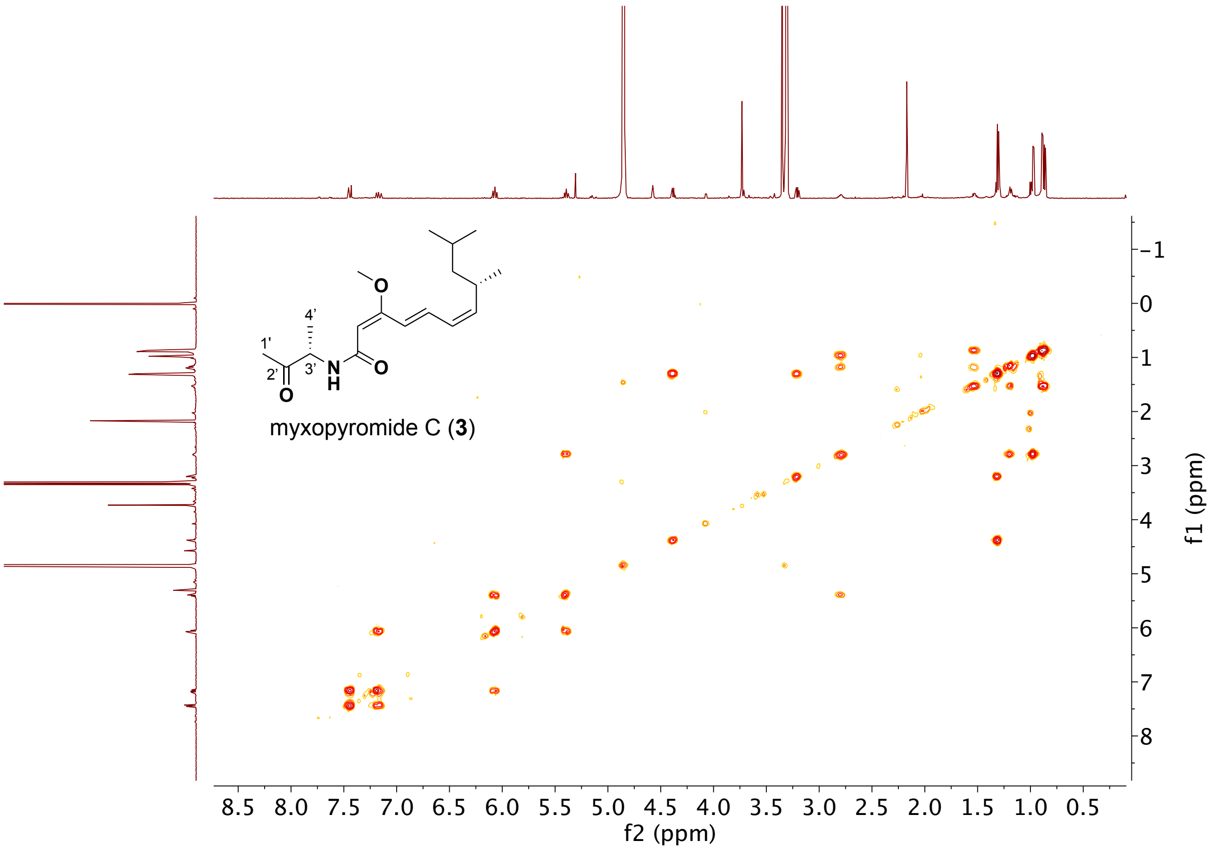

**Figure S30.** NOESY NMR (600 MHz, $CD_3OD$) spectrum of myxopyromide C (**3**).

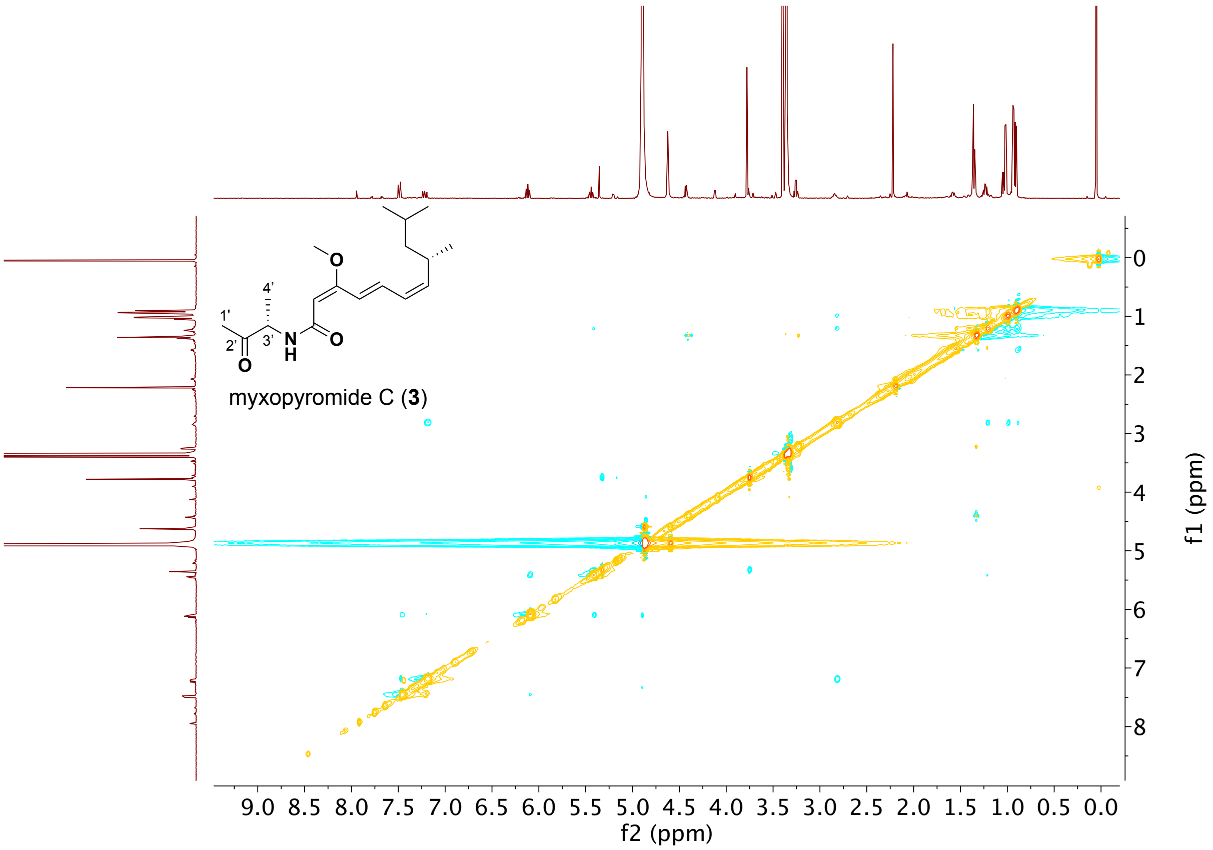

**Figure S31.** HR-ESIMS of myxopyromide C (**3**).

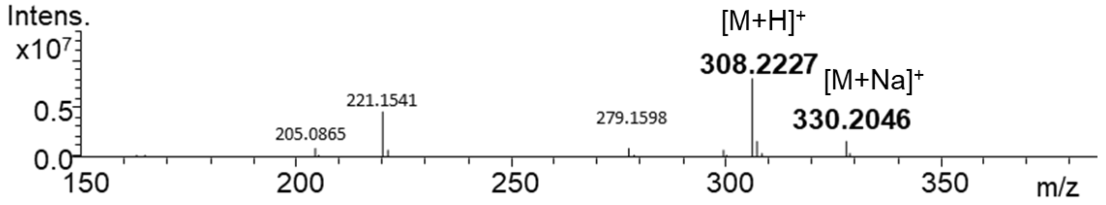


**Figure S32.** UV spectrum of myxopyromide C (**3**).

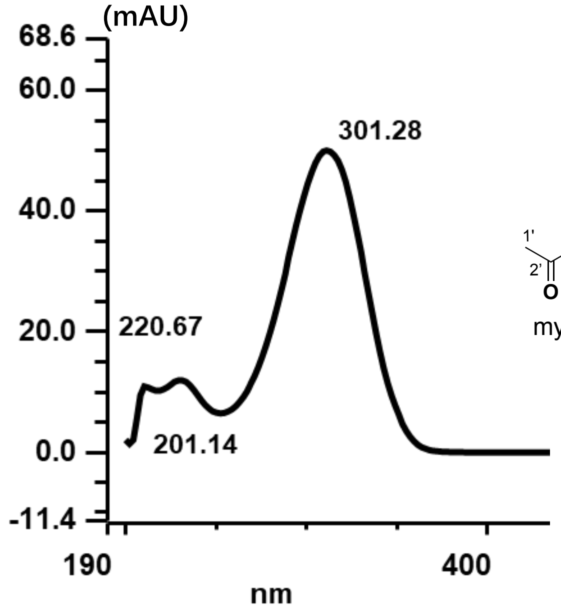


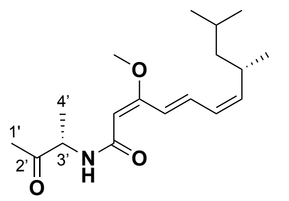


myxopyromide C (**3**)

**Figure S33.** UV spectrum of myxopyromide C (**3**).

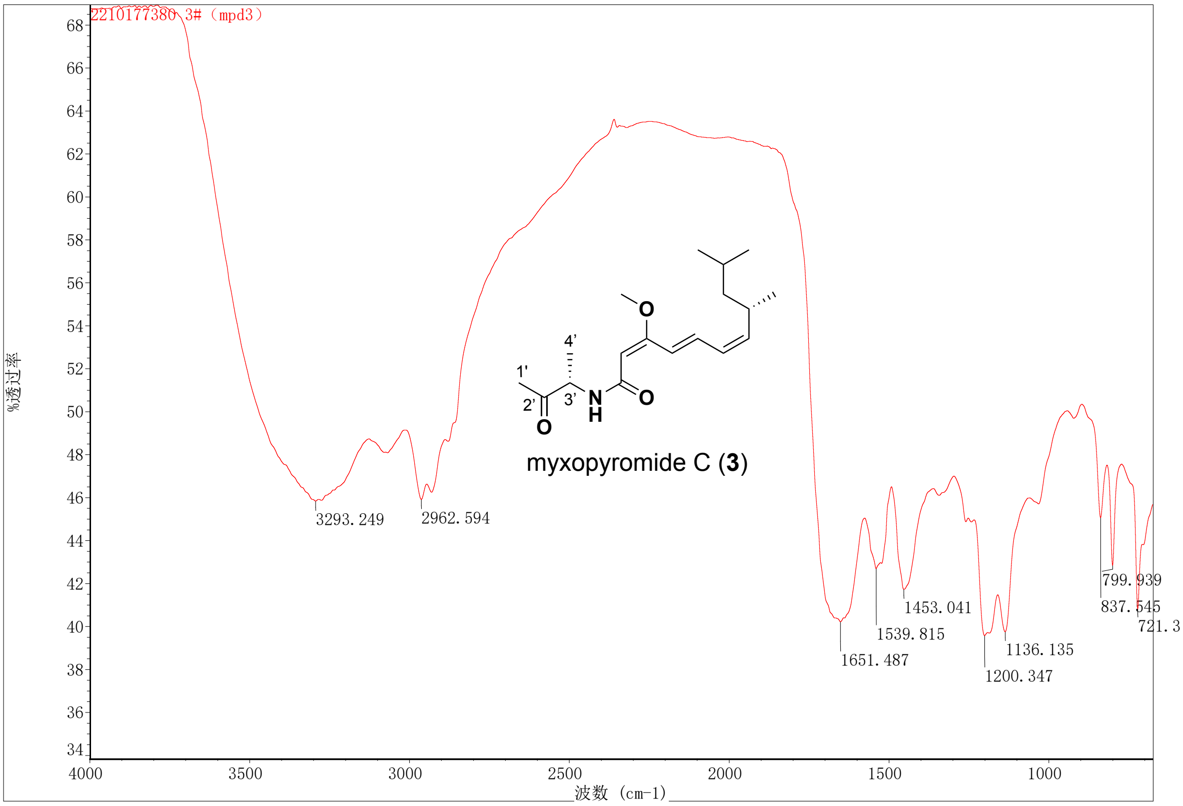

**Figure S34.** $^{1}H$ NMR (600 MHz, $CD_3OD$) spectrum of myxopyromide D (**4**).

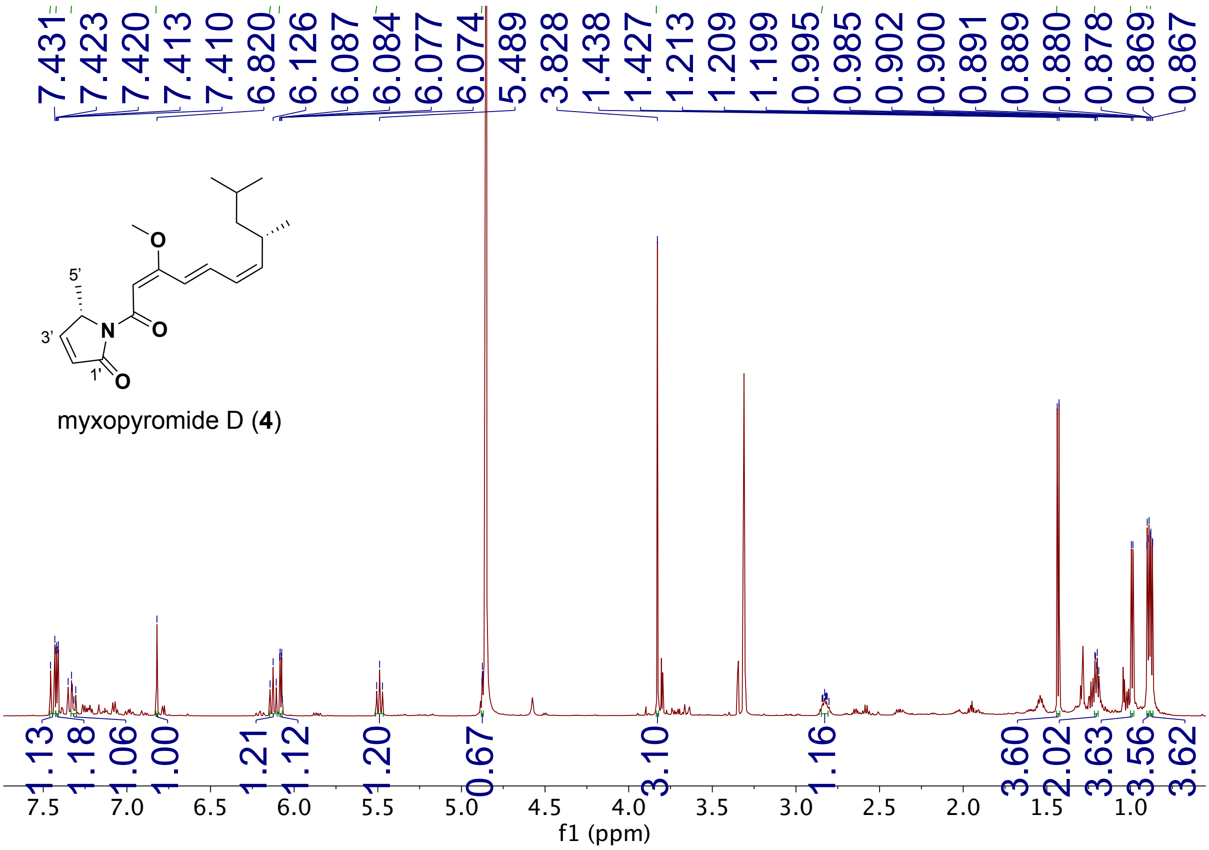

**Figure S35.** $^{13}C$ NMR (600 MHz, $CD_3OD$) spectrum of myxopyromide D (**4**).

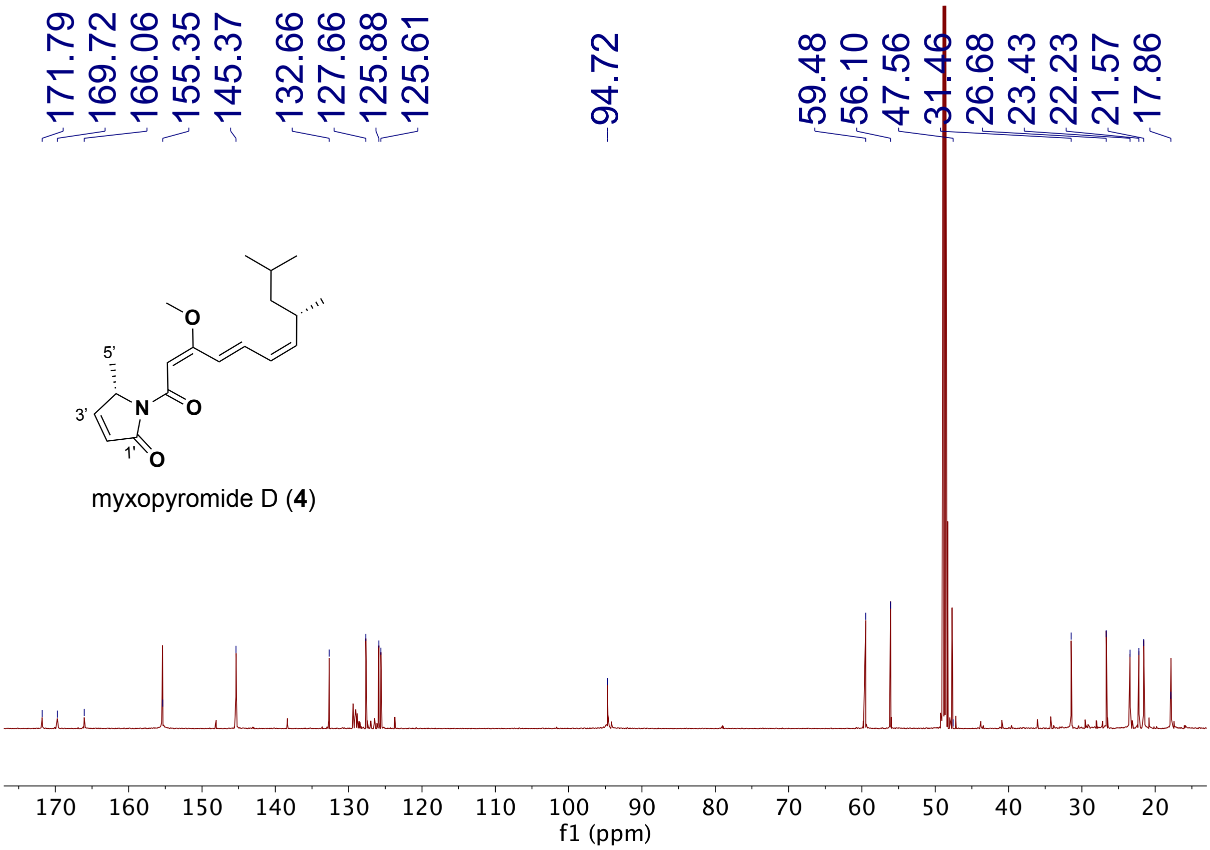

**Figure S36.** $^{1}H$-$^{13}C$ HSQC NMR (600 MHz, $CD_3OD$) spectrum of myxopyromide D (**4**).

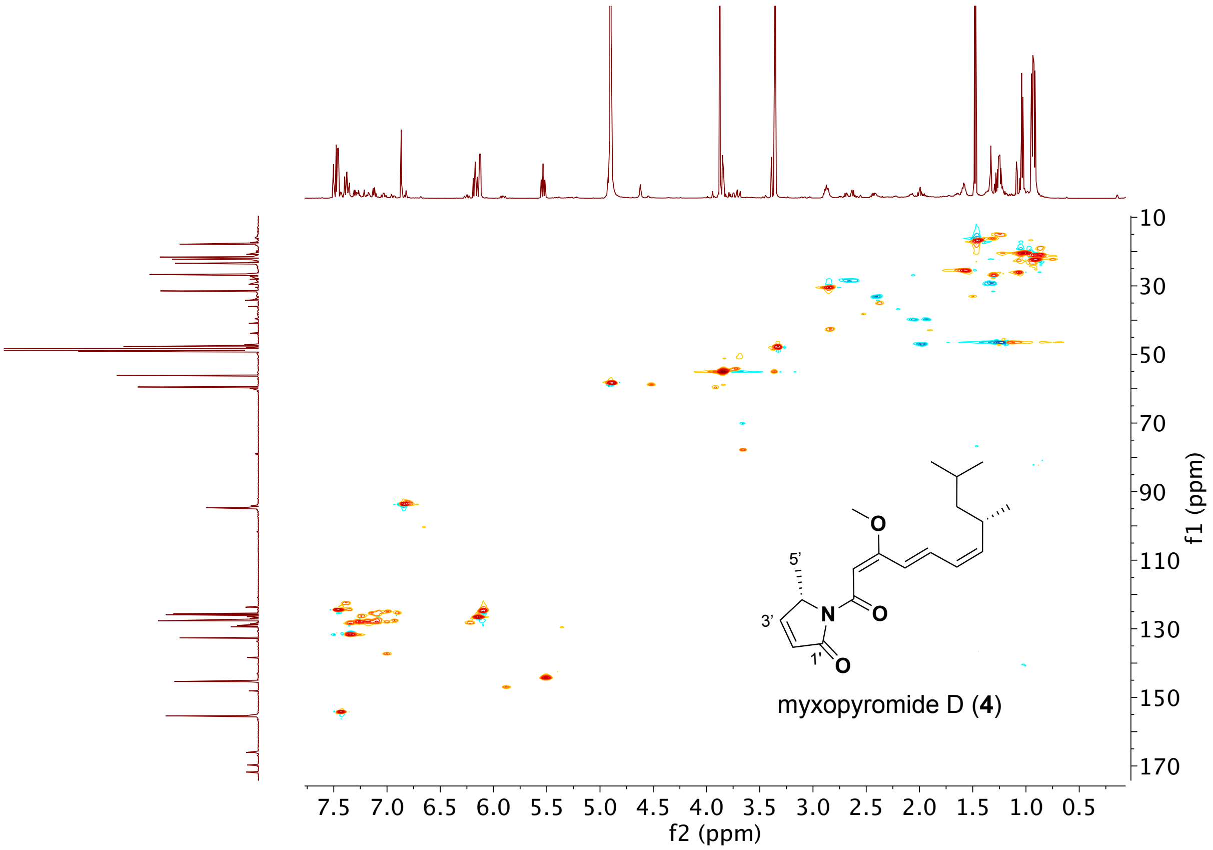

**Figure S37.** $^1H$-$^{13}C$ HMBC NMR (600 MHz, $CD_3OD$) spectrum of myxopyromide D (**4**).

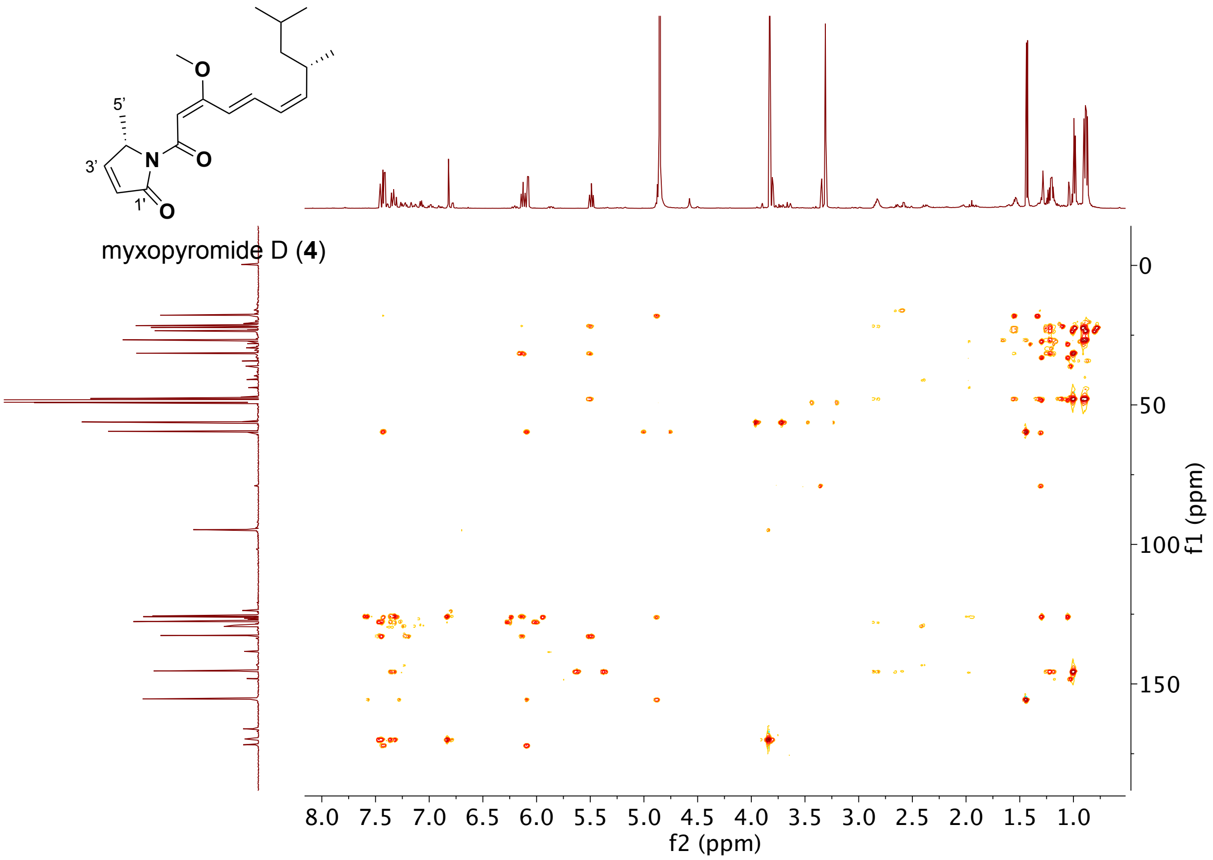

**Figure S38.** $^1$H-$^1$H COSY NMR (600 MHz, $CD_3OD$) spectrum of myxopyromide D (**4**).

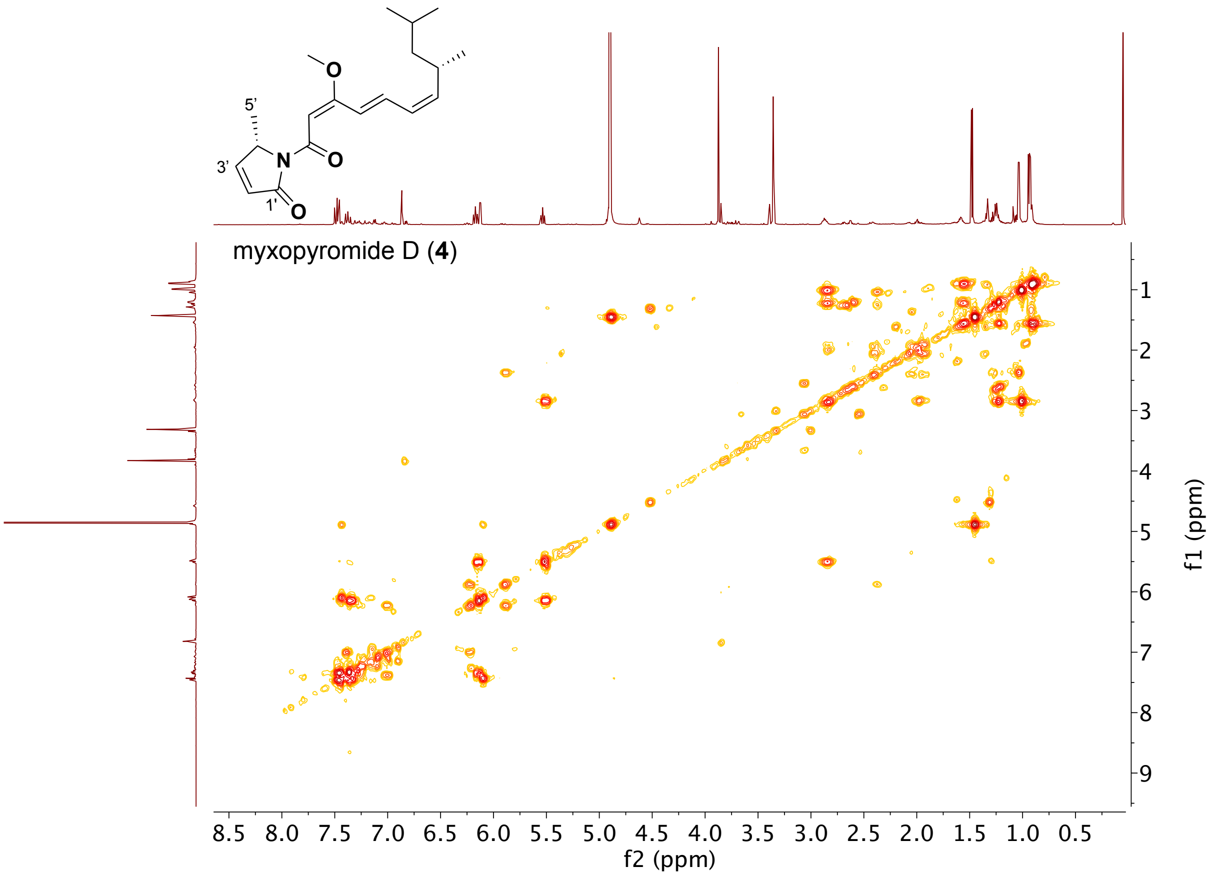

**Figure S39.** NOESY NMR (600 MHz, $CD_3OD$) spectrum of myxopyromide D (**4**).

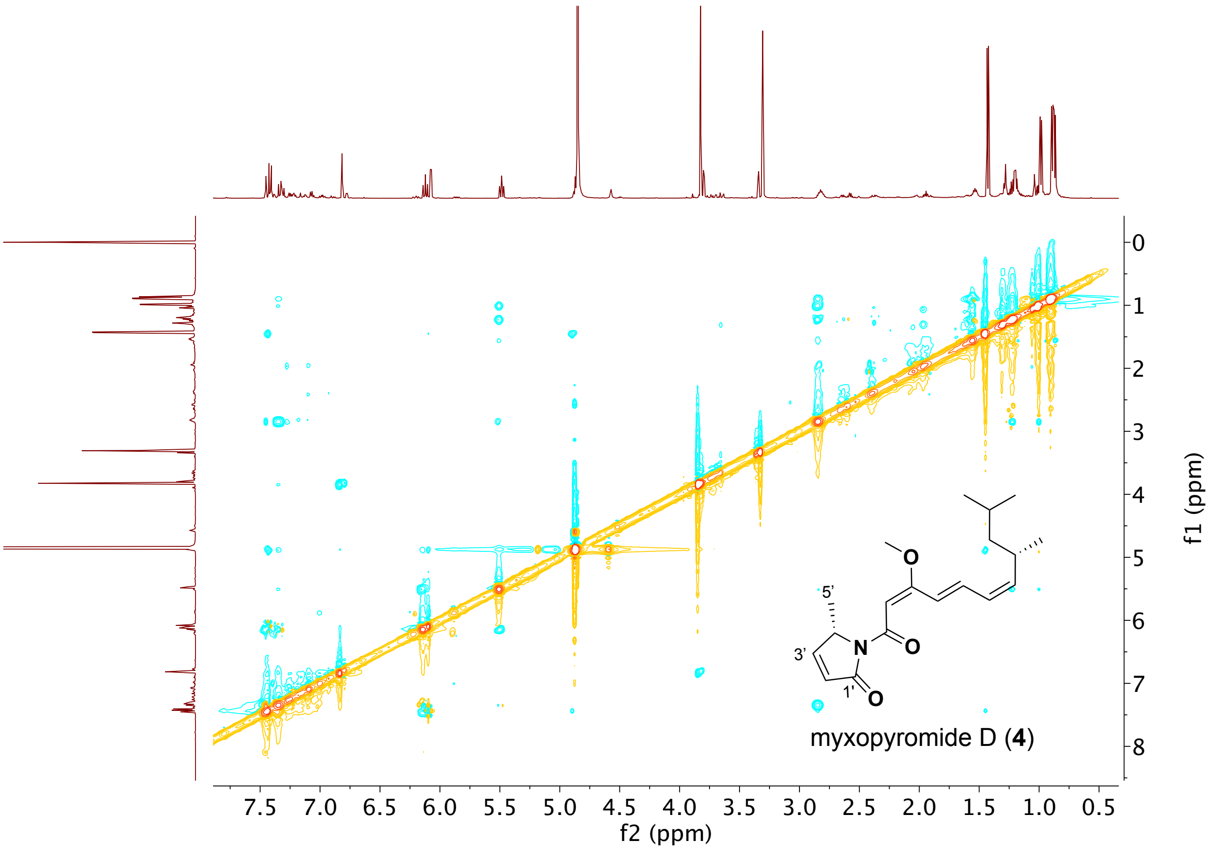

**Figure S40.** HR-ESIMS of myxopyromide D (**4**).

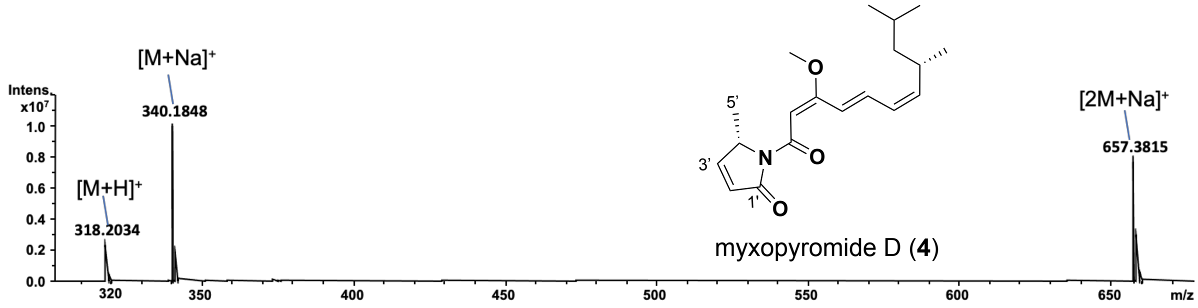


**Figure S41.** UV spectrum of myxopyromide D (**4**).

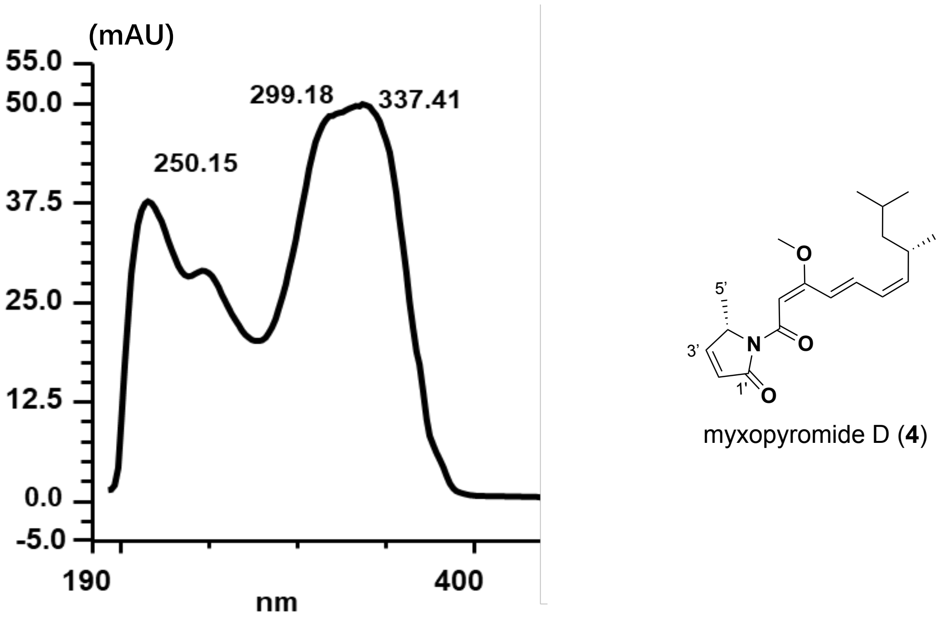

**Figure S42.** UV spectrum of myxopyromide D (**4**).

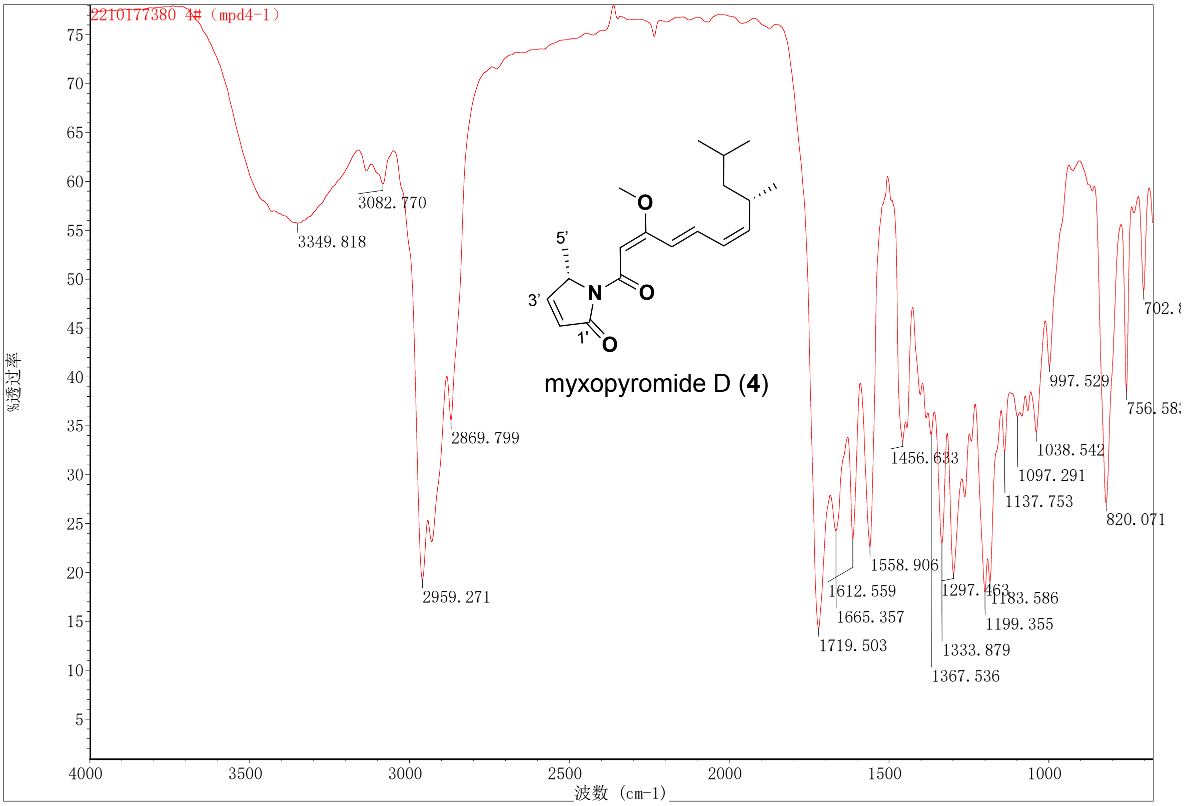

**Figure S43.** $^{1}H$ NMR (600 MHz, $CD_3OD$) spectrum of myxopyromide E (**5**).

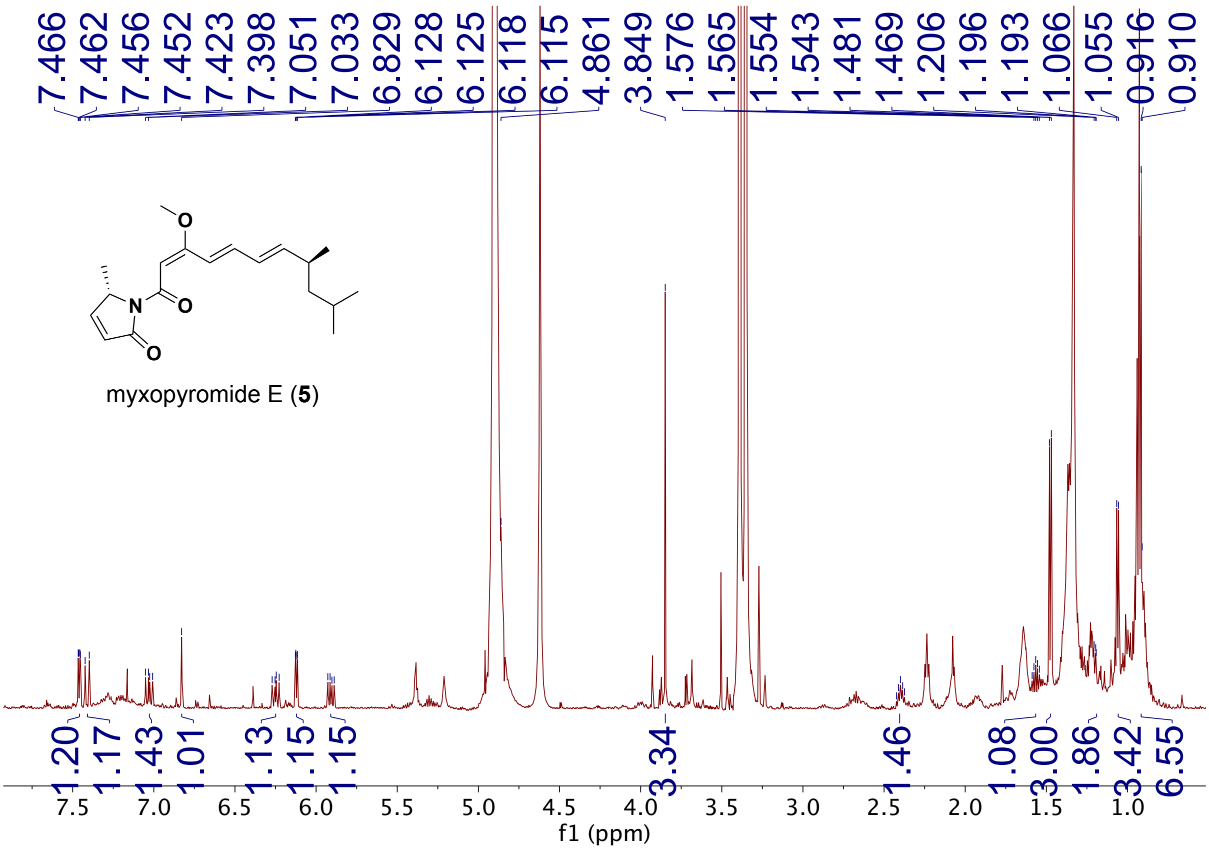

**Figure S44.** $^{13}C$ NMR (600 MHz, $CD_3OD$) spectrum of myxopyromide E (**5**).

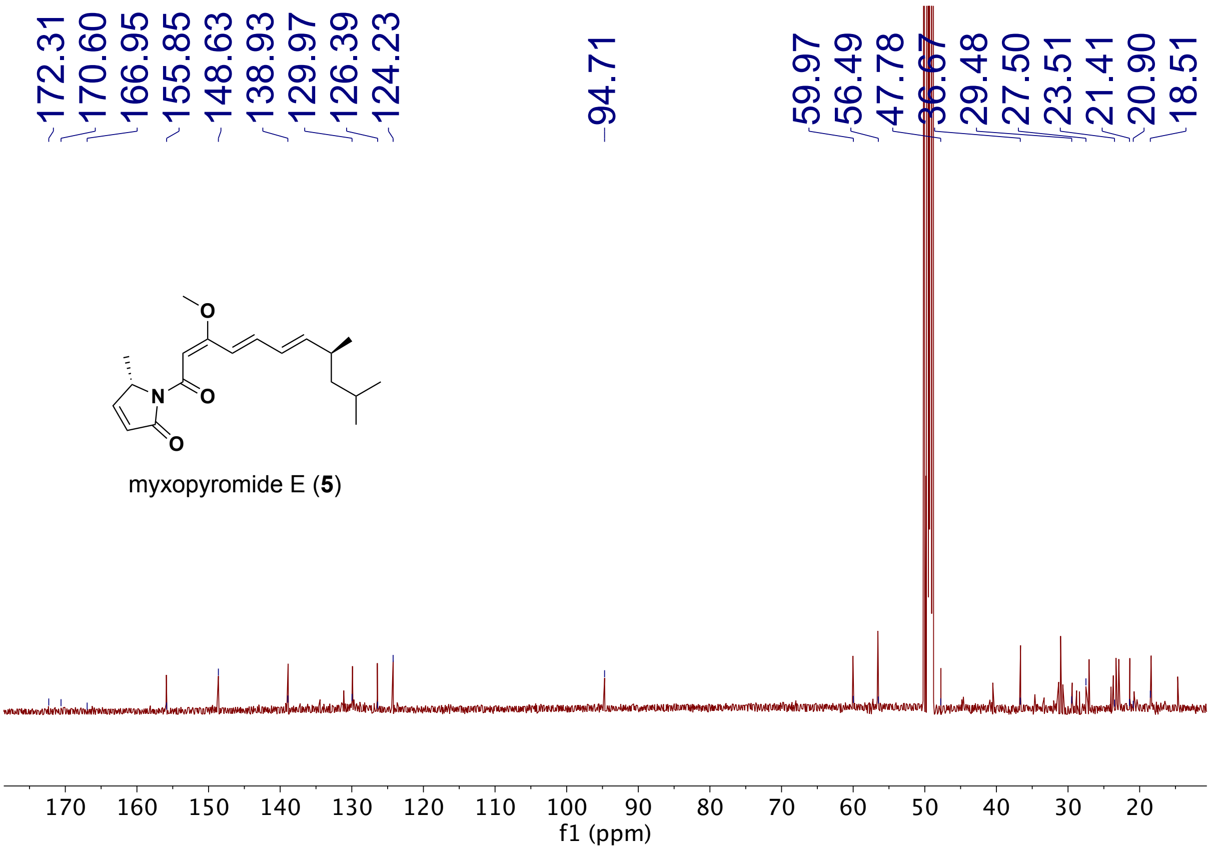

**Figure S45.** $^{1}H$-$^{13}C$ HSQC NMR (600 MHz, $CD_3OD$) spectrum of myxopyromide E (**5**).

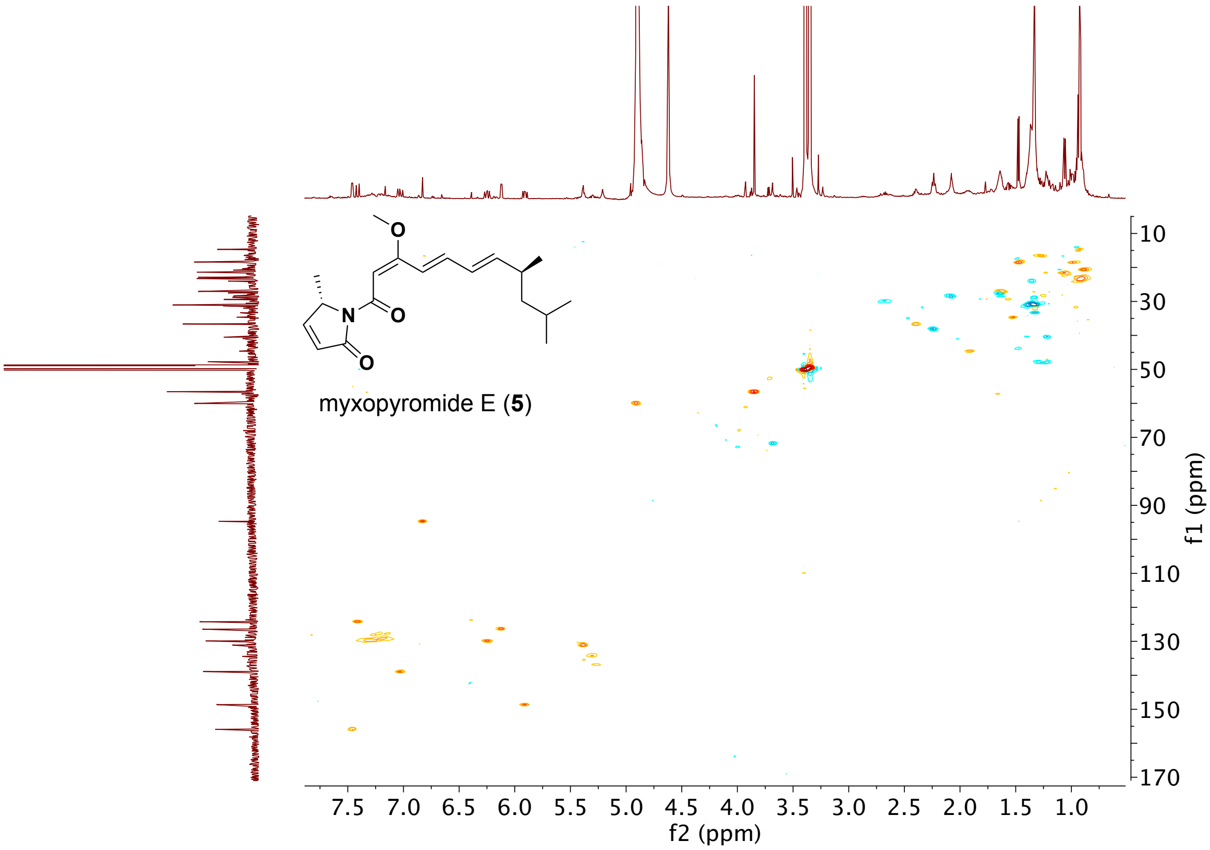

**Figure S46.** $^1H$-$^{13}C$ HMBC NMR (600 MHz, $CD_3OD$) spectrum of myxopyromide E (**5**).

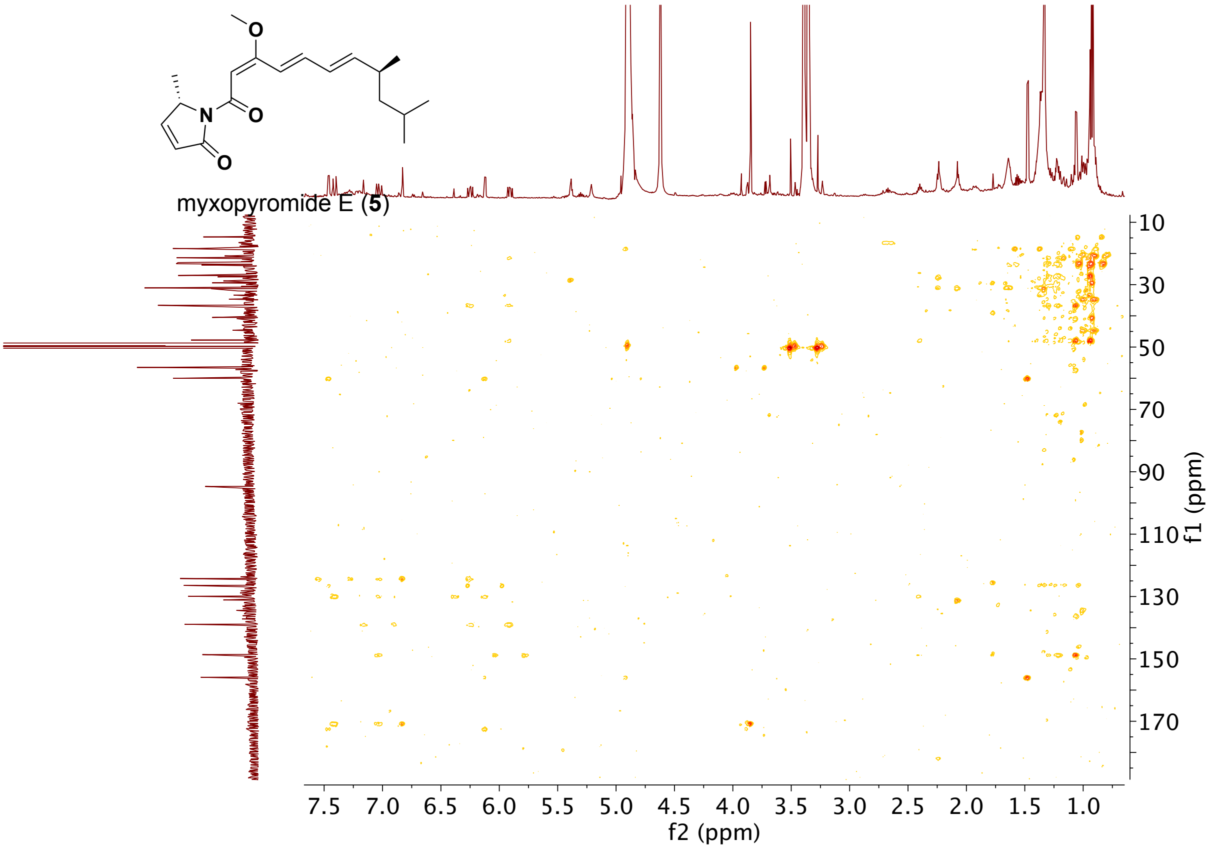

**Figure S47.** $^{1}H$-$^{1}H$ COSY NMR (600 MHz, $CD_3OD$) spectrum of myxopyromide E (**5**).

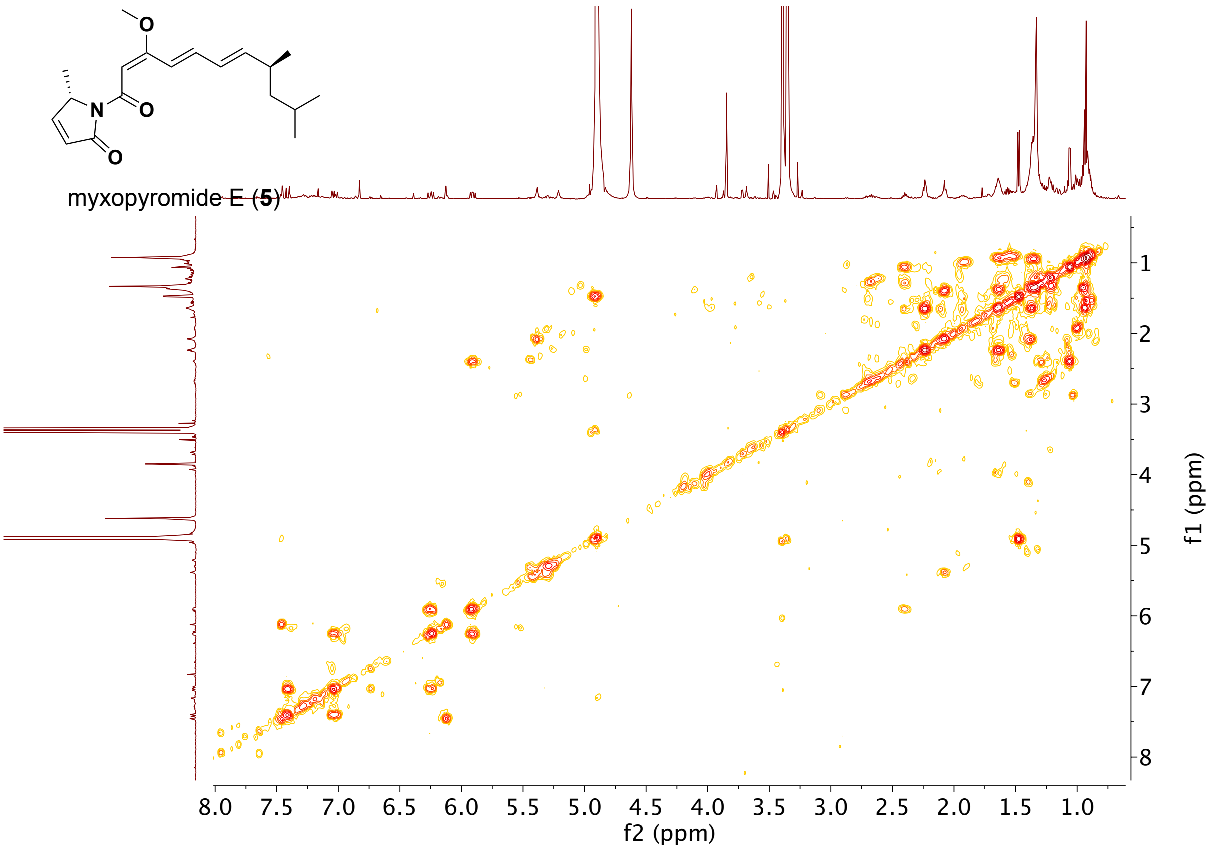

**Figure S48.** NOESY NMR (600 MHz, $CD_3OD$) spectrum of myxopyromide E (**5**).

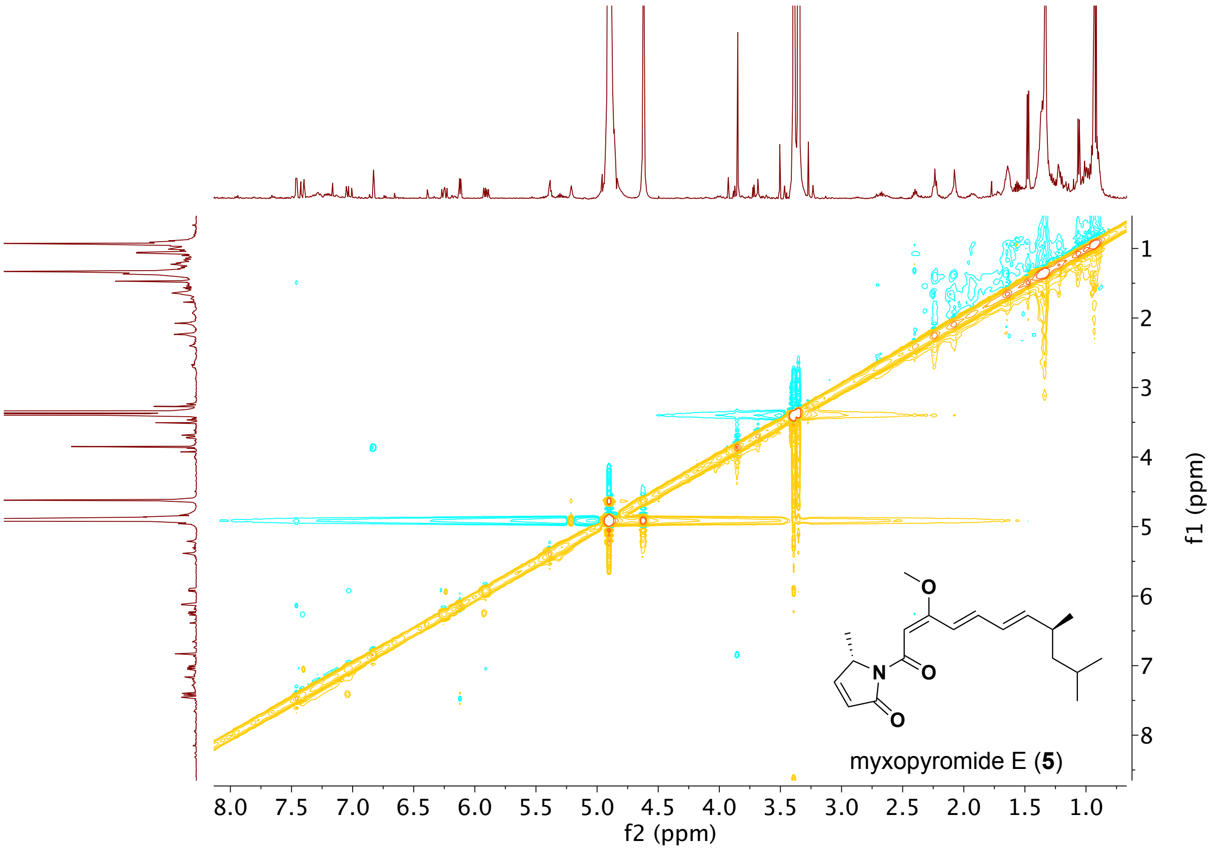

**Figure S49.** HR-ESIMS of myxopyromide E (**5**).

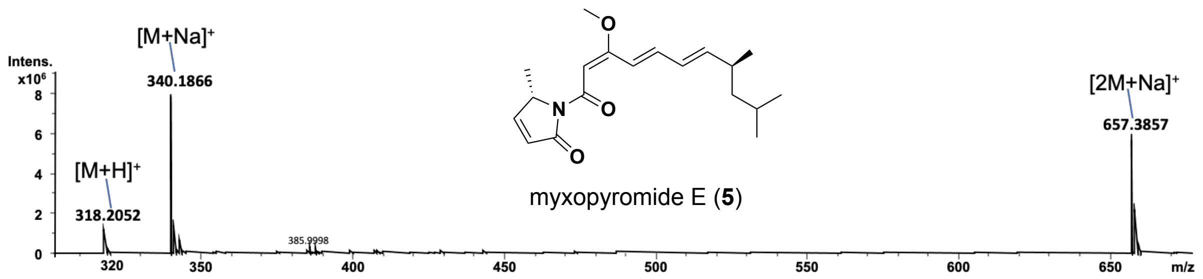


**Figure S50.** UV spectrum of myxopyromide E (**5**).

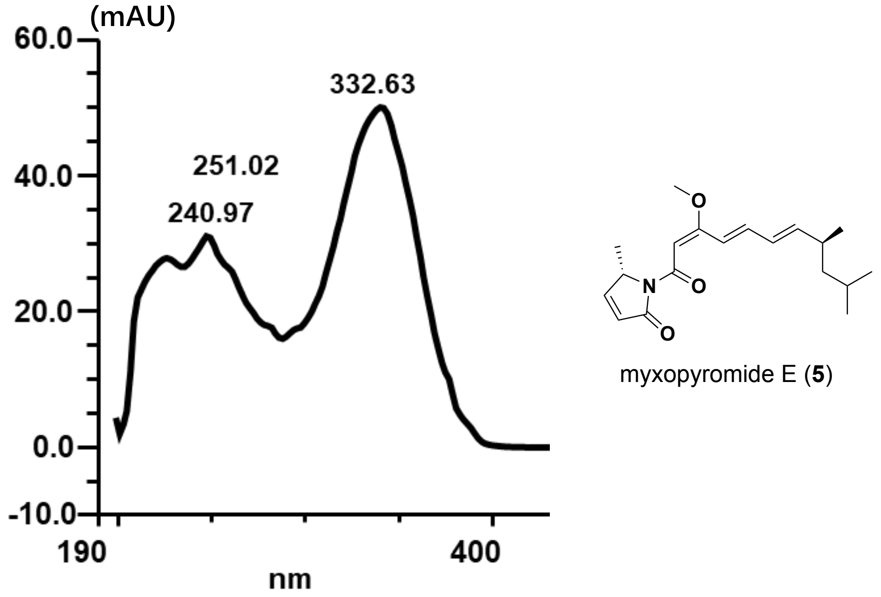

**Figure S51.** IR spectrum of myxopyromide E (**5**)

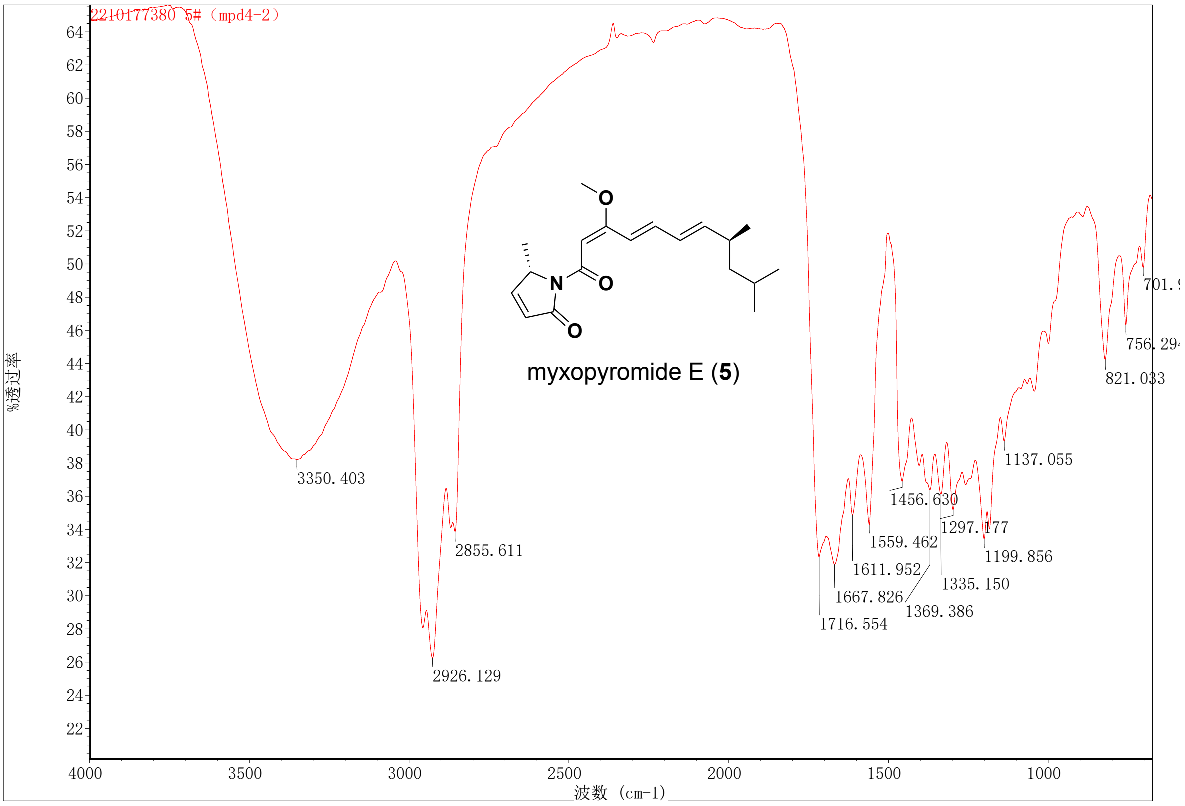
